# U5 snRNA interactions with exons ensure splicing precision

**DOI:** 10.1101/2021.03.07.434243

**Authors:** Olga V. Artemyeva-Isman, Andrew C.G. Porter

## Abstract

Imperfect conservation of human pre-mRNA splice sites is necessary to produce alternative isoforms. This flexibility is combined with precision of the message reading frame. Apart from intron-termini GU_AG and the branchpoint A, the most conserved are the exon-end guanine and +5G of the intron-start. Association between these guanines cannot be explained solely by base-pairing with U1snRNA in the early spliceosome complex. U6 succeeds U1 and pairs +5G in the pre-catalytic spliceosome, while U5 binds the exon-end. Current U5snRNA reconstructions by CryoEM cannot explain the conservation of the exon-end G. Conversely, human mutation analyses show that guanines of both exon-termini can suppress splicing mutations. Our U5 hypothesis explains the mechanism of splicing precision and the role of these conserved guanines in the pre-catalytic spliceosome. We propose: 1) Optimal binding register for human exons and U5 - the exon junction positioned at U5Loop1 C_39_|C_38_. 2) Common mechanism of base pairing of human U5snRNA with diverse exons and bacterial *Ll.*LtrB intron with new loci in retrotransposition - guided by base pair geometry. 3) U5 plays a significant role in specific exon recognition in the pre-catalytic spliceosome. Our statistical analyses show increased U5 Watson-Crick pairs with the 5’exon in the absence of +5G at the intron-start. In 5’exon positions -3 and -5 this effect is specific to U5snRNA rather than U1snRNA of the early spliceosome. Increased U5 Watson-Crick pairs with 3’exon position +1 coincide with substitutions of the conserved -3C at the intron 3’end. Based on mutation and X-ray evidence we propose that -3C pairs with U2 G_31_ juxtaposing the branchpoint and the 3’intron-end. The intron-termini pair, formed in the pre-catalytic spliceosome to be ready for transition after branching, and the early involvement of the 3’intron-end ensure that the 3’exon contacts U5 in the pre-catalytic complex. We suggest that splicing precision is safeguarded cooperatively by U5, U6 and U2snRNAs that stabilise the pre-catalytic complex by Watson-Crick base pairing. In addition, our new U5 model explains the splicing effect of exon-start +1G mutations: U5 Watson-Crick pairs with exon +2C/+3G strongly promote exon inclusion. We discuss potential applications for snRNA-therapeutics and gene repair by reverse splicing.

## 1. Introduction

Human genes can generate multiple protein isoforms by alternative splicing (AS) of different sets of pre-mRNA exons, which enables another layer of regulatory control over gene function in development and adaptive processes. AS is involved in regulation of cell fate from the earliest switch of pluripotent embryonic stem cells to specific lineages (Gabut et al., 2011; Fiszbein and Kornblihtt, 2017; Su et al., 2018) until terminal differentiation of somatic stem cells in adults (Nakka et al., 2018). AS controls proliferation and apoptosis of specialised cells such as T-cells (Corrionero et al., 2011) and response to genotoxic stress (Shkreta and Chabot, 2015; Shkreta et al., 2011; Munoz et al., 2017).

Pre-mRNA splicing is catalysed by the spliceosome, a multi-molecular dynamic complex, which shares a remarkably conserved ribozyme core with ancient mobile Group II introns, found in bacteria, archaea and eukaryotic organelles. In effect, the mechanism of splicing, a 2-metal-ion-ribozyme catalysis (Steitz and Steitz, 1993; Fica et al., 2013) much predates the origin of eukaryotes and is thought to have been driving molecular evolution in the primordial RNA world (Gilbert, 1986; Koonin, 2006; Check, 2012; Irimia and Roy, 2014).

The modern spliceosome combines the flexibility essential for the aforementioned complex gene regulation in metazoans with the routine precision of the RNA message to preserve the reading frame for the effective protein translation. The RNA components of the spliceosome, small nuclear U-RNAs (U1, U2, U6 and U5) pair short sequences in the pre-mRNA, which are imperfectly conserved to allow for alternative sites to be used. This choice of splice sites is often regulated by RNA binding proteins, RBP, and can be overruled by mutations that increase splice site complementarity to snRNAs (Hamid and Makeyev, 2017). While weak splice site conservation is clearly required to produce alternative isoforms, the exact mechanism that guaranties splicing precision in spite of these sequence variations is still unknown and is the focus of this study.

In human pre-mRNA introns, apart from the AG_GU di-nucleotides of the intron termini the most conserved bases are the branch point adenine, the exon-end guanine (-1G), and the +5G near the start of the intron (**Figure 1A**, Sheth et al., 2006; Mercer et al., 2015). The relationship between these conserved guanines has been scrutinised for over twenty years and linked to the initial recognition of the exon/intron boundary by U1 snRNA (**Figure 1B**). During the development of the GENESCAN algorithm for exon/intron gene structure prediction Burge and Karlin (1997) examined statistically the dependencies between the nucleotides at the exon/intron boundary. The authors reported a “*compensation effect”*: that in the absence of the intronic +5G the exon-end G (-1G) is almost invariant. Comparative analysis of substitutions in human and mouse orthologous 5’ splice sites also showed the same dependency between the exon-end guanine and +5G at the start of the intron (Carmel et al., 2004). A recent study (Wong et al., 2018) employed a focused massively parallel splicing assay (MPSA) to empirically examine the effects of all possible variants of the 9nt sequence NNN/GYNNNN of the exon/intron boundary on exon inclusion (percent spliced-in, PSI). This approach allowed to quantify the relationship of these conserved guanines by measuring PSI and the authors conclude, that previously observed *“seesaw linkage”* pattern, whereby exon-end G (-1G) permits any nucleotide at intron position +5 and vice versa +5G allows any nucleotide at the end of the exon, is in fact “*a strong positive interaction between -1G and +5G”*, such that a substitution at either of these conserved positions results in over 20% reduction of PSI.

**Figure 1.**
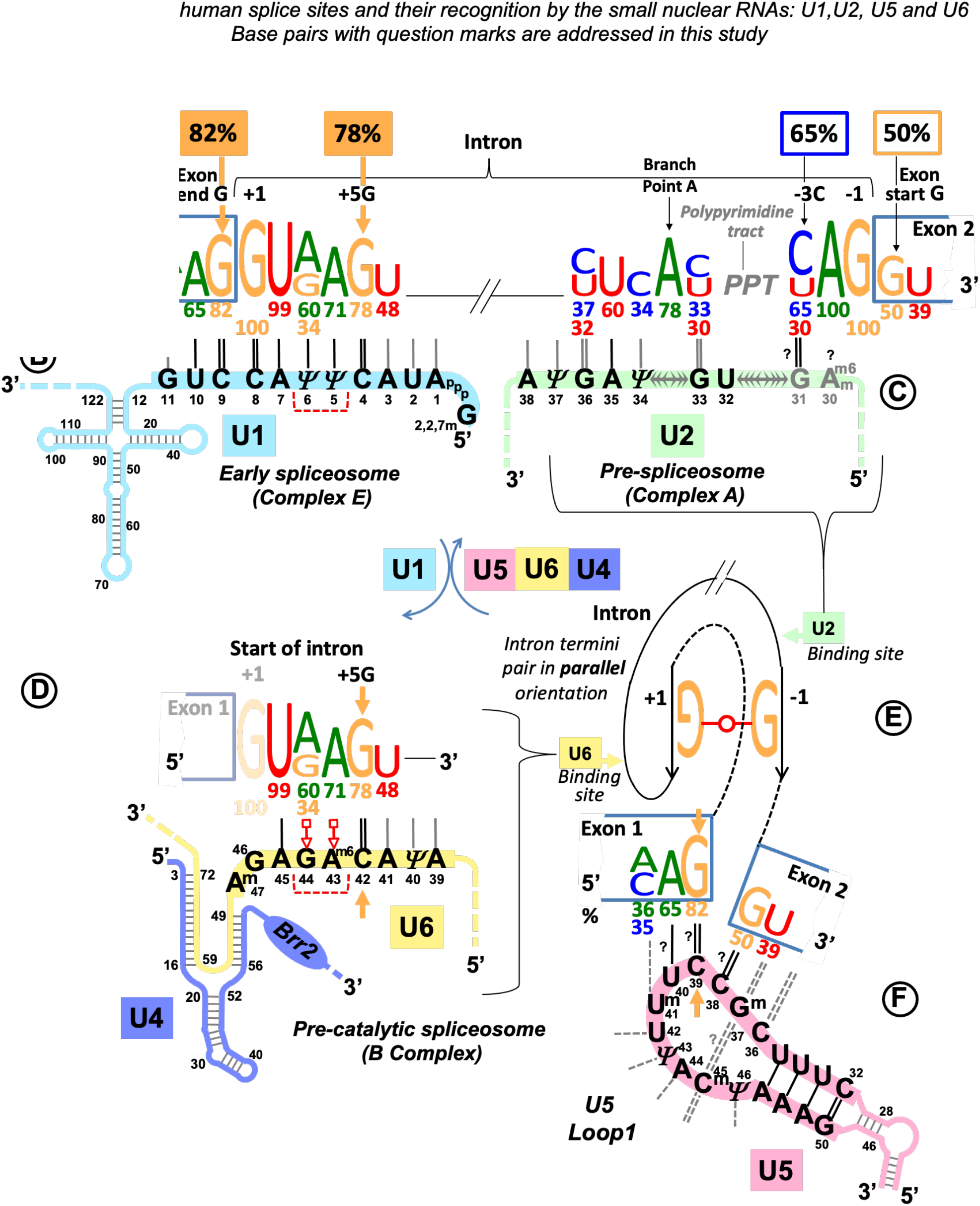
Multistage recognition of variable human splice sites by U snRNAs. **A:** Only 7 nucleotides in human introns are conserved above 75% (Sheth et al., 2006; Mercer et al., 2015). Apart from the terminal di-nucleotides and the branch point A the most important are the two guanines of the exon-end and at position +5 at the intron-start (orange labels and arrows). **B:** In the early spliceosome U1 snRNA forms on average 7 Watson-Crick pairs with human exon/intron boundaries (Carmel et al., 2004). **C:** In the pre-spliceosome U2 snRNA forms the branchpoint helix with an adenosine bulge. (**?**) Proposed U2 G_31_=C_-3_ pair, see Discussion **3.6.1 D:** In the pre-catalytic spliceosome U1 quits the complex and the start of the intron is passed on to U6 snRNA. +5G pairs with U6 42G (orange arrow). The conserved adenines +3, +4 form non-Watson-Crick pairs with U6 44G and 43A^m6^ (Galej et al., 2016, Konarska et al. 2006; shown here in red according to Westhof geometric classification: 10^th^ family, Leontis et al., 2002; role explained in Figure 14 legend). The stable U6/start of intron helix is a checkpoint for the later spliceosome activation by *Brr2* helicase (binding site on U4: blue oval). *Brr2* unwinds U6/U4 duplexes and frees U6 to configure the catalytic site of the spliceosome (Nielsen and Staley, 2012). **E:** The strictly conserved non-canonical G••G (2^nd^ Westhof geometric family, Discussion **3.6.2, Table 1B,** Scadden and Smith, 1995; Costa et al., 2016). **F:** At the pre-catalytic stage U5 snRNA comes into the complex together with U6 as part of U5•U4/U6 tri-snRNP (Wahl et al., 2009 and 2015; Scheres and Nagai, 2017; **Table S1**). As U1 quits the complex, the 5’exon is passed on to U5 snRNA Loop1. For the 3’exon see Discussion **3.6.2**. Aligned together the exons form the splice junction consensus AG|G (proto-splice site, Sverdlov et al., 2004) pictured here paired with complementary C_38_C_39_U_40_ of the U5 Loop1. In this way the most conserved exon-end G pairs with U5 39C (orange arrow). If so, in the pre-catalytic spliceosome the intron-termini pair and U6 non-Watson-Crick pairs are stabilised by flanking U5/5’exon and U6/intron-start helices, each secured by one of the two important guanines of the human splice signals (orange arrows in **E** and **D**). **Post-transcriptional base modifications of snRNAs**: Ψ: pseudouridine; Superscript m: 2’O-methyl; A^m6^: N6-methyladenosine; A^m6^ : 2’O-methyl,N6-methyladenosine (modification positions as in Anokhina et al., 2013).

**Table 1.** Intron termini pairs conserve parallel local orientation of the RNA strands

A.** *Examples of different pairs between the first and sub-ultimate bases of Group II introns\**
| Base pairs | Group IIA | Group IIB | Group IIC | Compensatory mutations |
| --- | --- | --- | --- | --- |
| **G←→A** | *L.l.LtrB* | *ai5γ* | *O.i.l1* |  |
|  | *al1 (S.c. cox1)* | *Avi.groEL (A.v.l1)* | *A.v.l2* |  |
|  | *al2 (S.c. cox1)* | *E.c.l3, l5, l8* | *E.c.l7* |  |
|  | *O.i.l2* | *Tel3c, 4c, 4f (Th.el.)* | *Sr.me.l5* |  |
| **G←→C** |  | *Rmlnt1 (Sr.me.l1)* |  |  |
|  |  | *E.c.l2* |  |  |
| U-G |  | *Tel4h* |  | Chanfreau and Jacquier 1993 |
| C-G |  |  |  | Chanfreau and Jacquier 1993 |
| G-U |  | *Sr.me.l2* |  |  |

**B.** *Occurrences of different base pairs at the intron termini in humans\*\**
| Base pairs | All human introns | Major (U2) spliceosome | Minor (U12)*** spliceosome | Compensatory mutations |
| --- | --- | --- | --- | --- |
| **G←→G** | **99%** | **99%** | **83%** |  |
| **A←→C** | **1%** | **0.01%** | **13%** |  |
| **A←→A** | 0.01% |  | 1% | Scadden and Smith 1995 |
| A-G | 0.01% | 0.003% | 1% |  |
| A-U | 0.005% |  | 0.7% |  |
| U-G | 0.005% | 0.005% |  |  |
| G-A | 0.004% | 0.005% |  |  |
| G-U | 0.003% | 0.002% | 0.3% |  |
| G-C | 0.0004% | 0.0005% |  |  |

\* Plante and Cousineau 2006; Eskes et al. 1997, 2000; Somarowthu et al. 2014; Ferat et al. 2003; Chillón et al. 2014; Toor et al. 2008; Mohr et al., 2010 and Zymmerly Database http://webapps2.ucalgary.ca/~groupii/
\*\*Based on Parada et al. 2014 and Sheth et al. 2006
\*\*\*Minor spliceosome (U12) processes less than 0.4% of all human introns (Sheth et al. 2006)

Experimentally changing a suboptimal exon-end nucleotide to G can completely supress the effect of various splicing mutations associated with genetic disease. *IKBKAP* IVS20 (+6T → C) mutation that causes a skip of exon 20 in 99.5% of patients with familial dysautonomia (a recessive congenital neuropathy) is completely neutralised by the exon-end A → G change leading to almost 100% exon 20 inclusion (Carmel et al., 2004). *ATR* c.2101A → G mutation within exon 9 is synonymous, however it appears to strengthen the exonic splicing silencer (ESS) and results in only a trace of the correct transcript and a very severe, but not lethal, phenotype (Seckel syndrome associated with dwarfism and microcephaly). The effect of this mutant ESS can be overruled by the change of exon-end T → G, that produced almost exclusively normally spliced product (Scalet et al., 2017). Another example, coagulation factor 5 has an alternative intron within a 2.7kb exon 13, that is spliced out in a small fraction of transcripts, leading to ∼1% of Factor5-short protein isoform in plasma normally. This alternative intron is preceded by an adenine: an A → G change in this case enhances exon-end definition and leads to the predominant exclusion of this alternative intron causing a rare bleeding disorder (F5-Texas phenotype, Vincent et al., 2013).

Currently, the only mechanistic explanation of the strong dependency of exon-end G and intron +5G, as well as the ability of exon-end G to supress splicing mutations or hyperactivate splicing is centred on the 5’ splice site selection by base pairing with U1 snRNA. Indeed, U1 specifically engineered to increase complementarity to 5’ss can also partially restore exon inclusion (as in Carmel et al., 2004), a discovery of Zuang and Weiner, 1986 which led to the development of snRNA therapeutics (see Discussion). However, the functional 5’ splice site is not defined only by complementarity to U1 snRNA, although shifts and bulges in the U1 binding register at divergent exon-intron boundaries have been proposed to fix the problem of poor conservation (Roca et al., 2012; Roca et al., 2013; Tan et al. 2016). In the early spliceosomal complex (complex E), U1 snRNA binds multiple alternative or cryptic sites, and the commitment to splicing depends on both the affinity to the target and relative positions of the U1 and U2 binding sites (Eperon et al., 1993; 2000). Multiple U1 snRNAs can bind initially and the surplus of U1 is removed after U2 snRNP interacts with U1 snRNP during the transition to complex A (Hodson et al., 2012; **Table S1** for successive spliceosomal complexes). It is also long known that the 5’ splice site is not defined relative to the base pairing with U1 snRNA.

Indeed, U1 snRNAs engineered to base pair in the vicinity, rather than exactly at the exon/intron boundary can rescue inclusion of exons with splicing mutations in cell culture and in mouse models (Fernandez Alanis et al., 2012; Rogalska et al., 2016; reviewed in Singh and Singh, 2019). This variability of U1 binding cannot support precise definition of the 5’ splice site, which means that the binding register of U6 snRNA is the final determinant of the intron 5’ boundary – an explanation put forward by Hwang and Cohen in 1996. In addition, U1-independent splicing was also discovered in HeLa nuclear extracts: Crispino and Sharp (1995) show that complementarity to U6 snRNA enhances splicing if U1 is depleted. The authors confirm experimentally that U6 can form Watson-Crick pairs with the intron until position +9. More recent studies (reviewed in Fukumura and Inoue, 2009) show that at least a fraction of human introns normally relies on U1-independent splicing. Moreover, engineering increased complementarity to U1 at the exon/intron boundary disrupts the normal splicing pattern, overruling exon exclusion prompted by Fox-1 RBP (Fukumura et al., 2009). A recent evolutionary insight provided by a monocellular red alga *Cyanidioschyzon merolae*, that lacks U1 snRNA and all it’s protein co-factors (U1 snRNP) shows, that U1 is altogether dispensable for pre-mRNA splicing (Matsuzaki et al., 2004; Stark et al., 2015). Single intron genes of this exceptional eukaryote do not require alternative processing. This indicates, that U1 is needed to facilitate flexible splice site choices, rather than splicing precision, and confirms that U6 controls the 5’ intron boundary definition. Moreover, formation of the U6 helix with the start of the intron (so-called U6 ACAGAGA interaction) is considered to trigger subsequent activation of the human pre-catalytic spliceosome (complex B, Charenton et al., 2019). The difficulty is that 1 in 5 human introns lack the essential +5G that pairs with U6 42C to secure this interaction **(Figure 1D**). In *S. cerevisiae,* the first model organism of the spliceosome studies, U_+4_G_+5_U_+6_ are absolutely conserved and all form Watson-Crick pairs with U6 snRNA. Upstream position +3 forms a non-Watson-Crick pair essential for the correct repositioning of the lariat intermediate after branching (Konarska et al., 2006). In humans, conservation of +3A is less reliable, so adenine is repeated in position +4, which ensures the presence of at least one of these key purine pairs. This, however, takes out a Watson-Crick pair and given that the conservation of +6U is below 50% preservation of this checkpoint U6 helix is altogether elusive, suggesting the need for other specific interactions in the pre-catalytic spliceosome. The 3’ intron-end motifs (**Figure 1A**, right) are initially recognised by proteins: SF1 binds the branchpoint (BP), the large subunit of the U2 snRNP auxiliary factor protein, U2AF^65^ tethers the polypyrimidine tracts (PPT), while the small subunit U2AF^35^ binds the intron-end AG. Only then U2 snRNA can pair with its target sequence around the branchpoint (**Figure 1C**, Complex A). This is quite unlike the usual way, when RNA guides a protein enzyme to the RNA or DNA target. Mutations at the 3’ intron end can be also suppressed by increasing complementarity of U2 snRNA to the sequence around the branchpoint (Zuang and Weiner, 1989). Polypyrimidine tract (PPT) between the branchpoint and the 3’ intron end is highly variable in length and nucleotide composition. Crystal structures of U2AF^65^ bound to poly-U indicate a sharp kink of the RNA strand (Sickmier et al., 2006) and, moreover, conversion of uridines to pseudouridines, which confers rigidity to the RNA backbone, blocks U2AF^65^ binding (Chen et al., 2010). Oddly, no RNA partner has been identified so far for the conserved position -3 at the end of the intron, although mutations in this position impair or block splicing completely. The effect of -3C → G change was explored in *Fas/CD95* intron 5: while U2AF^65^ binding was not affected, this mutation blocked U2 snRNA binding (Corrionero et al., 2011; see Discussion **3.6.1**).

Next, adenine at intron position -2 is absolutely conserved (**Figure 1A**). According to the Cryo-EM studies, -2A interacts with the branchpoint A (two H-bonds between Hoogsteen edges of adenines, glycosidic bonds in *trans* orientation), which helps to position the 3’ exon for the ligation (reviewed in Wilkinson et al., 2020). Intron termini guanines form a pair, that conserves local parallel strand orientation as in Group II introns and in eukaryotes can be substituted exclusively by A_C intron ends (pair configuration discussed in section **3.6.2**). As only 7 nucleotides of the human splice sites are conserved above 75%, the fact that half of all human exons start with a guanine is significant. Fu et al., 2011 reported that G → T mutations at the start of *GH1* exon 3, *FECH* exon 9 and *EYA1* exon 10 cause exon skipping, while G → T change at the start of *LPL* exon 5 and G → A change at the start of *HEXA* exon 13 do not affect exon inclusion. The authors explain it by the shorter PPT stretch that precedes mutations affecting splicing. However partial exon inclusion for the neutral substitutions of the exon-start G persists even if the PPT stretch is reduced to 2-5 nucleotides. Remarkably, both these neutral changes are followed by cytosines in exon position +2. Fu and colleagues continued to quantify variable splicing effect of further nine exon +1G mutations in different human genes. Here we re-examine their data and link the exon positions +2 and +3 with the inclusion efficiency.

As in the case of the exon-end guanine, experimentally changing a suboptimal exon-start nucleotide to guanine can supress *ATR* c.2101A → G mutation of the exon 9 ESS and restores exon inclusion to the wt level (Scalet et al., 2017). Unlike for the exon-end guanine, the mechanistic basis for the splicing re-activation by the exon-start G (+1G) or the variable effect of +1G mutations on splicing cannot be explained by the initial U1 selection, which points at a later stage interactions of exon sequences at splice junctions with U5 snRNA Loop1. Base-pairing of exons with U5 proved to be the most challenging of all pre-mRNA interactions with snRNAs, possibly due to the fact that as opposed to U6 binding site at the start of the intron and the sequence around the branchpoint, the exon sequences at splice junctions are less conserved in *S. cerevisiae,* than they are in humans. So, even the binding register of the exons with U5 Loop1 presented a problem. While for the intron interactions with U6 and U2 easy alignment facilitated compensatory double mutation analyses (Zuang and Weiner, 1989, Hwang and Cohen in 1996, Crispino and Sharp, 1995) mutation analysis of U5 Loop1 was jumbled up by the absence of the interaction model, although mutant U5 variants promoted activation of new splice sites (Cortes et al., 1993). Crosslinking experiments of the 1990s involved 4-thiouridine (4sU) substitutions of the conserved guanines of the exon termini and could not show the wild type base pairing with U5 Loop1 (Newman et al.,1995; Sontheimer and Steitz, 1993; schematics in supplementary **Table S2)**. Both 5’ exon and 3’ exon 4sUs crosslinked to two positions of the loop: 5’ to U_40_ and U_41_ and 3’ to C_39_ and U_40_. Since the start of CryoEM structural studies of the *S.c.* and human spliceosomes, the pains to place the exons relative to U5 Loop1 produced no less than five different binding registers (Rauhut et al., 2016; Galej et al., 2016; Wan et al., 2016; Yan et al., 2016; Bertram et al., 2017^a,b^; Zhang et al., 2017; schematics in supplementary **Table S2**). Initially Rauhut et al., 2016 modelled 11nt U5 Loop1 with the exon-end G unpaired and the U_+2_ interactions as in the crosslinking experiments. Galej et al., 2016 presented 7nt U5 Loop1 with the exon-end G also unpaired and A_+2_A_+3_A_+4_ forming Watson-Crick pairs with U5 U_97_U_98_U_99_ (human U^m^_41_U_42_Ψ_43_). Wan et al., 2016 were first to present exon-end G paired to U5 C_95_ (human C_39_) and A_+2_A_+3_ paired with U_97_U_98_ (human U^m^_41_U_42_) of the 7nt U5 Loop1. Yan et al., 2016 then placed G_+1_A_+2_A_+3_ pared with U_97_U_98_U_99_ (human U^m^_41_U_42_Ψ_43_). All Cryo-EM reconstruction for the human spliceosome use MINX pre-mRNA substrate, which contains a small composite adenovirus intron (**Tables S2, S3**), yet Bertram et al., 2017^a^ places the 5’ exon end G_+1_C_+2_A_+3_ paired with U^m^_41_U_42_Ψ_43_ and Bertram et al., 2017^b^ with U_40_U^m^_41_U_42_, with U5 Loop1 open to 11nt. The other structures by Zhang et al., 2017, Zhan et al., 2018 also place this 5’ exon end G_+1_C_+2_A_+3_A_+4_ paired with U5 U_40_U^m^_41_U_42_Ψ_43_ but with the small 7nt version of U5 Loop1. The binding register for the 5’ exon with the exon-end G paired to U5 U_40_ of the 7nt U5 Loop1 currently prevails (**Figure 12B** in Discussion), as it is featured in the most recent structures with the best resolution (Zhang et al, 2019). On the contrary, base pairing for the 3’ exon is still unresolved. The root of the problem is timing of this interaction and the mechanistic challenges of bringing the 3’ exon into the catalytic core with the variable PPT stretch between the branchpoint and the 3’ss. The timing is not a problem for the 5’exon, as when U1 quits the complex at the pre-catalytic stage (complex B), the start if the intron is passed on to U6 snRNA and the end of the 5’ exon binds U5 snRNA Loop1. We consider the key role of the intron termini pair in the mechanism of splicing catalysis (see Discussion **3.6.2**) to adjust the timing for the 3’ exon interaction with U5.

These varying Cryo-EM reconstructions of U5 base pairs with the 5’ exon inclusive of the latest version and the lack of clear base pairing for the 3’ exon with the remaining part of the loop do not seem to connect to genetic studies reviewed above. The latest binding register for the 5’ exon does not include a G=C pair for the 82% conserved exon-end guanine and the 7nt loop is so small that it does not allow much base pairing for two exons (**Figure 12B** in Discussion). The structures endorse that U5 Loop1 plays little role in recognition of the exon sequences at splice junctions and thus cannot contribute to splicing fidelity. However, poor conservation of every nucleotide in human splice site sequences must be accounted for with a specific interaction, which all combined have to ensure splicing precision. We ask a question if this can be managed by U1, U2 and U6 without a substantial contribution from U5.

We start with a different approach to U5 modelling and first compare splicing with retrotransposition of a mobile bacterial Group IIA intron. Small nuclear RNAs U2, U6 and U5, that assemble on the pre-mRNA in the spliceosome core, are homologous to Group II RNA domains (Zimmerly and Semper, 2015; Galej et al., 2018; detailed Discussion **3.2**). In particular, the U5 Loop1 homologue, Id3 Loop of Domain I, controls the specificity of the Group IIA intron splicing by Watson-Crick base pairing with the exons. In retrotransposition, mobile Group IIA introns invade new loci by splicing in reverse into genomic targets ‘similar’ to their exons. The ‘similarity’ of retrotransposition sites is so far not clearly defined, but in effect, the unique DId3 Loop pairs variable target sites just like the universal U5 snRNA Loop1 fits all the diverse exon junctions in the human genome.

Here we compare the alignments of human splice junctions with U5 Loop1 to the alignments of bacterial retrotransposition sites with Group IIA DId3 loop. We propose a common mechanism of base pairing for human U5snRNA with diverse exons and bacterial *Ll.*LtrB intron with new loci in retrotransposition: recognition guided by base pair geometry. Statistical analysis of U5 interactions with human exons lends support to our alignment model with the optimal binding register for the splice junction of exons positioned at U5 Loop1 C_39_|C_38_. We find that U5 Watson-Crick pairs with the exons show a clear pattern of compensation for substitutions of the conserved nucleotides in human introns, indicating a collective mechanism whereby U5, U6 and U2 recognise their variable binding sites. We suggest that snRNAs in the pre-catalytic spliceosome together ensure fidelity before the committed ribozyme core is configured. In addition, we clearly explain the effect of human mutations on splicing (Fu et al., 2011) by base-pairing of the 3’ exon with U5 Loop1.

## 2 Results

### 2.1 Modelling U5 Loop1 base pairing with human exons on Group IIA intron interactions with retrotransposition sites

We considered that the types of pairs acceptable in the interactions of Group IIA introns with variable target sites might provide a clue to the way human exons pair with U5 snRNA. A pilot investigation of a small number of published sequences of *L.l.*LtrB retrotransposition sites and splice junctions of just one human gene, albeit a giant *dystrophin* was performed. Detailed examination and sequence alignments of these small datasets provided a pilot hypothesis and guided the design of a series of statistical tests on a large number of human splice junctions and intron sequences.

#### Base pair types in the interactions of *Ll.*LtrB with retrotransposition sites

We chose *Ll.*LtrB, a well-studied mobile Group IIA intron from *Lactococcus lactis* (Ichiyanagi et al., 2002; Coros et al., 2005; Novikova et al., 2014; Dong et al., 2020; Ll.I1 in the Zymmerly Lab Group II intron database http://webapps2.ucalgary.ca/~groupii/ Candales et al., 2012). This 2.5kb intron of the *ltrB* gene (encoding a relaxase found in the conjugative elements) folds into a typical structure of the Group IIA ribozyme: RNA domains DI to DVI (Dong et al., 2020). The intron catalyzes its own splicing, and the excised intron lariat can undergo specific reverse splicing to insert into the ‘homing’ site of the intron-less allele (retrohoming) or invade a new genomic locus choosing a ‘similar’ target sequence (retrotransposition). *Ll*.LtrB Id3 Loop is uracil-rich like U5 snRNA Loop1 (5 out of 11 nucleotides are uracils **Figure 2A,B**). Seven nucleotides of the loop bind the end of the 5’exon and 4 nucleotides bind the start of the 3’ exon (**Figure 2A**). This is a typical pattern of exon binding by the intron ribozyme of the subclass IIA (supplementary **Figure S1**).

**Figure 2.**
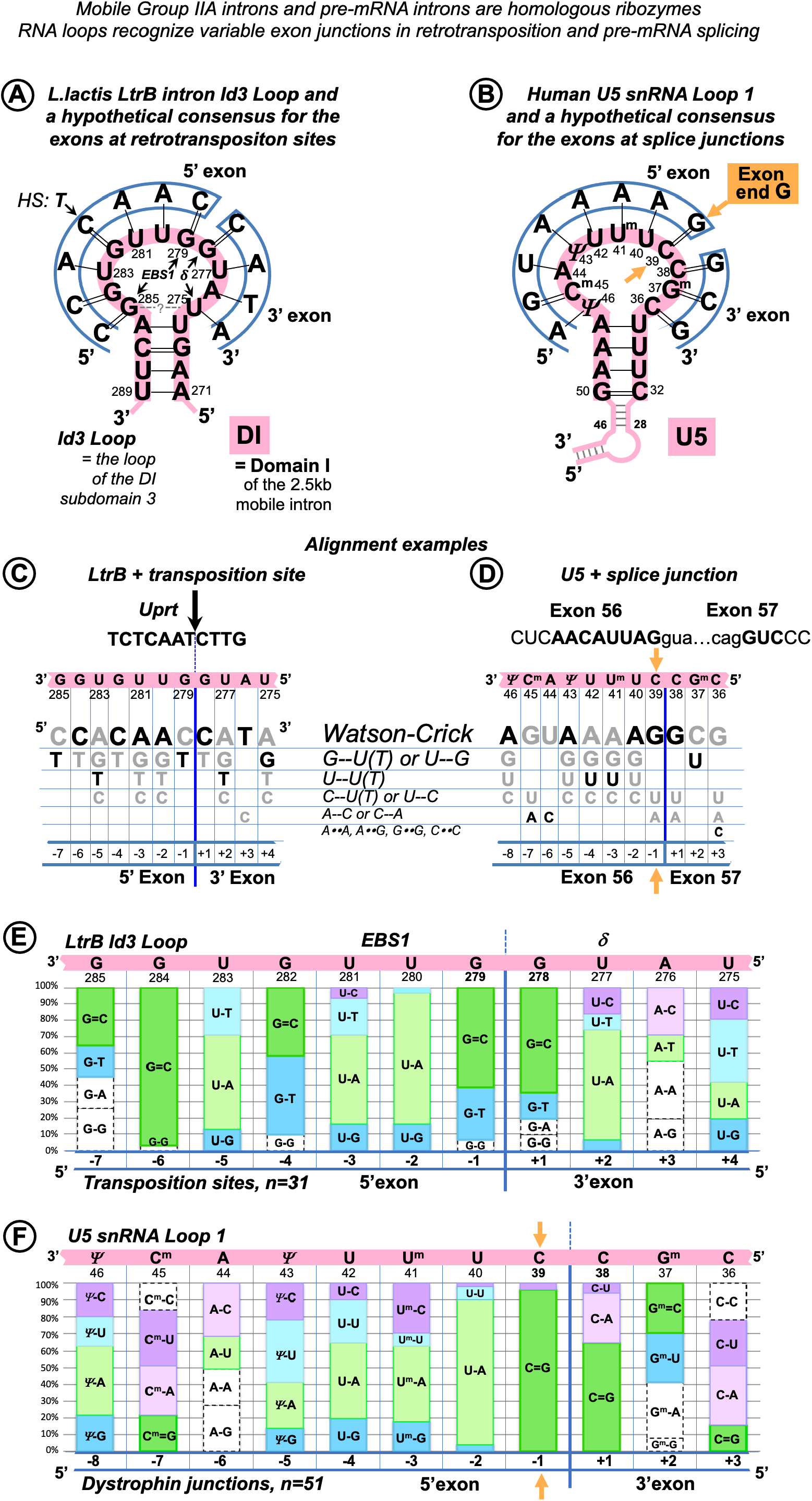
Base pair composition for the target recognition by mobile *L.l.*LtrB intron helps to identify the binding register for the human exons and U5 snRNA. A: The 11nt Id3 Loop in Domain I (**DI**) is the element of the *Ll.*LtrB intron responsible for the specific recognition of the exons. C=G pairs with guanines in Id3 positions 278 and 279 coordinate the splice junction. Id3 loop of the excised intron can pair with genomic targets similar to the homing site and guide retrotransposition. We derived a hypothetical consensus for retrotransposition sites based on the complementarity to Id3 Loop. The homing site (*HS*) in the *ltrB* gene differ from this consensus in the 5’ exon position -4 (thymidine). *EBS1*: 7 nucleotides of the Id3 loop (positions 279-285) pair with the end of the 5’ exon. *δ:* the remaining 4 nucleotides (positions 275-278) form a helix with the start of the 3’ exon. **B:** Assuming that like in Group IIA introns Watson-Crick pairs are preferred, we derived a hypothetical consensus complementary to U5 Loop1. The actual human splice junction consensus AG|G (Figure 1F) appears incorporated into this hypothetical sequence and G=C pairs with cytosines in U5 positions 38 and 39 coordinate the splice junction (orange arrow: U5 39C pairs the conserved exon-end G). **C:** We derived a grid for manual alignment of the retrotransposition sites with the Id3 loop that listed Watson-Crick and frequent mismatched pairs. In this way we recorded base pairs involved in the recognition by the LtrB intron of 31 targets in the *L.lactis* genome (Ichiyanagi et al., 2002; one example is shown here). **D:** Assuming that U5 snRNA Loop1 has the same base pair preferences as *Ll.*LtrB Id3 Loop and that the 5’ exon helix is longer than the 3’ exon helix we superimposed each of the 78 dystrophin gene splice junctions on the grid in the 5 possible binding registers (as in example here). Alignments with most Watson-Crick pairs were chosen as most likely with 65% of dystrophin exon junctions unambiguously aligned to U5 C_38_|C_39_ (as in **B** and **D)** and further 30% also fit this and one or two alternative binding registers. **D:** Summary for *Ll.*LtrB Id3 Loop of the total **n=341bp** with 31 retrotransposition sites (Ichiyanagi et al., 2002). **E:** Summary for human U5 snRNA Loop 1 of the total **n=561bp** with only 51 dystrophin splice junctions that anumbigously aligned with U5 C_38_|C_39_. Base modifications as in Figure 1 caption.

We examined the published sequences (Ichiyanagi et al., 2002) of retrotransposition sites (n=31) for base pair content of their interactions with *Ll*.LtrB Id3 Loop (**Figure 2C**). Apart from canonical Watson-Crick pairs (55%, 186 of 341), G-T/ U-G (17%) and U-T pairs (9%) appeared to be most frequent in these interactions (**Figure 2E**).

#### Binding register and base pair types in the interactions of U5 snRNA with the dystrophin exons

We examined possible U5 binding registers individually for 78 splice junctions of the human dystrophin full length skeletal muscle mRNA. We assumed that: 1) the end of the 5’ exon forms a longer helix with the recognition loop than the start of the 3’exon as in Group II introns; 2) the preferred pairs are Watson-Crick; 3) the types of frequent mismatched pairs are common for these RNA loops: we used the same grid, that lists all possible base pair types in order of frequency observed in *Ll*.LtrB retrotransposition (first Watson-Crick followed by mismatches G-U(T)/U-G, U-U(T), C-U/U-C) for the human U5 snRNA Loop1 (**Figure 2C,D**) to align manually 10 nt at the end of each exon joined to 5nt of the start of the next exon. The sequence was superimposed on the grid for all 5 binding registers that allow for a longer 5’exon helix and alignments with the most Watson-Crick pairs were chosen as most likely. 65% of dystrophin exon junctions unambiguously aligned to U5 positions C_38_|C_39_ (**Figure 2B**), a further 30% also fit this and equally one or two alternative binding registers. Therefore, a total of 95% of dystrophin mRNA splice junctions match the same U5 position, indicating that U5 C_38_|C_39_ is the optimal fixed binding position for the exon junction. This position is subsequently referred to as ‘the proposed binding register’ and used for the statistical analysis below.

As 5% of dystrophin exon junctions appear to match alternative positions better than U5 C_38_|C_39_, a possibility of an occasional shift of the U5 binding register cannot be outruled. A single relevant piece of evidence concerns the reverse splicing of a Group II intron into a mutant **h**oming **s**ite (HS, exon junction in the intronless allele): Su et al., 2001 reported a shift in the binding register by one nucleotide that secured a G=C pair.

While we assume that possible shifts in the U5 binding register are rare, the incorporation of non-canonical mismatched pairs alongside canonical Watson-Crick is inevitable in exon recognition helices. Accordingly, the base pair composition for dystrophin junctions that aligned unambiguously to U5 C_38_|C_39_ (n=51) was as follows: 45% Watson-Crick (252 of 561), 14% C-U/ U-C, 11% A-C/ C-A, 10% G-U/ U-G and 9% U-U (**Figure 2F**).

#### Common mismatched pairs are interchangeable for Watson-Crick pairs

What makes these mismatched pairs acceptable in the interactions of *L.l*.LtrB with diverse genomic targets, and the proposed interactions of U5 with the multitude of exon sequences? It appears that G-U, A-C, C-U and U-U pairs have an important quality in common: they can assume Watson-Crick-like geometry in different cellular molecular systems (Bebenek et al., 2011; Wang et al., 2011; Rozov et al., 2015; Rypniewski et al., 2016; detailed Discussion below). In effect, a single repositioning of a proton (prototropic tautomerization) or addition of a proton (protonation) for one of the bases in these pairs can produce configurations resembling the shape of the canonical pairs (**Figure 11)**.

Further in this paper these pairs are termed *‘isosteric’* as opposed to A-G, G-G, A-A and C-C pairs that are always distinct from Watson-Crick geometry and thus disrupt the architecture of the recognition helices. (The theoretically possible Watson-Crick-like C-C configuration requires both imino tautomerization and protonation – a pair not featured in any structures to date, Discussion, **Figure 11.**) For convenience, isosteric pairs are subsequently represented by a double dash: ‘G--U’, non-isosteric with a double dot: ‘G••A’, canonical Watson-Crick with a single dash for A-U and an equals sign for the triple-H-bonded G=C and non-isosteric ‘wobble’ pairs with a single dot: ‘G•U’. **Figure 2E** is the essential evidence of covariation of Watson-Crick and isosteric mismatched pairs. During self-splicing or retro-homing (reverse self-splicing) the *L.l.*LtrB Id3 Loop forms Watson-Crick pairs at every position of the splice-junction, except position -4 of the 5’ exon. Assuming that in retrotransposition the shape of pairs is the key to target recognition, G--U, A--C, C--U and U--U are acceptable only in their isosteric configuration. Remarkably, position -4 demonstrates a reciprocal example: during self-splicing or retro-homing the 5’ exon of the ‘home’ gene forms a U_-4_--G_282_ pair with the Id3 loop of LtrB intron. In retrotransposition, whereas 48% of integration sites conserve U_-4_--G_282_, 42% change to canonical Watson-Crick C_-4_=G_282_. Isosteric U--G with either base in enol configuration is a high frequency pair (previous NMR data - Kimsey et al., 2015 - discussed below) and as opposed to differently shaped wobble U•G explains the occurrences of U--G/G--U pairs in various positions in the interactions of many other Group II introns with the exons of their home genes (**Figure S1**).

**Figure 2F** presents a homologous covariation of Watson-Crick and isosteric pairs for the U5 Loop1 with human dystrophin gene exons. In particular, at position -3 of the 5’exon the proposed binding register shows covariation of C--U and A-U pairs. In the early spliceosomal complex, that precedes U5 binding, exon positions -1 to -3 interact with U1 snRNA C_9_U_10_G_11_ and select for exon-end C_-3_A_-2_G_-1_ (**Figure 1B**). In fact, in the dystrophin gene the ratio of C/A in position -3 is ¾ (in the whole human genome it is near 1:1, **Figure 1A**). Although Cryo-EM studies pictured 5’exon paired with U5 in different registers, position -3 was always aligned with one of the uracils (**Table S2**), effectively admitting the A-U/C--U covariation. In our model, exon position -3 pairs with U5 41-2’O-methyl-uracyl with covariation of the A_-3_-U^m^_41_ and C_-3_--U^m^_41_ (**Figure S3**, **Figure 1F)**.

In summary, we suggest that base pair geometry is the key to recognition of exon junctions by the spliceosome and retrotransposition sites by the LtrB GroupIIA intron. Watson-Crick pairs are selected for, isosteric pairs (G--U, A--C, U--U and U--C) are accepted, while pairs that perturb the helix architecture are kept out of these interactions. An important quality of uracil is that it can form isosteric pairs with any other base (**Figure 11**), so uracil-rich RNA loops like spliceosomal U5 Loop1 and *Ll.*LtrB Id3 Loop are useful for semi-specific sequence recognition, relying on isosteric mismatched pairs supported by Watson-Crick pairs to preserve the shape of the RNA helix. This mechanism explains how the universal U5 Loop1 can bind the multitude of diverse human exon junctions and equally explains *Ll.*LtrB intron mobility by reverse splicing into new genomic targets, with sequence ‘similarity’ defined by acceptable base pair geometry.

#### U5 Watson-Crick pairs with the exons in the proposed binding register compensate for substitutions of the conserved +5G in the dystrophin introns

Some dystrophin gene exons cannot form any Watson-Crick pairs with U5 snRNA in the proposed binding register. We noticed, that in such cases there is always a perfectly conserved U6 binding site G_+5_U_+6_A_+7_(U_+8_) at the start of the intron that forms Watson-Crick pairs with U6 positions (39)40 to 42 (**Figure 3B**). Conversely, among the 78 dystrophin gene introns, 18 (23%) lack the conserved +5G and all of these introns are preceded by exons that form multiple Watson-Crick pairs with U5 snRNA in the proposed binding register (**Figure 3D**).

**Figure 3.**
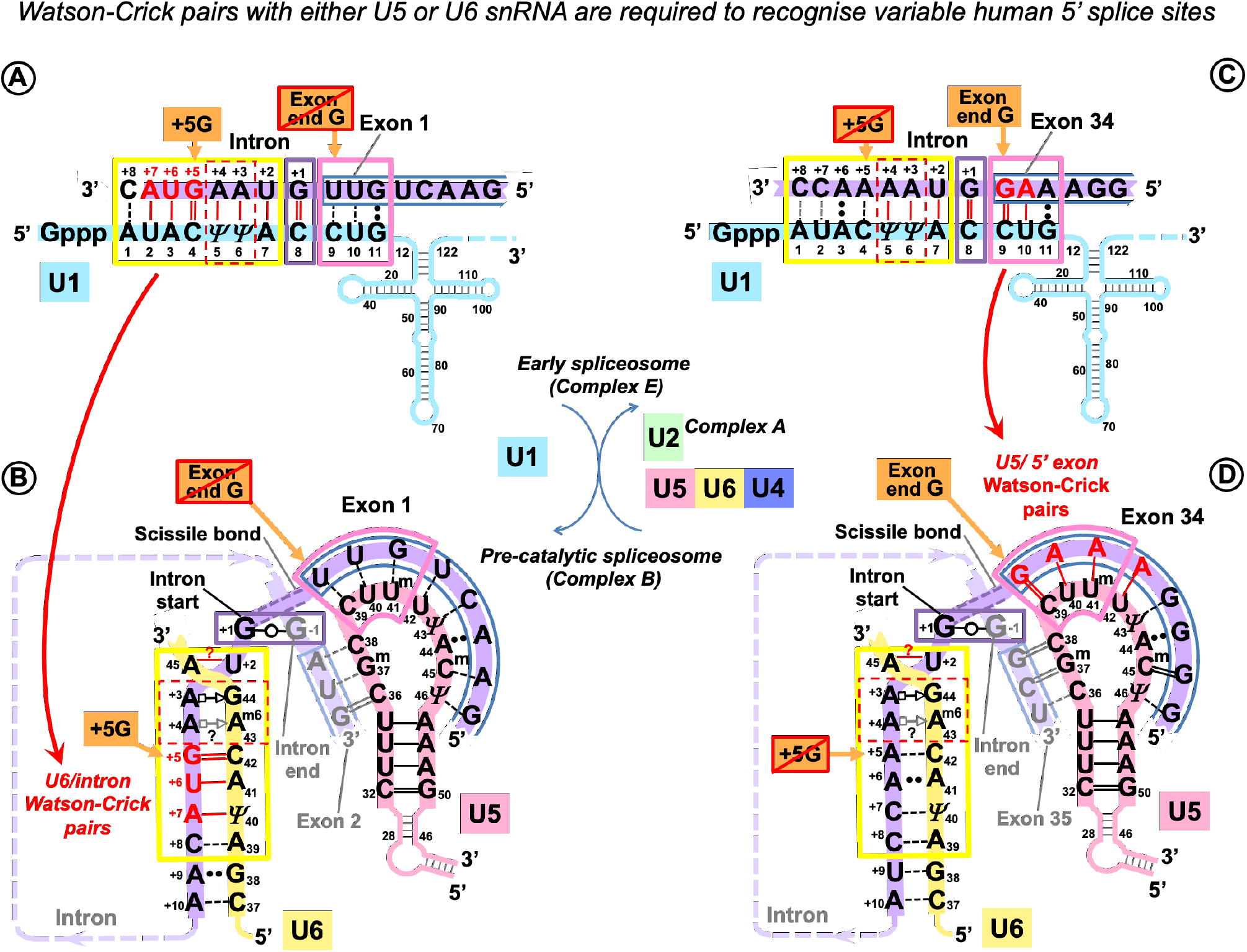
U5 and U6 recognise variable human exon/intron boundaries by Watson-Crick base pairing at the pre-catalytic stage and cooperatively ensure splicing fidelity A and B: In the early spliceosome (complex E) U1 snRNA forms on average 7 (minimum 5) Watson-Crick pairs with the exon/intron boundary (Carmel et al., 2004; Ketterling et al., 1999). U1 can bind multiple alternative and cryptic targets (Eperon et al., 1993; 2000) and is known to initiate correct splicing when bound in the vicinity rather than at the actual exon/intron boundary (Fernandez Alanis, et al., 2012; Singh and Singh, 2019), presumably leaving the fidelity check for the next stage. **C and D:** In the pre-catalytic spliceosome (complex B) U1 is replaced by U5 snRNA at the exon-end (pink boxes). U6 snRNA replaces U1 at the intron-start (yellow boxes). The well conserved adenines at intron positions +3 and +4 are enclosed in a red dashed box. Initially these adenines pair with U1 pseudouridines 5 and 6, then in the pre-catalytic complex they form non-canonical pairs with U6 _43_G_44_ (Figure 1D legend). Intron termini pair (purple box): see Figure 1E legend and Discussion **3.6.2**. **C:** Lack of exon complementarity to U5 is compensated by a strong intron interaction with U6 (as an example: human dystrophin intron 1). D: *Vice versa*, lack of intron complementarity to U6 is compensated by U5 Watson-Crick pairs with the 5’exon: (as an example: dystrophin intron 34). Base modifications as in Figure 1.

Effectively, in the human dystrophin gene U5 and U6 snRNAs mutually compensate for the loss of complementarity at their binding sites, stabilising the pre-catalytic complex with Watson-Crick base pairing. These observations in this small dataset hinted that it is the collective effect of U5 and U6 that ensures splicing precision in the context of variable splice signals of human genes.

### 2.2 Statistical testing of the new model of the interactions of U5 snRNA with human splice junctions

The pilot hypothesis indicates a distinctive binding register for the exons and U5 snRNA and places the splice junction so that the end of the 5’ exon is paired with U5 39C and the start of the 3’ exon binds U5 38C. This binding register appears to be linked to the mechanism of coordinated and mutually supportive splice signal recognition by U5 and U6 snRNAs.

In order to investigate if what is true for the dystrophin pre-mRNA is a general rule, we planned statistical tests that compare base pair distributions in the interactions of U5 and U6 snRNAs, placing U5 interactions according to the new model.

We also paid special attention in distinguishing the roles of U5 and U1 snRNA, which binds the last 3 positions of the exons during the initial selection of exon/intron boundaries. The focus of the series of statistical tests described below is to validate the functional importance of the new model of the U5 interactions with the exons.

#### 2.2.1 Dataset of base pairs in the interactions between human pre-mRNAs and snRNAs

Rather than scoring nucleotide distributions in exons and introns, we generated datasets of base pairs of their interactions with snRNAs. In order to enable analysis of the role of variation at specific positions in the splice junction, it is necessary to have a sufficiently large dataset of base pairs for U5 and U6 snRNAs at their pre-mRNA binding sites. Aiming to create a dataset of approximately 2000 introns, we paired *in silico* the **s**plice **s**ites (ss) from 132 well-studied human genes (supplementary **List S1**) with the snRNAs and computed for each ss position the frequency of base pairs grouped into three categories depending on their geometric properties. These are Watson-Crick pairs (G=C/C=G and A-U/U-A), isosteric pairs as defined above (G--U/U--G, U--U, C--U/U--C and A--C/C--A) and non-isosteric pairs (A••G/G••A, A••A, G••G and C••C).

The 132 genes contain 2007 introns and their respective exon junctions (**List S1**). 4 minor introns with U6atac binding site motif (processed by the alternative spliceosome, Discussion **3.6.2**, **Figure 10C**) were excluded from subsequent analysis (**List S2**). 13 atypical major introns with substitutions of the usual +2U (+2C in 12 introns and +2A in 1 intron, **List S3**) were also excluded from the analysis of the 5’ splice site interactions with U5 and U6, as the observed multiple Watson-Crick pairs on both sides of exon/intron boundary are likely to stabilize the unusual U6 A_45_--C_+2_(A_+2_) pair, rather than indicate any correspondence between the end of exon and start of intron positions +5 to +8. The final dataset consisted of 1990 major spliceosome GU_AG introns and their respective exon junctions.

#### 2.2.2 The effects of intron +5G and exon -1G substitutions at the 5’ splice site

The analyses below show that U5 snRNA contributes to precise definition of the 5’ splice site in the pre-catalytic complex forming more Watson-Crick pairs with the 5’ exon positions -5, -3, -2 and -1 to compensate for the loss of the +5G, the most conserved residue of the U6 binding site at the start of the intron. Reciprocally, U6 snRNA forms more Watson-Crick pairs with the intron positions +5 to +8 to compensate for substitutions of the most conserved exon-end guanine (-1G) of the U5/splice-junction interaction.

##### U5 Watson-Crick pairs with the 5’ exon compensate for substitutions of the conserved +5G in the following exon

For the first experiment we sorted exon junctions into two groups, the first one contained introns that conserved +5G and the second: introns with substitutions of +5G (+5Gsub).

Plotting the proportion of Watson-Crick, isosteric and non-isosteric pairs as a function of position, we observed an increase of Watson-Crick pairs between the 5’ exon and U5 snRNA in the absence of the conserved U6 C_41_=G_+5_ pair at the start of the intron (**Figure 4A-C**). However, there is no change in the base pair composition of the interaction of the 3’ exon with U5 between +5G and +5Gsub groups.

**Figure 4.**
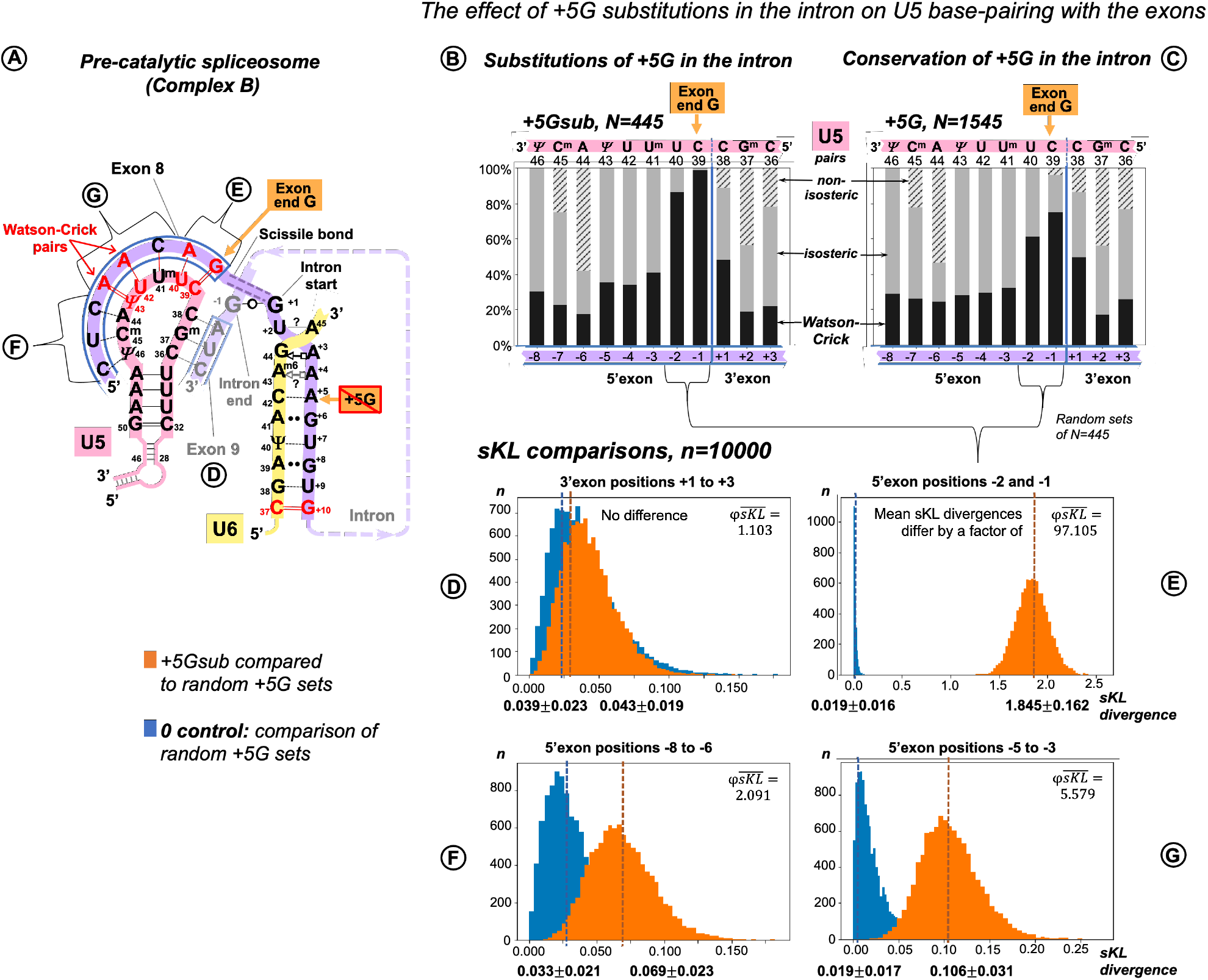
Additional U5 Watson-Crick pairs with the 5’exon compensate for substitutions of the conserved +5G at the start of the intron. **A:** Schematic of the interactions with U5 and U6 snRNAs (using, human dystrophin splice junction of exons 8 and 9 and the start of intron 8 as an example). The 11 base pairs of the U5 interaction with the exons are here subdivided into 4 groups that correspond to sKL distributions (**E-D). B:** Base pair frequencies for 445 human splice junctions related to introns lacking +5G (+5Gsub). **C:** Base pair composition for 1545 splice junctions of introns with conserved +5G. **D-F:** Distributions of symmetrised Kullback-Leibler divergences (sKL) for each positional group between the +5Gsub and 10000 random non-redundant +5G sets of n=445 (orange histograms) and control distributions of random non-overlapping and non-redundant sets of the same size from within the +5G dataset (blue histograms).

We compared the distribution of U5 base pair types by computing Kullback-Leibler (KL) divergence, a statistic used previously to compare distributions of nucleotides at splice sites (Sheth et al., 2006). Originating in information theory (Kullback and Leibler, 1951), KL divergence is a measure of relative Shannon entropy (variation) between two probability distributions, a cumulative statistic that sums up *all* the changes in the two distributions as logs of relative probabilities. The original KL divergence is not symmetric: KL(P,Q)≠KL(Q,P). The symmetrised KL (sKL) divergence is a sum of KL divergences of distribution P from Q and Q from P (*“there and back again”*). To assess the effect of base pair position we divided the splice junction into subsites (**Figure 4A**) and evaluated the extent of changes by sKL at each of these subsites (**Figure 4D-G).**

Our two datasets are naturally of unequal size: introns with conserved +5G, N=1545 and introns with substitutions +5Gsub, N=445. sKL divergence is a relative measure, so it is useful to have a control with ‘no difference’. Here as control we used pairs of random non-overlapping and non-redundant sets of N=445 drawn from the +5G dataset. One +5G set of each such pair was also compared to the +5Gsub dataset. 10000 iterations of these procedures returned distributions of sKL divergences within +5G sets (control) and between the +5Gsub dataset and +5G sets. If these distributions superimpose, there is essentially no difference between the two cases as exemplified by **Figure 4D** – the 3’exon. The sKL distributions are well separated in **Figure 4E** (mean sKL divergences differ by a factor of 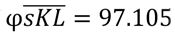), which shows that the base pair composition is very different at exon positions -2 and -1. This is easily seen in the increase proportion of Watson-Crick pairs with U5 snRNA in these positions in the +5Gsub dataset (**Figure 4B** vs. **4C**). An ∼17-fold smaller effect of the +5G substitutions is evident at 5’ exon positions -5 to -3 (**Figure 4G**, 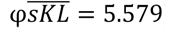). While in positions -8 to -6 sKL distributions almost superimpose (**Figure 4F**, 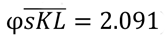), indicating no or little effect of the conserved intron position +5.

However, our comparisons of sKL divergence distributions shows the cumulative change of all the three base pair types at 2 or 3 positions of the splice junction at a time. The smaller change at 5’exon positions -5 to -3 is not convincing to distinguish between the U5 interaction in the pre-catalytic spliceosome and U1 interaction in the early spliceosome. To overcome these limitations, we measured variation of distinct U5 base pairs at individual positions in the splice junction corelated with substitutions of the conserved guanine at the intron position +5.

##### Functional importance of distinct U5 base pairs at specific positions

We can make a more detailed comparison and test for differences in the frequency for each type of base pair at each of the 11 splice junction positions between our two datasets: +5Gsub and +5G, using a bootstrap procedure (see Methods). Distributions of the bootstrap differences are summarised in 3 sets of violinplots, one set for each base pair geometry: Watson-Crick, isosteric and non-isosteric (**Figure 5A-C**). The grid line 0 represents the null hypothesis of no difference between the two datasets. The violin plots represent frequency estimates: the variations in the difference that could arise from variation in sampling of the transcriptome (i.e.: the uncertainty associated with the observed differences for our dataset). The null hypothesis p-value is indicated (statistically significant values are marked with asterisks – see Methods regarding multi-comparison corrections). The bootstrap differences above 0 indicate an increase of base pair frequency in the +5Gsub dataset compared to +5G dataset.

**Figure 5.**
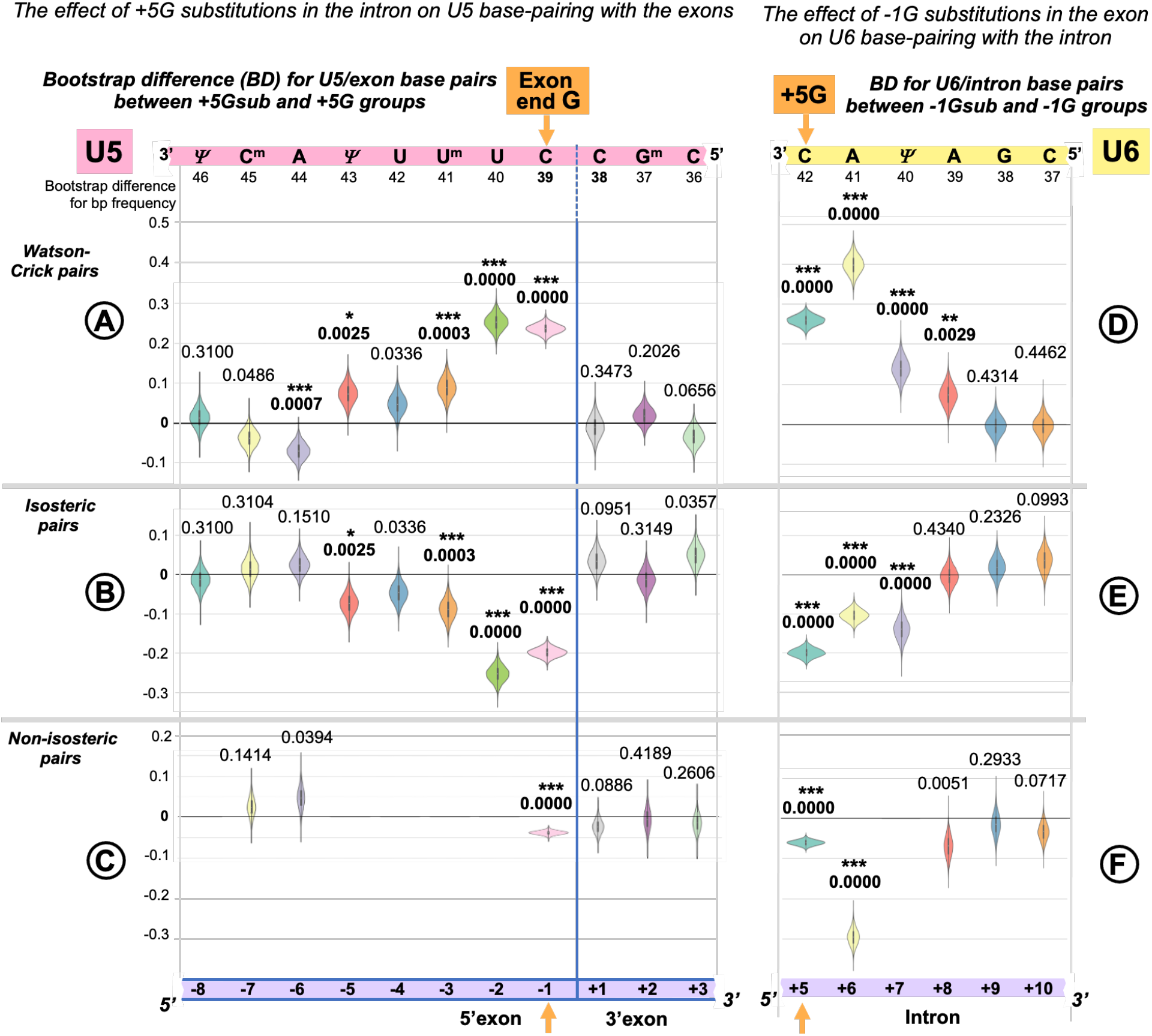
Reciprocal effects of +5G intron substitutions on the 5’ exon base pairs with U5 and the exon-end G substitutions on the intron base pairs with U6. Distributions of 10000 bootstrap differences for the frequency of Watson-Crick, isosteric and non-isosteric base pairs at each position of the exon junction (U5 biding site) between junctions flanking introns that conserve +5G and those where this base is substituted (**A-C**). Differences for the bp frequencies of the U6 binding site at the start of the intron positions +5 to +10 between introns preceded by exons that conserve exon-end G (-1G) and those that do not (**D-E**). The null hypothesis probability, *P*(*H*_0_) of no difference is indicated above each violin, asterisks mark significant changes after the correction for multiple testing (see Methods for details).

The violinplots show that the changes for the frequency of Watson-Crick pairs are largely mirrored by those of isosteric pairs (**Figure 5A, B**): these pairs replace each other in the interactions of U5 snRNA with the splice junctions. In accordance with sKL divergence evaluation, there are no substantial (or significant) changes in the base pair composition for the 3’ exon. Significant increase of U5 Watson-Crick pairs with 5’ exon positions -1, -2, -3 and -5 are observed, which apparently compensate for the absence of the key U6 C_42_=G_+5_ pair. The small decrease of Watson-Crick pairs at position -6 shows that a 5bp-long U5 interaction is sufficient, if it is a perfect helix rich in Watson-Crick pairs. Conversely, introns with the conserved +5G are preceded with exons that form fewer proximal Watson-Crick pairs with U5 snRNA and their helices have more Watson-Crick pairs in the distal positions.

There are 5 violins missing in **Figure 5C**, as non-isosteric pairs do not exist for these positions. 5 nucleotides out of 11 in the U5 snRNA Loop1 are uracils, which can form isosteric pairs with any base. The only change observed for non-isosteric pairs is that they completely disappear in position - 1 if +5G is missing in the following intron.

##### The effect of the +5G is specific to U5 rather than U1 snRNA interaction

In principle, it is possible that observed changes in U5 base pairs could be simply a consequence of the initial interaction of U1 snRNA with the 5’ splice site as this requires a threshold number of Watson-Crick pairs (5-6bp, Kettlering et al., 1999), which can be on either side of the exon/intron boundary. The evidence against this argument is twofold: First, we observe an increase of Watson-Crick pairs at 5’exon position -5 and the initial interaction with U1 snRNA does not extend to this position. Secondly, U1 and U5 have different preferences for base pairing at 5’exon position -3. We applied bootstrap analysis to nucleotide changes at position -3 linked to +5Gsub and +5G introns (**Figure 6**). There is a significant increase in the adenine required for forming Watson-Crick pairs with the U5 uracil U^m^_41_, and no significant change in cytosine for Watson-Crick pairs with U1 guanine G_11_. Thus, we can unambiguously link the observed changes to the interactions of U5 snRNA with the 5’exon.

**Figure 6.**
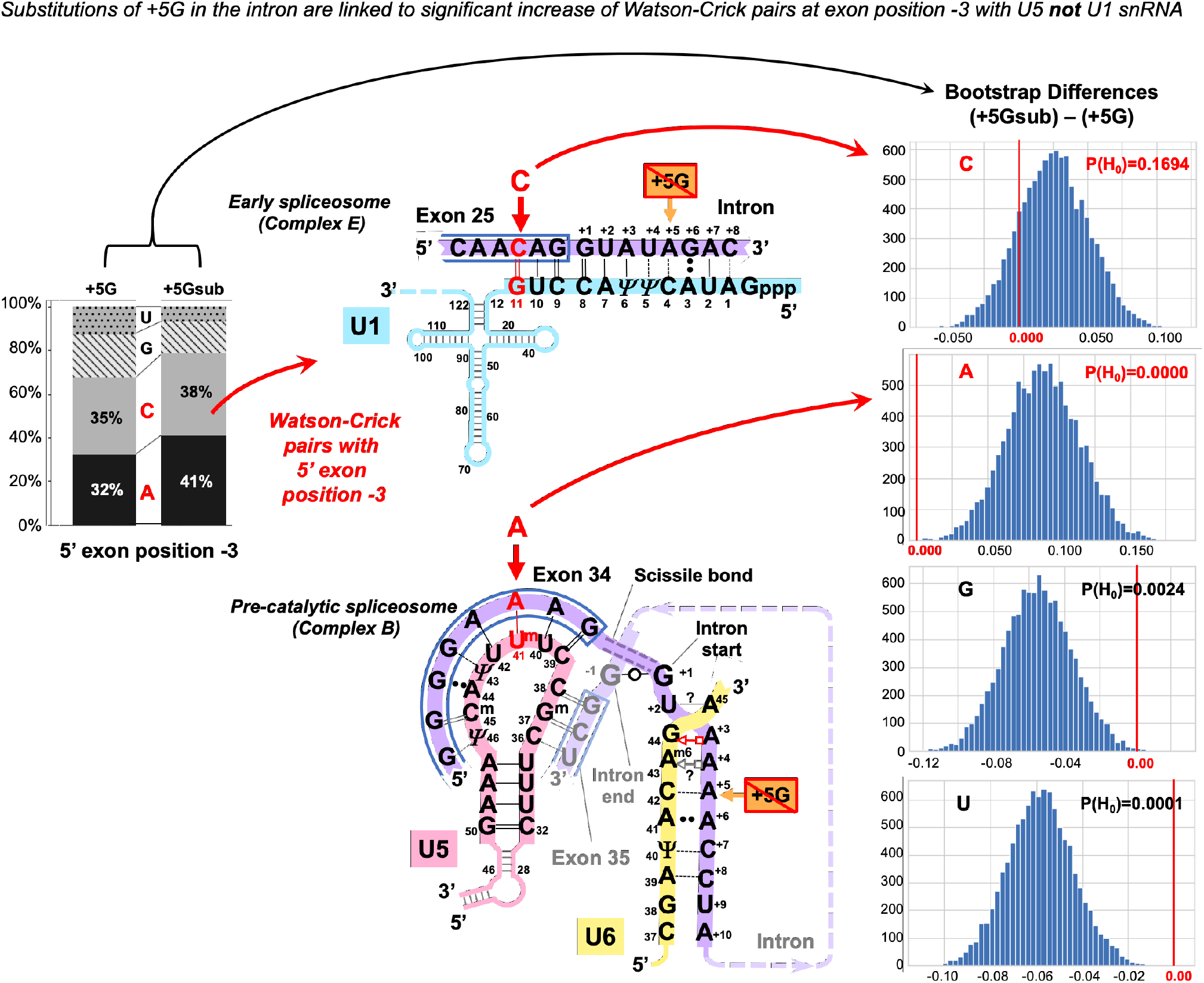
The +5G effect is specific to U5 snRNA, not U1 snRNA. The barchart on the left shows the distribution of nucleotides for the 5’exon position -3 in the presence and absence of the conserved +5G at the start of the intron. The schematics show the different nucleotide preferences for U1 and U5 paired to exon position -3 (indicated in red). Histograms for distributions of 10000 bootstrap differences for the frequency of each nucleotide at position -3 show that significant increase for A_-3_, creating a Watson-Crick pair in the U5 interaction, but not C_-3_ compensates for the loss of the U6 C_42_=G_+5_ pair.

##### U6 Watson-Crick pairs with intron positions +5 to +8 compensate for substitutions of the conserved -1G in the preceding exon

In a reciprocal experiment we separated introns preceded by exons that conserved exon-end G (-1G) and introns preceded by exons with -1G substitutions (-1Gsub). For the sKL divergence comparison we followed the same procedure as described for the exon junction (U5 binding site, +5G/+5Gsub, see above). Again, our two datasets were of unequal size: exons with conserved -1G, N=1598 and exons with substitutions -1Gsub, N=392, Consequently, the control in this case is provided by pairs of random non-overlapping and non-redundant sets of size N= 392 drawn from the -1G dataset. One of the -1G sets from each pair was compared to the -1Gsub dataset. Investigating dinucleotide subsets, we observed a strong base-pair type divergence corresponding to loss of the exon -1G at intron positions +5 and +6, a smaller effect at +7 and +8, and no effect further downstream (**Figure 7D-F**). We can see an increase of Watson-Crick pairs in positions +5 to +8 in the absence of exon-end guanine (**Figure 7B vs. 7C**).

**Figure 7.**
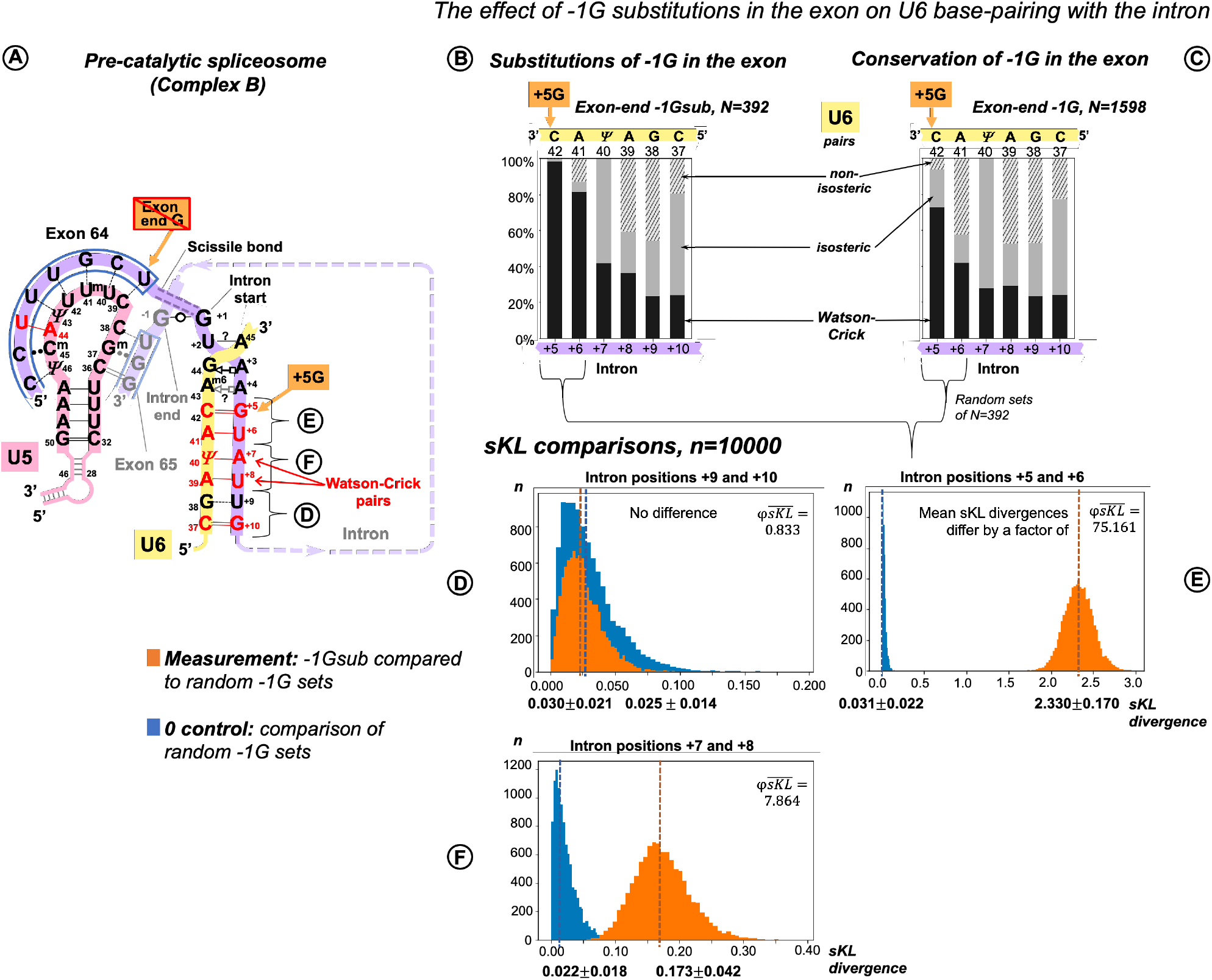
Additional U6 Watson-Crick pairs with the start of the intron at positions +5 to +8 compensate for substitutions of the conserved exon-end G. **A:** Schematic of the examined interactions with U6 and U5 snRNAs (using human dystrophin intron 64 and the splice junction of exons 64 and 65 as an example). The U6 interaction with the intron positions +5 to +10 is subdivided into dinucleotides that correspond to sKL distributions (**E-F). B:** Base pair type frequencies for 392 human introns preceded by exons lacking -1G (-1Gsub). **C:** Base pair composition for 1598 introns preceded by exons with conserved -1G. **D-F:** Distribution of symmetrised Kullback-Leibler divergences (sKL) for each dinucleotide between the -1Gsub dataset and 10000 random non-redundant -1G sets of n=392 (orange histograms) compared to a control distribution of the same size between random non-overlapping and non-redundant subsets from with the -1G dataset (blue histograms).

##### Functional importance of distinct U6 base pairs at specific positions

Exactly as we have done for U5 binding site (see above) we applied bootstrap resampling to test the null hypothesis of zero difference for the frequency of each base pair type at each of the six intron positions +5 to +10.The result is summarised in 3 sets of violinplots (**Figure 5D-F**).

There is a rise in Watson-Crick pairs at positions +5 through +8 with the largest rise in position +6 (**Figure 5D**). The importance of these interactions downstream of position +6 only becomes apparent in the absence of -1G, with no detectable genomic conservation (**Figure 1**). The nature of changes is somewhat different from the reciprocal changes of the U5 binding site, as the increase of Watson-Crick pairs is accompanied by significant decrease of both isosteric and non-isosteric pairs (**Figure 5F**). This is a result of non-isosteric pairs being tolerated at these positions in the dominant -1G case (**Figure 7C**), reflecting less constraints on the geometry of the helix for U6 binding with the intron, than for U5 Loop1 presenting the exon junction in the catalytic core of the spliceosome.

##### Explaining rare introns missing both the conserved +5G and end of exon -1G

In our sample of 1990 human introns, 6 (0.3%) lacked both these conserved guanines. We observed that multiple other Watson-Crick pairs stabilise both U5 and U6 interactions in these cases. (supplementary **List S4**).

#### 2.2.3 The effect of intron -3C substitutions at the 3’ splice site

The bootstrap difference analysis below shows that the 3’ intron end substitutions of the conserved -3C are supported by the increase of U5 Watson-Crick pairs with the 3’ exon position -1. This indicates a similar role of U5 in the correct recognition of the 3’ splice site, as at the 5’ splice site. The timing of the 3’ exon interaction with U5 and a possible RNA partner for intron position -3 are addressed in the Discussion **3.6.**

##### U5 Watson-Crick pairs with the 3’ exon compensate for substitutions of the conserved -3C in the preceding intron

Following up on our observation that in the human dystrophin gene the absence of -3C in the intron makes Watson-Crick pairs with the 3’ exon twice as likely, we sorted our sample of exon junctions of the major introns (N=2003, inclusive of introns with +2U substitutions, see above) into two subsets: - 3Csub, N=792 and -3C, N= 1211 (**Figure 8**). In this case over this large sample the differences are smaller than for the previous comparisons (**Figure 8B-F**).

**Figure 8.**
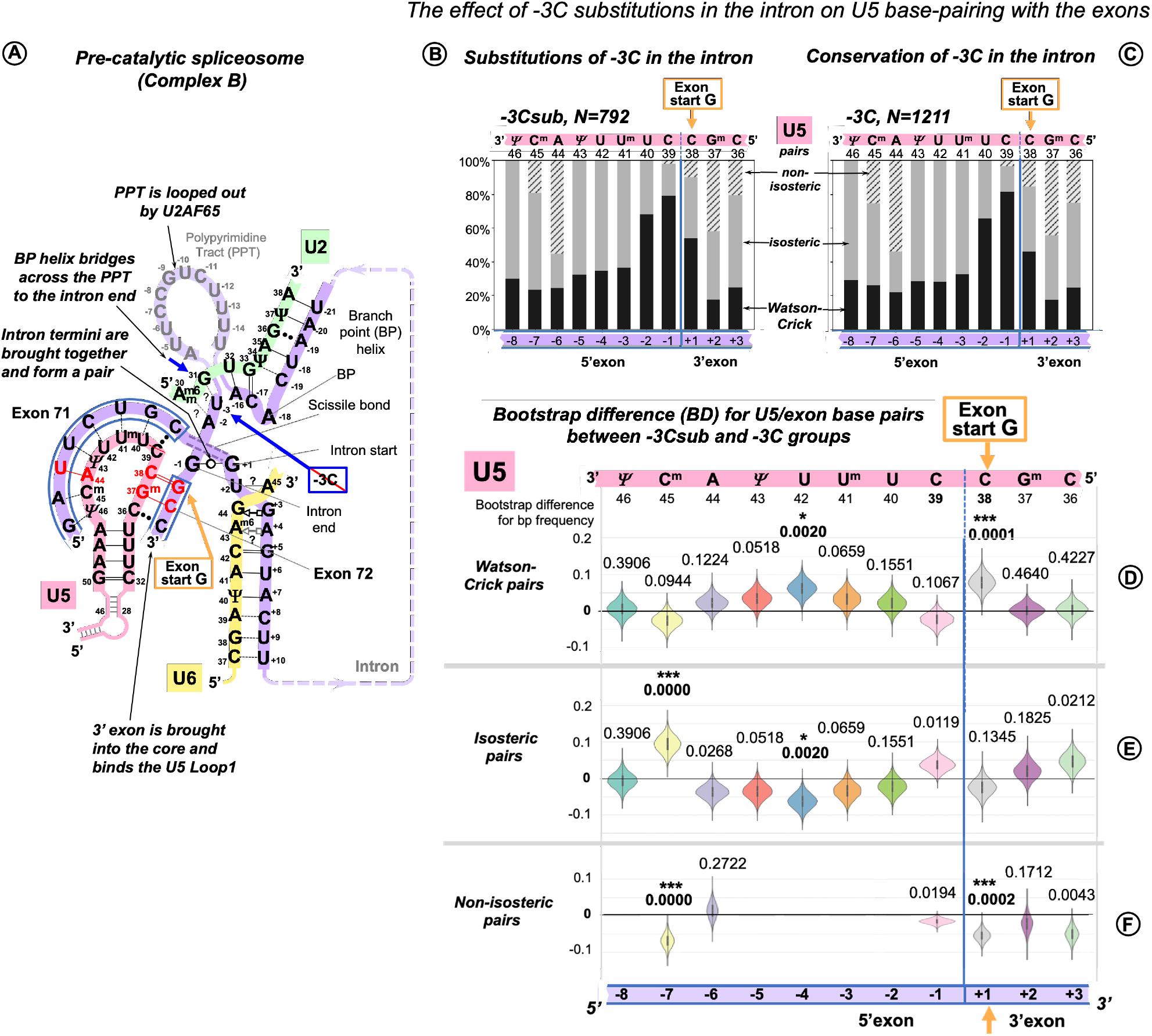
Additional U5 Watson-Crick pairs with the 3’ exon compensate for substitutions of the conserved -3C at the end of the intron. **A:** Schematic of the examined interactions with U5 and U2 snRNAs (using human dystrophin splice junction of exons 71 and 72 and intron 71 as an example). We propose (Discussion **3.6.1**) that BP helix bridges across the PPT to the end of the intron secured by the conserved -3C paired with U2 G_31_. This agrees with the previous mutation evidence (Corrionero et al. 2011; Brock et al. 2008), biochemical studies (Kent et al., 2003; Chen et al. 2010) and X-ray structures (Kent et al., 2003; Sickmier et al. 2006). The example here shows U2 G_31_--U_-3_ pair instead, as dystrophin intron 71 lacks -3C. **B:** Base pair type frequencies for 792 human splice junctions related to introns lacking -3C (-3Csub). **C:** Base pair type composition for 1211 splice junctions of introns with conserved -3C. **D-F:** Distributions of 10000 bootstrap differences for the frequency of Watson-Crick, isosteric and non-isosteric base pairs at each position of the splice junction (U5 binding site). The null hypothesis probability, *P*(*H*_0_) of no difference between the two cases is indicated above each violin, asterisks mark significant changes after correction for multiple testing (see Methods for details).

In the absence of -3C, U5 Watson-Crick pairs do increase at position +1 of the 3’ exon replacing non-isosteric pairs thus strengthening the U5 interaction with the 3’ exon. The 5’exon interaction with U5 shows an increased proportion of Watson-Crick pairs centred at position -4 and a rise in isosteric pairs at position -7 due to a drop in non-isosteric pairs, possibly indicating that stabilising distal positions of the 5’exon helix is important for the intron complex overall (**Figure 8A**).

### 2.3 U5 Watson-Crick pairs with the 3’ exon promote inclusion of exons with +1G mutations

The effect of human mutations of the conserved exon-end guanine (-1G) is currently explained by the base pairing with U1 snRNA, so we cannot use it as an evidence to support our new U5 model. Therefore, we concentrated on the mutations of the exon-start guanine (+1G). However, mutation databases do not document the effect of these mutations on splicing. Thankfully, Fu et al., 2011 specifically examined 14 mutations of the exon-start guanine and quantified their effect on exon inclusion using minigene constructs in human cells (HEK293). Each measurement was a mean result of a triplicate experiment. The authors report that 6 of these +1G mutations (in *LPL* exon 5, *HEXA* exon 13, *LAMA2* exon 24, *NEU1* exon 2, *COL6A2* exon 8, *COL1A1* exon 23) did not have any effect on exon inclusion at all (percent spliced in, PSI=100%). *PKHD1* exon 25 +1G → *T* mutation resulted in a cryptic 3’ splice site activation with 99% inclusion of a longer exon. While the splicing effect for the other seven +1G mutations (in *CAPN3* exon 10, *CLCN2* exon 19, *EYA1* exon 10, *COL1A2* exon 37, *FECH* exon 9, *GH1* exon 3 and *CAPN3* exon 17) did not involve any cryptic sites and varied from 91% PSI to complete exon skipping (respectively). Variable branchpoint sequences did not offer a clear explanation, instead Fu and colleagues proposed that long polypyrimidine stretch promotes exon inclusion in spite of +1G mutations. However, reducing the length of this stretch to 5bp in *LPL* exon 5 minigene still produced PSI of 63-83% (depending on the position of pyrimidines). Only 2 pyrimidines in *HEXA* exon 13 minigene resulted in PSI of 59-69%. The observed highly variable efficiency of exon inclusion with +1G mutations and the fact that the length of PPT does not always provide a clear explanation points out that other factors are also involved: the branchpoint helix and the conserved intron position -3 are also expected to contribute to splicing efficiency. Indeed, PSI was brought down to 7% for *HEXA* exon 13 minigene when -3C was substituted to G.

We re-examined the exon sequences for these fourteen +1G mutations, looking specifically for cytosine in exon position +2 and guanine in exon position +3, because they form Watson-Crick pairs with U5 C_36_G^m^_37_ according to our proposed binding register. We found that cytosine occurs in position +2 in two mutations that did not affect splicing (PSI=100%: *GH1* exon 3 and *FECH* exon 9) and in *CAPN3* exon 10 with 91% correct exon inclusion. Guanine occurs in position +3 in further 2 mutations with 100% PSI: *LAMA2* exon 24, *NEU1* exon 2. Finally, both +2C and +3G are involved in the cryptic 3’ss activated by +1G mutation in *PKHD1* exon 25. If we plot exon inclusion efficiency (PSI) for +1G mutations with +2C/+3G and without +2C/+3G, we can see that while the latter group is highly variable as can be expected if many factors are involved, the former group is clearly clustered at the top end, indicating that the presence of +2C or/and +3G is a very strong factor that promotes exon inclusion in spite of +1G mutations (**Figure 9A).** Although both ANOVA with Welch’s correction for unequal variances (the greater variance for the larger group makes false negatives more likely, McDonald, 2014) and non-parametric Kruskal-Wallis rank sum test show that there are significant differences between means and locations of these two groups, statistical tests for N=14 are implied only to complement the obvious differences between the boxplots (**Figure 9A**).

**Figure 9.**
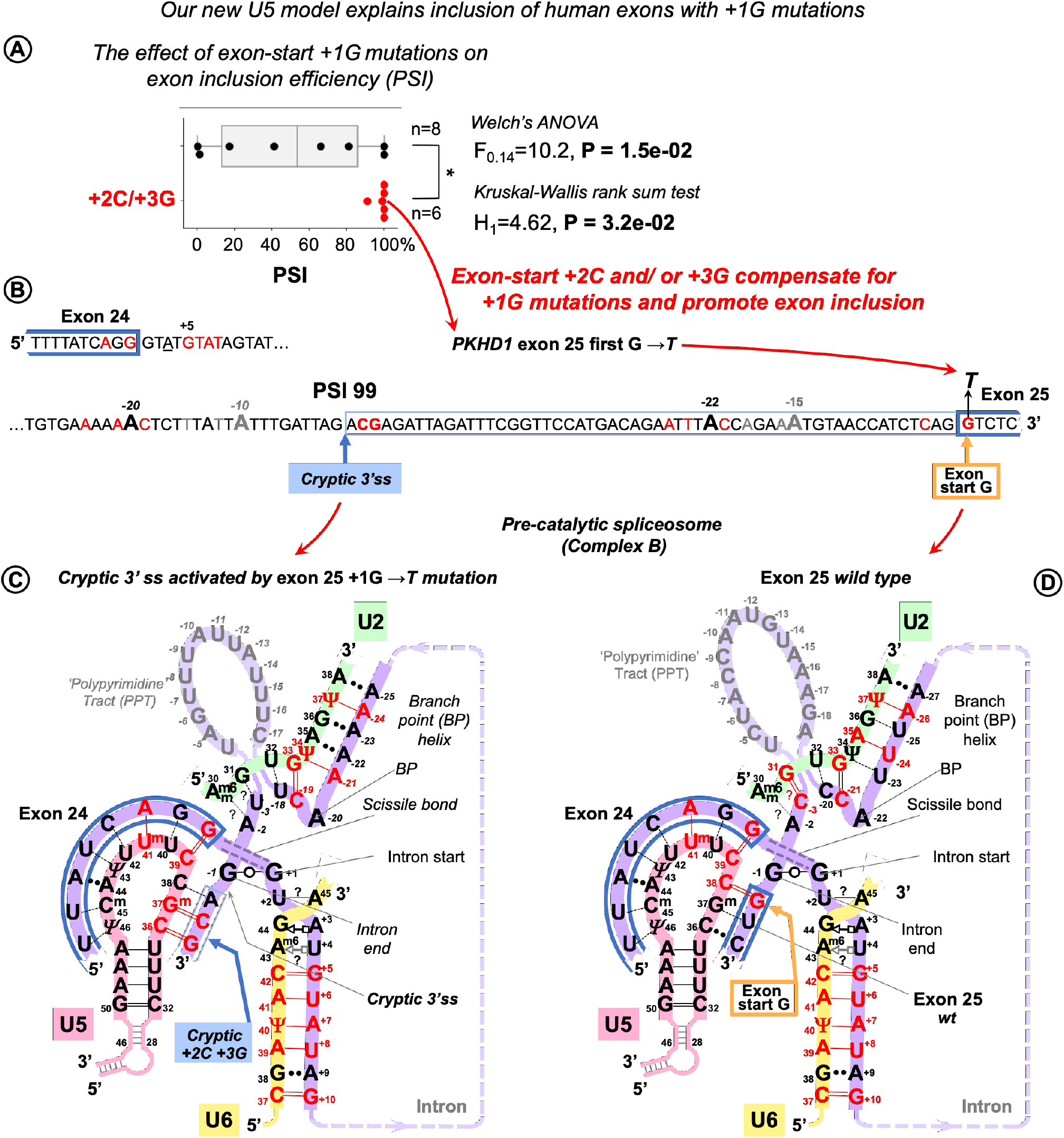
U5 Watson-Crick pairs with exon positions +2 and +3 promote inclusion of exons with +1G mutations. Fu et al., 2011 quantified the variable effect on splicing for 14 exon-start +1G mutations in human genes. **A.** Boxplots show that exon inclusion (**PSI**, percent spliced-in) is strongly influenced by the presence of exon +2C or +3G. **B.** Splice site sequences of the *PKHD* gene intron 25 and the flanking exons. +1G mutation in exon 25 completely blocks normal 3’ splice site and activates a cryptic 3’ss tag|A<u>CG</u> leading to the predominant inclusion of a longer exon and only 1% exon-skipping. **C.** Base pairing scheme for the cryptic 3’ss with U5 snRNA Loop1 secured by U5 G^m^_37_=C_+2_ and C_36_=G_+3_ according to our new U5 model. **D.** Base-pairing of the normal wt exon 25 with U5 snRNA. +1G mutation abolishes the U5 C_38_=G_+1_ pair which leads to exclusive use of the upstream cryptic 3’ss. **C,D**: Recognition of all splice sites is complete in the pre-catalytic spliceosome (complex B) before *Brr2* promotes catalytic core formation (see **Discussion 3.6**).

We further compared the effect of exon +2C/+3G with other factors, that are expected to influence exon inclusion efficiency (**Figure S11, Table S4**) Apart from aforementioned PPS length (as in Fu et al., 2011), +1G → *A* mutation is much better tolerated than +1G → *T*, so a change for purine emerges as a second strongest factor after +2C/+3G, which is to be expected, as generally in the human exons +1A is twice more likely than +1T. An example of the cryptic 3’ss activated by +1G mutation in *PKHD1* exon 25 is detailed in **Figure 9B-D.**

Identifying that exon +2C and +3G compensate for +1G mutations and strongly promote exon inclusion provides a clear explanation of the human mutation analysis and allows to conclude that the interaction of the 3’ exon with U5 Loop1 in the proposed binding register plays an important role in splicing precision. Moreover, this interaction of the 3’ exon, now confirmed by the mutation analysis is possible only for the fully open 11nt U5 Loop1 that we consider, as opposed to the 7nt version that prevails in Cryo-EM reconstructions (Discussion compare **Figure 12A** and **B**).

## 3 Discussion

### 3.1 The U5 hypothesis summary

- **Optimal binding register for diverse exons and U5 snRNA Loop1: the exon junction is positioned at U5 C_39_|C_38_**. This U5/exons model is based on homologous interactions of a mobile GroupIIA intron Id3 Loop with genomic retrotransposition sites in bacteria.
- **Common mechanism of base pairing for U5 snRNA with diverse human exons and *Ll*.LtrB intron with new loci in retrotransposition**: we suggest that these RNA loops recognize their variable target sequences by helix architecture, accepting Watson-Crick and isosteric base pairs and rejecting geometrically different pairs.
- **Significant role of U5 snRNA in specific exon recognition in the pre-catalytic spliceosome.** U5 Watson-Crick pairs with the exons in the proposed binding register compensate for substitutions of the conserved intron positions. In addition, our binding register explains human mutation data: U5 Watson-Crick pairs with exon positions +2 and +3 compensate for +1G (exon-start) mutations and strongly promote exon inclusion.

This last point, based on statistical analyses of base pairs at specific positions and further supported by human mutation evidence, directly proves the first point that the exon junctions is positioned at U5 C_39_|C_38_. (The timing for the 3’ exon interaction with U5 Loop1 is specially discussed in Section **3.6.2.)** The second point on the geometric sequence recognition cannot be directly tested by statistics, however it is our explanation of the observed common base pair types used by both RNA loops.

### 3.2 Modelling U5 Loop1 base pairing with human exons on Group IIA intron interactions with retrotransposition sites

Our new model of the interactions of the exon junction with U5 Loop1 is inspired by the homologous interactions in *Ll.*LtrB, bacterial GroupIIA intron (**Figure 10**).

**Figure 10.**
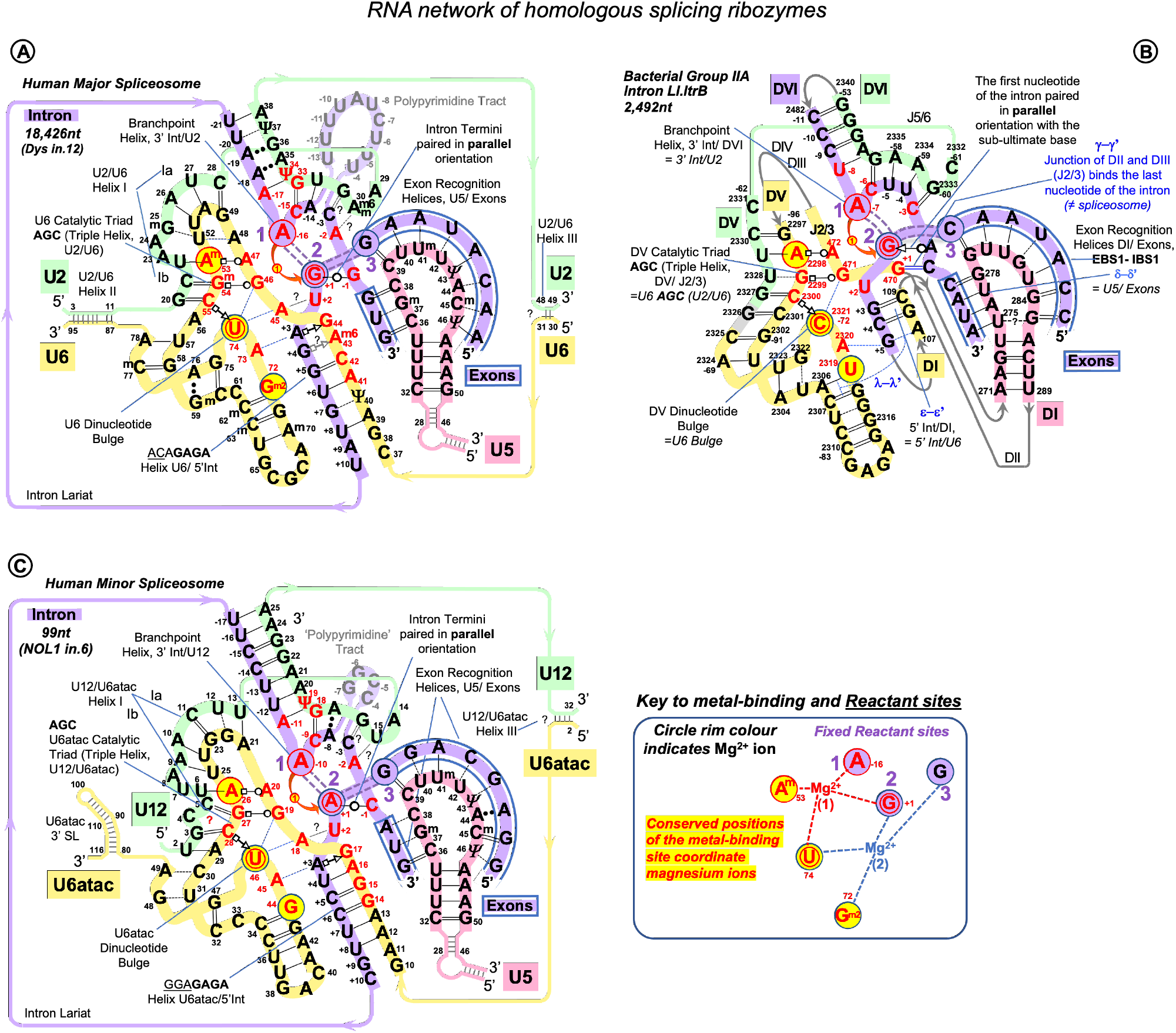
RNA network of the homologous ribozymes: human major and minor spliceosomes and GroupIIA intron. 1st catalytic step, spliceosomal complex C (successive spliceosome complexes are detailed in **Table S1**) The nucleophilic attack by the branchpoint adenosine: curved red arrow. The intron breaks off the 5’exon end and bonds with the 2’O of the branching A: double purple dashed lines indicate the scissile (purple fill) and emergent (no fill) covalent bonds. Splicing catalysis requires two Mg^2+^ ions at a fixed distance from three reactant sites (Steitz and Steitz, 1993; Fica et al., 2013). At the first catalytic step Mg^2+^(1) activates 2’OH of the BP A in the <u>Reactant site 1</u>. Mg^2+^(2) stabilizes the leaving 3’OH of the last nucleotide of the 5’ exon in <u>Reactant Site 3</u>. Both magnesium ions form a complex with the scissile phosphate of the N_+1_ of the intron in <u>Reactant site 2</u>. **A:** Human major spliceosome (intron 12 of the dystrophin gene as an example). The ribozyme is an assembly of three separate snRNAs with a record number of modified residues. The structure of the U6/U2 catalytic triplex is inferred from Galej et al. 2016; Keating et al. 2010 and U6/intron duplex as in Fica et al. 2017. Non-canonical RNA pairs are shown with Westhof geometric symbols (Leontis et al., 2002). Tertiary interactions as in Anokhina et al., 2013: blue dashed lines; Mimic Watson-Crick-like base pairing: black dashed lines; Base pairing with unknown non-Watson-Crick geometry: double dots. Base modifications as in Figure 1 and G^m2^: N2-methylguanosine. **B:** LtrB Group IIA intron *(Lactococcus lactis)*. Interactions of the catalytic triplex are extrapolated from *O.i*. structure (Keating et al. 2010). Core motifs of this large RNA molecule are coloured as homologous RNA components of the spliceosome. Greek letters: tertiary interactions in Group II introns, shown in blue. γ−γ’ and λ−λ’ interactions do not have homologues in the spliceosome. All other core interactions and catalytic structures of the ribozyme are labelled with spliceosome homologues in *italics*. Domains of the *Ll*.LtrB ribozyme: DI-DVI; Junctions between domains II and III or V and VI: J2/3 or J5/6. Double numbering is used for the residues starting from domain V, the negative number indicating position from the 3’ end. **C:** Human minor spliceosome (Tarn and Steitz, 1996a,b; Widmark et al., 2010; Ederly et al., 2011; He et al., 2011; Younis et al., 2013; reviewed in Turunen et al., 2013). U5 is the only snRNA shared with the major spliceosome. A lot fewer residues are modified in U12 and U6atac snRNAs compared to U2 and U6 paralogues. Perfect conservation of the BP helix and the U6atac snRNA AAGGAGAGA box interaction with the 5’ intron end is characteristic of the minor spliceosome. (**?**): an odd U12 C_4_ bulge (see **Comment S1**) here reproduced as in Tarn and Steitz, 1996b and Turunen et al., 2013. The minor introns are expression regulators of critical genes: the example here is intron 6 of the human **<u>N</u>**ucle**<u>ol</u>**ar Protein **<u>1</u>** (NOL1/NSUN1; Brock et al., 2008) gene encoding an RNA:5-methylytosine-methyltransferase (known as proliferation antigen p120 overexpressed in virtually all types of cancer cells).

Like Group II introns, human spliceosome is a metalloribozyme: protein-free small nuclear RNAs U6 and U2 are capable to catalyse splicing *in vitro* (Valadkhan et al., 2007; Jaladat et al., 2011). The core RNA components of the catalytic spliceosome resemble closely the domains of the Group II intron (**Figure 10A,B**): the branch point helix with the adenosine bulge, the intron termini pair with the parallel orientation of the RNA strands (discussed in **3.6.2**) and the catalytic metal binding site (Nguyen et al., 2015; Zhao and Pyle, 2017; Galej et al., 2014; Fica et al., 2013; Keating et al., 2010; Galej et al., 2018). The similarities are so great that the studies of the spatial organisation and mechanism of pre-mRNA splicing are much in debt to the structural and genetic studies of Group II introns. Both in the spliceosome and in Group II introns the two-step splicing mechanism (Steitz and Steitz, 1993) proceeds through 2’O nucleophilic attack or ‘branching’ of the sugar-phosphate backbone at the adenine base leading to the formation of an intron lariat excised after the exon ligation. Both steps of splicing are reversible. Group II introns use reverse splicing for retrohoming into the intronless alleles or retrotransposition into other genomic loci with sequence similarities (Lambowitz and Zimmerly, 2011; Lambowitz and Belfort, 2015; Eskes et al., 2000; Ichiyanagi et al., 2002; Zhong and Lambowitz, 2003; Griffin et al, 1995). Reverse splicing by the spliceosome was demonstrated *in vitro* (Tseng and Cheng, 2008) and suggested to be implicated for splicing quality control (Smith and Konarska, 2008).

In focus here are homologous U5 Loop1 and Group IIA Id3 Loop. Both these loops are 11nt long and contain 5 uracils. They bind both 5’ and 3’ exons aligned for ligation in the forward splicing process and the exons to be separated by the intron precisely at the junction in the reverse splicing process.

However, Group IIA intron self-splicing is based on near perfect complementarity with the exons (**Figure S1**). On the contrary, pre-mRNA splicing and Group IIA intron retrotransposition are equally challenged by variable exon junctions, and we looked for a common mechanism of sequence recognition by these homologous RNA loops.

The published data on retrotransposition of LtrB intron in *Lactococcus lactis* genome loci shows without doubt that the binding register for Id3 Loop and the ‘exon’ junctions in retrotransposition stays fixed and is the same as for the intron self-splicing: 7 positions of the Id3 loop pair with the sequence upstream of the intron insertion as with the 5’ exon and 4 positions of the loop form base pairs downstream of the retrotransposition site as with the 3’ exon (Ichiyanagi et al., 2002, **Figure 2A,C**). Retrotransposition sites are ‘similar’ to the homing site in a sense that they have on average 55-53% of sequence identity to the exon junction of *L.lactis ltrB* gene interrupted by LtrB intron. However, we gained a better insight into the mechanism of sequence recognition when we observed that the mismatched pairs are not random, and the preferred mismatches are limited to G--U/ T--G, T--U and C--U (**Figure 2E**).

By analogy, we manually aligned U5 Loop1 with the exon junctions for human dystrophin with maximum possible Watson-Crick pairs and the same preferred mismatches as for Id3 Loop and found that indeed 95% of dystrophin junctions align to the same U5 positions and that the mismatched pairs are not random: C--U/ U--C, A--C/ C--A, G--U/ U--G and U--U are strongly preferred (**Figure 2B,D,F**). The mechanistic explanation for this preference is discussed in the next section **3.3**. However, the first point of the U5 hypothesis, which we seek to prove by statistical analysis (see section **3.4**) is that U5 Loop1 has a fixed optimal binding register for human exons: the end of the 5’ exon pairs with U5 C_39_ and the start of the 3’ exon pairs with U5 C_38_. This is contrary to the CryoEM models for U5 Loop1 of which the most recent places the conserved guanine at the end of the 5’ exon paired with U5 U_40_. Alignment of the interacting RNA sequences is an obvious starting point (surprisingly, it was never previously performed for U5 Loop1 and the exon junctions of pre-mRNA introns), however, the way to prove that our proposed U5 binding register is true, can be to show that it has a role in exon recognition, as is the case for Id3 Loop of Group IIA introns. Our statistical analysis indeed shows this role: U5 Watson-Crick pairs with the exons in the proposed binding register compensate for substitutions of the conserved +5G and -3C in the intron splice sites (discussed in section **3.4**). Moreover, our model explains the effect of mutations in human exon sequences, which cannot be explained by Cryo-EM models (Discussion **3.5**).

### 3.3 The explanation for acceptable mismatches can be base pair geometry

The geometric principle for variable exon junction recognition in Group IIA intron retrotransposition and pre-mRNA splicing was suggested by the mismatched pairs *Ll*.LtrB Id3 loop and human U5 Loop1 employ: G--U, U--U, A--C and C--U. Bountiful literature on Watson-Crick like geometry of these base pairs is very briefly discussed below.

In order to explain spontaneous mutagenesis in replication, Watson and Crick themselves put forward the idea that G--T or A--C pairs can assume dimensions of canonical pairs if one of the bases adopts its rare tautomeric configuration (Watson and Crick,1953; **Figure 11A,B**). X-ray crystallography provided evidence of a G--T pair mimicking WC geometry in the active site of the human DNA polymerase λ and likewise an A--C pair adopting a clear WC like shape within the active site of *Bacillus stearothermophilus* DNA polymerase I (Bebenek et al., 2011; Wang et al., 2011; reviewed by Kimsey and Al-Hashimi, 2014 as ‘high energy purine-pyrimidine base pairs’). Apart from provoking mistakes in DNA synthesis, the biological significance of mismatched pairs assuming WC-geometry became further apparent, when crystal structures of codon-anticodon duplex of *Thermus thermophilus* 70S ribosome revealed that G--U mismatches in 1st and 2nd positions are isosteric to canonical pairs (Rozov et al., 2015; Westhof 2014a, Westhof et al., 2014b). This finding proves that mispairs mimicking WC geometry are also responsible for translational infidelity.

**Figure 11.**
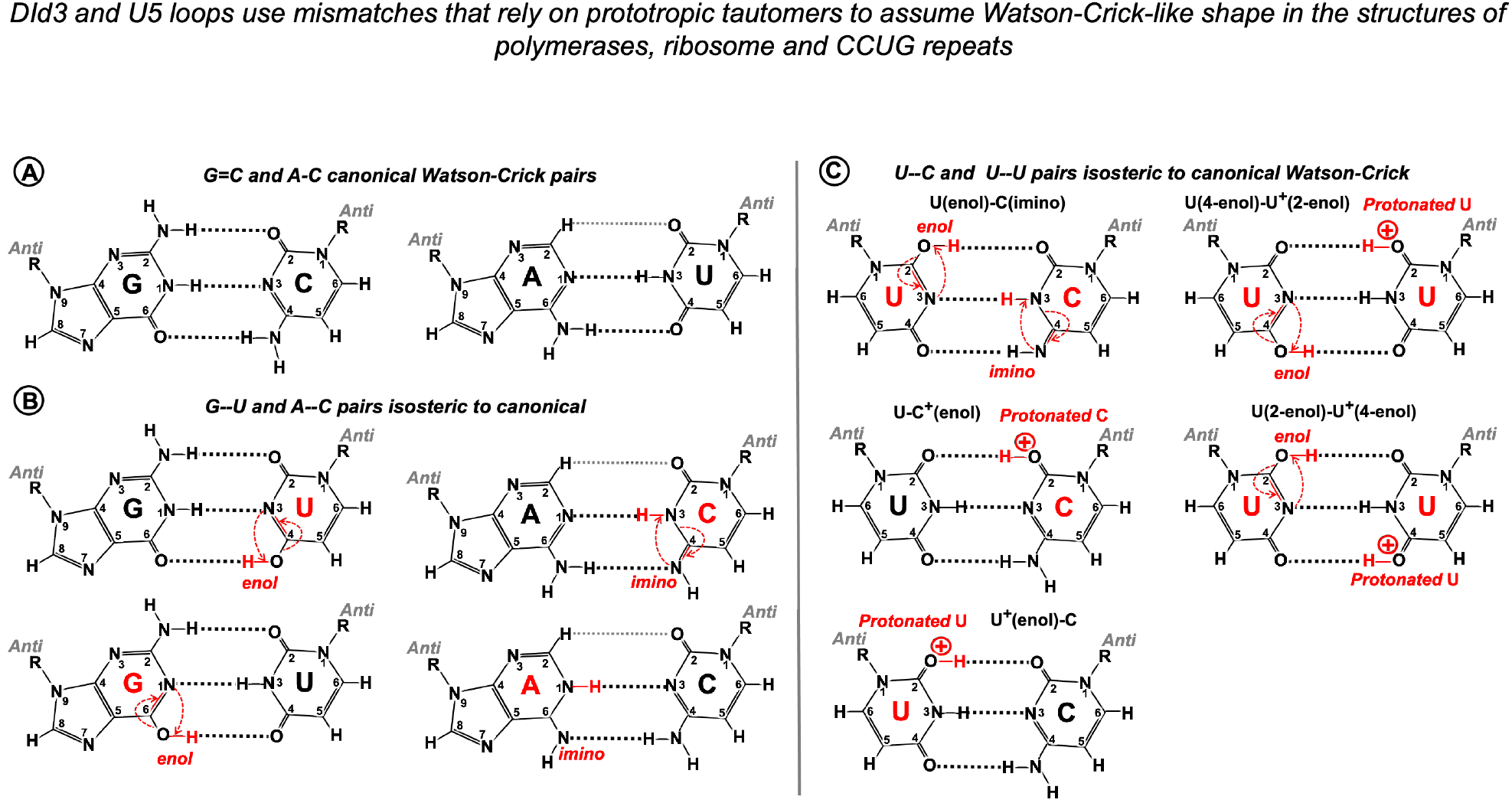
Watson-Crick like geometry of G--U, A--C, C--U and U--U pairs is supported by rare tautomerisation and protonation. **A:** Canonical Watson-Crick G=C and A-U pairs. **B:** Predicted in the 1950s (Watson and Crick, 1953) and confirmed in 2011 by X-ray structures (Bebenek et al., 2011; Wang et al., 2011) Watson-Crick-like (isosteric to canonical) G--U pairs with either base in *enol* configuration and A--C pairs with *imino* tautomers of adenine or cytosine. Watson-Crick like G--U is a high frequency pair (NMR, Kimsey et al., 2015), which reflects the ease of proton repositioning provoked by the oxygen of the carbonyls. **C:** Watson-Crick-like U--C and U--U pairs were reported by Rypniewski et al., 2016 in XR structures of CCUG repeats, associated with molecular pathology of myotonic dystrophy type 2. Possible configurations of U--C pairs are as in Rypniewski et al., 2016. The only configurations of the U--U pair that abolish the repulsion between the carbonyls and fit the reported structure (Rypniewski et al., 2016) is suggested here. Watson-Crick-like C--C pairs have not been reported, theoretical configuration requires imino tautomerisation of one cytosine and protonation of the other (4-imino-C)--(2-enol-C^+^). Imino tautomerisation is more difficult compared to enol, as the proton movement is between the two nitrogens.

Mismatched pairs isosteric to canonical are recently discovered in helix structures of accumulating human microsatellite expansion transcripts (reviewed in Błaszczyk et al., 2017). X-ray crystallography revealed WC-like C--U and U--U pairs stabilized by tautomerism or protonation (**Figure 11C**) in crystal structures of CCUG repeats associated with molecular pathology of myotonic dystrophy type 2 (DM2, Rypniewski et al., 2016).

NMR analysis of synthetic RNA and DNA duplexes provided exciting evidence that G•U and G•T wobbles exist in dynamic equilibrium with short-lived WC-like G--U and G--T pairs, stabilized by tautomerisation (one of the bases adopting a rare enol configuration) or ionization (one of the bases in anionic form) – Kimsey et al., 2015. The authors estimate that these rare tautomeric and anionic nucleobases occur with probabilities 10^-3^-10^-5^ and imply the universal role of WC-like mispairs in routine cellular processes.

Here we suggest that mimic pairs are routinely implicated in pre-mRNA splicing and Group IIA intron mobility. U5 and DId3 loops recognize their variable target sequences by helix architecture, accepting Watson-Crick and isosteric base pairs and rejecting geometrically different pairs, which perturb the helix architecture and make it unstable or incompatible with the spatial restrictions of the catalytic core. Although statistical testing cannot provide a direct prove of this second point of the U5 hypothesis, analysis of two thousand human exon junctions shows that the base pairs that cannot support Watson-Crick geometry by prototropic tautomerisation stay under 15% in the interactions of U5 and the exons, which means that on average there is only one such geometrically awkward pair per exon junction. Moreover, non-isosteric pairs are exceptionally rare in the 5’exon position -1, and cannot occur in positions -2 to -5, as these pair with U5 uracils, that are capable to form isosteric pairs with any other base, so the 5’ exon end normally has a perfect helix of at least 5bp. In the absence of +5G in the following intron significantly more Watson-Crick pairs are observed in these positions in place of isosteric pairs – statistical analysis discussed below. 3’ exon also very rarely has non-isosteric pairs in position +1, and in the absence of the conserved -3C in the upstream intron, there are significantly more Watson-Crick pairs in the exon position +1. Thus, generally geometrically awkward pairs occur in distal positions, and the shape of the U5 helix at the splice junction is preserved by isosteric pairs with prototropic tautomers, which allows for exon sequence diversity.

Finally, as we propose a mechanism that implies tautomerisation of RNA bases, we remark that the predominant tautomers in RNA are a general convention for ‘physiological conditions’, rather than a fact supported by evidence for the discussed U5 interactions with the exons in the spliceosomal ribozyme core. Tautomer diversity is often at the basis of RNA catalysis and ligand recognition, as demonstrated by structural studies of ribozymes, RNA aptamers and riboswitches (reviewed in Singh et al., 2015).

### 3.4 Statistical testing of the new model of the interactions of U5 snRNA with human exon junctions

We took advantage of our pilot investigation of the human dystrophin gene to plan our statistical analyses and looked specifically at the interactions of exons with U5 Loop1 linked to the introns that lack conserved positions +5G at the start and -3C at the end. We also took care to distinguish between U1 and U5 interactions with the 5’ exon, paying attention to the role of distinct positions. We generated datasets of U5 and U6 base pairs in the interactions of 2000 human splice junctions and their introns and analysed these datasets for corelated base pair variation at specific position. Symmetrised Kullback-Leibler (sKL) divergence shows, that +5G substitutions in the introns are associated with changes in the distribution of U5 base pairs with the 5’ exon, but not with the 3’ exon. While sKL divergence is the largest for 5’exon positions -1 and -2, which also pair with U1 snRNA in the early spliceosome, there is some divergence for 5’ exon positions -3 to -5. U1 does not bind positions -4 and -5 and selects a different base in position -3. Therefore, we observe the change in the distribution of U5 base pairs. Reciprocally, sKL divergence indicates that exon-end G substitutions are linked to changes in the distribution of U6 base pairs in the following intron positions +5 to +8. Divergence is largest for positions +5 and +6, there is some divergence for position +7 and +8, while positions +9 and +10 do not show a change in base pair distributions. We then enhanced the resolution of our analysis by bootstrap resampling of each U5 base pair frequency at each individual position of the exon junction and calculated bootstrap differences between splice junctions linked to introns that either carry +5G substitutions or conserve +5G. We found a significant increase of U5 Watson-Crick pairs with 5’ exon positions -1, -2, -3 and -5. Positions -3 and -5 indicate that this effect is specific to U5, rather than U1 snRNA. For the sake of comparison, we re-aligned the 5’ exon with U5 Loop1 according to the most recent Cryo-EM model, which means the loss of Watson-Crick pairs in positions -1 and -5. The lack of U6 C_42_=G_+5_ pair is compensated only by the increase of A-U pairs in 5’ exon positions -2 and -3 without the superior energy benefit of the G=C pair in exon position -1, which is an argument in favour of our new model (compare **Figure 12A and B**).

**Figure 12.**
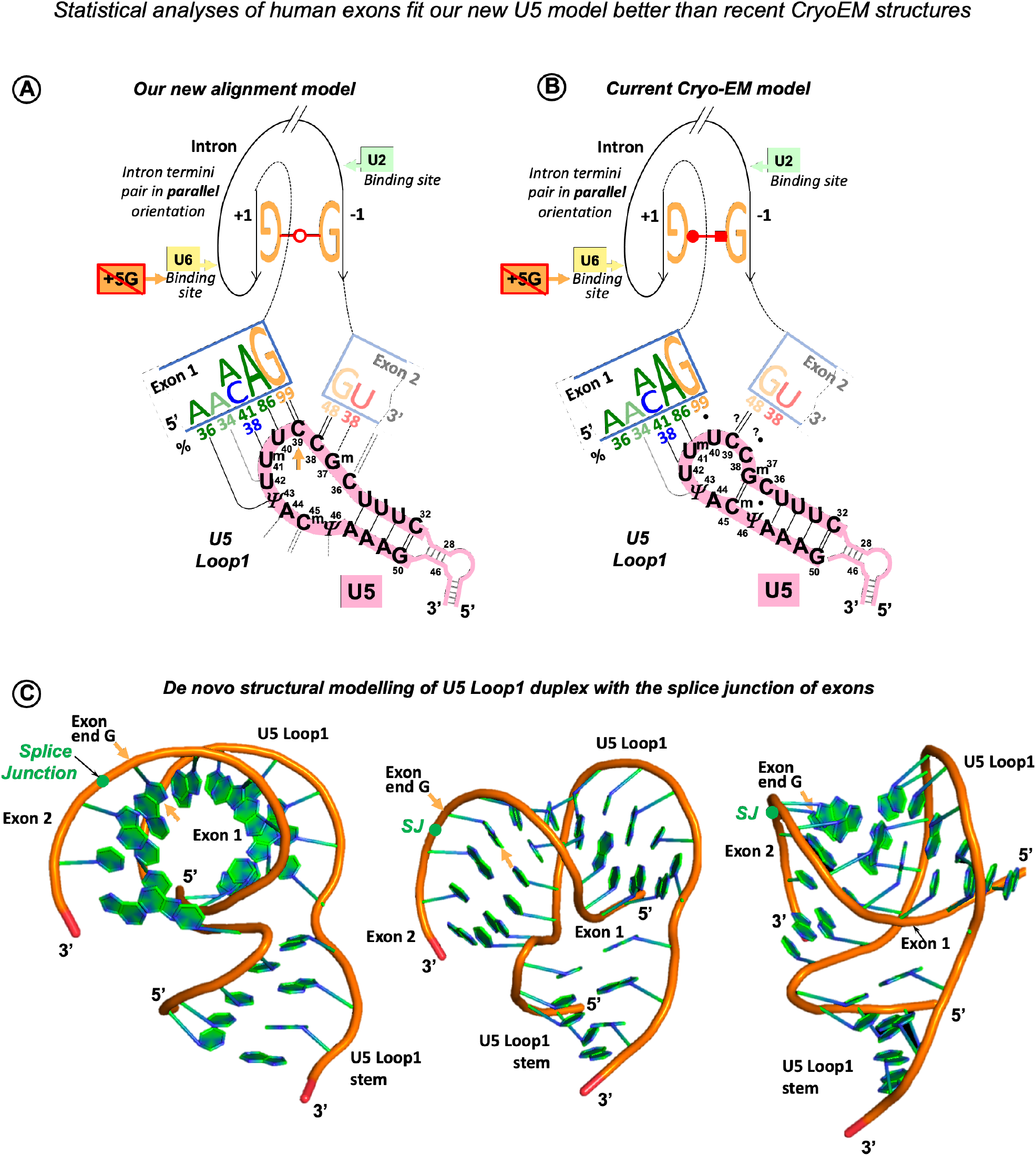
Our new U5 Loop1 interactions model compared to the current Cryo-EM model. The exon junction logo of the +5Gsub group (frequencies as in Figure 4B) reflects that substitutions of the conserved +5G in human introns are associated with significant increase of -5A, -3A, -2A and -1G in the 5’exon (Exon 1) and no changes in the sequence of the 3’exon (Exon 2) – compare to the exon junction logo for all human introns in Figure 1F. Here we fit this +5Gsub exon junction logo alternatively to our new U5 model and the current CryoEM model and argue that the new model is a better match. **A.** Our U5 Loop1 model is based on the initial alignment of human splice junctions with U5 Loop1 in parallel with the alignment of bacterial retrotransposition sites with homologous *Ll.*LtrB Id3 loop. According to our model substitutions of the conserved +5G in human introns are compensated by the additional Watson-Crick pairs with U5 Loop1 in the 5’ exon positions -1, -2, -3 and -5. In addition, our model explains the effect of mutations of exon-start G (Fu et al., 2011) by Watson-Crick base-pairing of exon positions +2 and +3 with U5 Loop1 C_36_G^m^_37_ (Figure 9). The intron termini pair is shown in the configuration of the 2^nd^ Westhof geometric family in agreement with the previous mutation analyses (Scadden and Smith 1995). This pair must be formed in the pre-catalytic spliceosome (complex B) to play a central role at the transition stage (complex C*). The intron termini pair brings the 3’ Exon 2 in contact with U5 Loop1 in the pre-catalytic spliceosome (Discussion **3.6**). **B.** The CryoEM model for U5 that currently prevails features a 7nt Loop1 and places the 5’ exon paired with U5 U_40_U^m^_41_U_42_ in the pre-catalytic complex B (Zhang et al, 2017, 2018, 2019). This eliminates the energy benefit of the G=C pair for the 82% conserved intron-end G. Accordingly, +5G substitutions are only supported by the increase of A-U pairs in exon positions +2 and +3. The intron termini pair was captured only in the post-catalytic spliceosome (complex P, Zhang et al., 2017), although the authors suggest that it must be present at the transition stage (complex C*). The configuration of this pair corresponds to the 3^rd^ Westhof geometric family, which is not consistent with the previous mutation analysis, as the covariant A••C pair or compensatory A••A and I••I pairs) are impossible in this configuration (Scadden and Smith 1995, Discussion **3.6.2**). Base-pairing for the 3’ exon is still not resolved (Zhang et al., 2019). We placed 3’ exon aligned with only two possible unpaired positions of the U5 Loop1 (base-pairing with question marks). However, exon +2C or +3G cannot form Watson-Crick pairs with U5 Loop1 in this binding register, so the fact that exon +2C/+3G promote inclusion of exons with +1G mutations cannot be explained by the CryoEM model. The 7nt U5 Loop1 is too small to accommodate specific interactions with both exons. **C.** Our *de novo* structural model of U5 Loop1 duplex with the splice junction of exons. We used hypothetical exons complementary to U5 Loop1, while our comparison with *Ll.*LtrB Id3 loop suggests that the real exon junctions form Watson-Crick-like pairs to fit diverse sequences and preserve the shape of the helix. Remarkably, a turn of the A-helix contains 11bp, so 11nt loop as shown here can well accommodate specific interactions with both exons. The U5 Loop1 helix appears hollow along the axis, which is typical of the A-helix (Heinemann and Roske, 2020).

Further, bootstrapping U6 base pair frequency at intron positions +5 to +8 shows significant increase of Watson-Crick pairs linked to substitutions of exon-end G. We conclude that U5 and U6 snRNAs collectively ensure precise definition of the exon-intron boundary and mutually compensate for their variable splice sites by Watson-Crick base pairing to stabilise the pre-catalytic complex.

We continued to examine the 3’ intron/exon boundary by bootstrap resampling of U5 base pair frequency at each individual position of the splice junction and found a significant increase of U5 Watson-Crick pairs with 3’ exon-start position +1 linked to the introns with substitutions of conserved -3C. This result shows that the U5 interaction with the 3’ exon is important for splicing fidelity.

### 3.5 Splicing effect of human exonic mutations can be explained by our U5 interactions model

The effect of mutations of the 82% conserved exon-end guanine are currently explained by base pairing with U1 snRNA in the early spliceosome complex. However, our new U5 model is the first clear base pairing scheme for the exon-start guanine, that is 50% conserved in humans. We seek to connect our U5 interactions model with the real mutation data. Fu et al., 2011 examined the effect of 14 exon-start G mutations on exon inclusion using minigene constructs in cultured human cells and report highly variable PSI ranging from 0 to 100%. The authors suggested that the length of the polypyrimidine stretch is an explanation for the observed variation. However, mutant exons with 100% PSI were included persistently at 59-83% efficiency even if most of these pyrimidines in the preceding intron were changed for purines. While we expect that the efficiency of exon inclusion also depends on the branchpoint sequence and conserved intron position -3, we opted to check if the exon interaction with U5 snRNA has an influence on PSI. We re-examined the exon sequences from Fu et al., 2011 according to our scheme of base pairing with U5 snRNA and found that the presence of our predicted Watson-Crick pairs at exon positions +2 and +3 emerge as a strong factor that promotes exon inclusion in spite of +1G mutations (**Figure 9A**). This real mutation data has two important implications: it supports our proposed binding register of the exon junction and U5 snRNA and requires fully open 11nt U5 Loop1 for compensatory pairs with +2C and +3G (**Figures 9C**, **12A**). On the contrary, we cannot explain this mutation data according to the recent CryoEM reconstruction of U5 Loop1 (Zhang et al., 2019). Base pairing with the exon-start (or 3’exon, Exon 2) has not been resolved yet, however, if we align the exon-start to U5 C_38_C_39_ left unpaired because of the 5’exon shifted by 1 nucleotide in the 7nt loop, exon +2C or +3G cannot provide any obvious energy benefit to this structure (**Figure 12B**). We suggest that specific binding of both exons by U5 can be spatially resolved only if Loop1 extends to 11nt, as a turn of the A-helix accommodates 11bp (recently reviewed in Heinemann and Roske, 2020). To demonstrate this, we created a *de novo* structural model of U5 Loop1 using simRNAweb server (https://genesilico.pl/SimRNAweb, Magnus et al., 2016) which features a hypothetical exon junction complementary to the loop, while the real exon junctions will contain mimic Watson-Crick like pairs to preserve the geometry and the overall helix architecture (**Figure 12C**). Our challenge now is a structural model of U5 and the exons before the ligation, that should include the intron termini pair. The configuration of this universally conserved pair is specifically discussed below in section **3.6.2**.

### 3.6 The new U5 model implies changes to the putative RNA network in the spliceosome

Statistical analysis of human exon and intron sequences and the available human mutation data show that the interaction of U5 Loop1 with the 3’ exon is important for splicing fidelity. The effect of -3C substitutions (Results **2.2.3**) suggests that at the 3’ intron/exon boundary a similar mechanism is at work to that of the 5’exon/intron boundary. If so, what is the RNA partner for intron position -3? and When does the collective recognition by U5 and this other RNA partner occurs? Hence, we are obliged to discuss the interactions of the 3’ intron end and the timing of the 3’ exon interaction with U5.

#### 3.6.1 U2 snRNA pairs with the 3’ intron end skipping the PPT in spliceosomal complex A

In the spliceosome the branch point adenosine is distanced from the intron end by the highly variable polypyrimidine tract (PPT), a protein interface for alternative splicing regulation, which does not belong to the ribozyme core. A solution first proposed by Kent et al., 2003 (on the basis of the Fe-EDTA probing of the U2AF^65^/RNA interactions and a comparison with X-ray structures of related **<u>R</u>**NA **<u>R</u>**ecognition **<u>M</u>**otifs, RRM) is that *‘U2AF65 bends the RNA to juxtapose the branch and 3’ splice site’*. This model is in agreement with the later X-ray structure of U2AF^65^ bound to polyU, which reports 120^0^ kink in the RNA strand (Sickmier et al., 2006). Indeed, the flexibility of the RNA chain is essential for U2AF^65^ binding, as it is blocked if uracils in the PPT are converted to pseudouridines (Chen et al. 2010), which conveys rigidity to the sugar-phosphate backbone (Charette and Gray, 2000). In fact, U2 snRNA binds the branch point site only after U2AF^65^ appropriately shapes the PPT. It can be explained if we imagine that U2 bridges across the looped out PPT and pairs with the end of the intron. Crucially, human mutation analysis indicates the involvement of the conserved intron -3C in the branchpoint helix. Corrionero et al., 2011 explored the underlying mechanism of splicing failure caused by -3C substitutions in intron 5 of the *Fas/CD95* gene (a jammed apoptotic receptor switch in T-cells leads to **<u>A</u>**utoimmune **<u>L</u>**ympho**<u>p</u>**roliferative **<u>S</u>**yndrome, ALPS). They demonstrated that while the U2AF^65^ binding efficiency is not affected, substitutions of -3C block the U2 snRNA binding. The nucleotide distribution at position -3 in human introns (**Figure 1A**) is consistent with covariation of C_-3_=G/ U_-3_--G pairs, which points at U2 G_31_ as the only possible partner base (**Figure 10A**), doubled by U12 G_16_ in the minor spliceosome, a paralogous complex that processes 0.4% of human introns (**Figure 10C**). The question remains if the invariant A-_2_ can pair with U2 A^m,m6^ (2’O-methyl,N6-methyladenosine). This adenine modification suggests a sugar edge interaction with the Hoogsteen edge of A_-2_ (similar to U6 A^m6^ ••A_+4_ pair in **Figure 1D**) However, Hoogstein edge of A_-2_ interacts with BP A according to CryoEM (Wilkinson et al., 2020). In the minor spliceosome, that has a lot fewer base modifications in snRNAs, U12 U_15_-A_-2_ pair is a perfect match (**Figure 10A,C**). Regardless of possible A_-2_ partners, the proposed U2 G_31_=C_-3_ pair explains the need for the protein co-factors SF1 and U2AF^65,35^ to bind the branchpoint, polypyrimidine tract and the 3’ intron end before the U2 RNA component: PPT needs to be looped out to allow the U2 snRNA to bridge across it to the end of the intron. Extending the BP helix beyond the variable PPT brings the bulged adenosine at a fixed distance of 4nt from the intron termini pair. Accordingly, in Group IIA introns 4bp is a conserved distance between the BP A and the pair formed by the first nucleotide of the intron and the sub-ultimate base (bacterial *Ll*.LtrB intron - **Figure 10B**, Group IIA introns in general - Zymmerly Lab Group II intron database http://webapps2.ucalgary.ca/~groupii/ Candales et al., 2012).

#### 3.6.2 Intron termini pair is formed and the 3’ exon binds U5 Loop1 in the pre-catalytic spliceosome (complex B)

The principal interaction between the bases at the intron termini provides the necessary structural link for the transition between the two catalytic steps of splicing. This interaction is universally conserved in all Group II and pre-mRNA introns and involves non-Watson-Crick base pairing (Chanfreau and Jacquier, 1993; Parker and Siliciano, 1993; Chanfreau et al., 1994; Scadden and Smith 1995). In eukaryotic introns the first and last guanines form such a pair, however, human introns occasionally accommodate A••C in place of G••G (**Table 1B)**. Compensatory double mutation analysis showed that it is also true for *S. cerevisiae* introns (G••G can be exclusively substituted for A••C Parker and Siliciano, 1993; Chanfreau et al., 1994). Scadden and Smith, 1995 explored the exact configuration of the intron termini pair in mammalian introns and showed that substitution of guanines for inosines does not affect the pair formation. Inosine is a guanine analogue that lacks N2-amino group which means that −*NH*_2_ hydrogen bonds are not involved in the pair configuration. In addition, it appears that A••A also weakly supports splicing. The predicted configuration that does not involve N2-amino groups of guanines and allows G••G to be exchanged for A••C and A••A involves H-bonds between Watson-Crick edges with *trans* orientation of glycosidic bonds and parallel sugar-phosphate backbone orientation (**Figure 13A,** explanatory **Figure S3)** as opposed to *cis* glycosidic bonds orientation of the canonical Watson-Crick pairs with antiparallel strands orientation (**Figure S4**). CryoEM studies (Bai et al., 2017, reviewed in Wilkinson et al., 2020) differ from the configuration predicted by mutation analyses as Watson-Crick edge of intron G_+1_ appears to form H-bonds with the Hoogsteen edge of intron G_-1_ (with glycosidic bonds in *cis* orientation and parallel strands). This is problematic, as this configuration involves N2-amino group of G_+1_ and A••C or A••A do not exist in this configuration (3^rd^ geometric family according to Westhof classification - on line RNA base pair catalogue http://ndbserver.rutgers.edu/ndbmodule/services/BPCatalog/bpCatalog.html). Group II intron ends are joined by base pairing of the first and sub-ultimate nucleotides. This is often the G••A pair featured in diverse introns of IIA, IIB and IIC subclasses (**Table 1A**) and the configuration of this pair was captured by X-ray crystallography (Costa et al., 2016, **Figure 13**).

**Figure 13.**
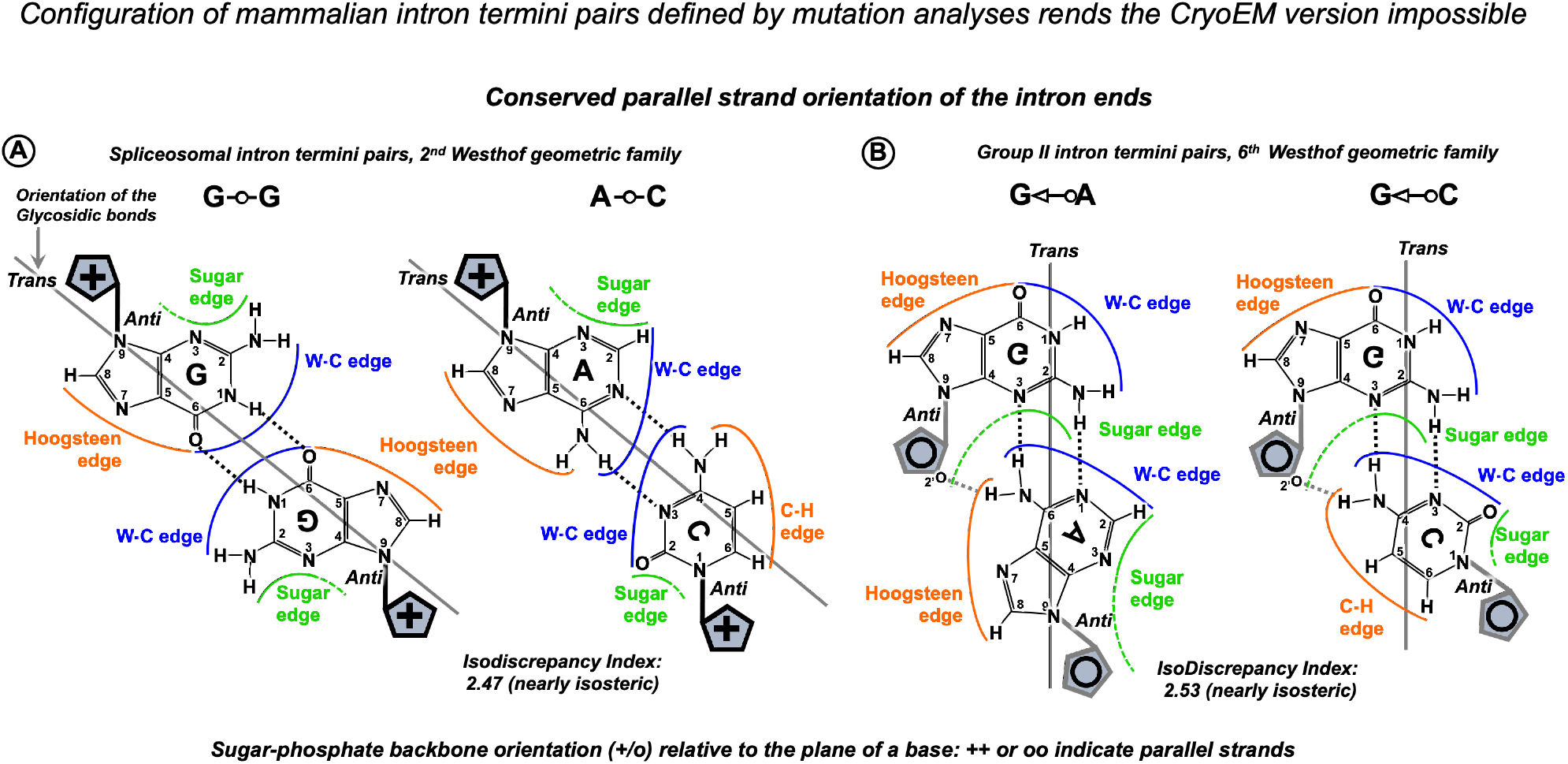
Interactions of the intron termini are base pairs with parallel strand orientation. **A**: The configuration of mammalian intron termini pairs was defined by mutation analyses (Scadden and Smith, 1995) as G••G: N1, carbonyl symmetric, A••C: reverse wobble, both correspond to the 2^nd^ Westhof geometric family. CryoEM configuration corresponds to the 3^rd^ geometric family, which is impossible for A••C pairs (for further explanation see text) **B**: Base pair configuration for Group II intron first and sub-ultimate nucleotides captured in the recent crystal structure (after Costa et al. 2016, confirmed by personal communication with Professor Eric Westhof) and shown here with additional hydrogen bonds formed by 2’O of the riboses (after Leontis et al. 2002) Parallel strand orientation is characteristic of these pairs. Ribose is located on the perpendicular plane and is shown as a schematic blue pentagon with **+** and **o** indicating the opposite directions of the sugar-phosphate backbone. For further explanation see **Figure S3**. IsoDiscrepancy index is a numerical measure of geometric similarity (isostericity) of base pairs (on-line RNA base pair catalogue http://ndbserver.rutgers.edu/ndbmodule/services/BPCatalog/bpCatalog.html).

Although the position of the Group II intron termini pair is shifted to sub-ultimate nucleotide and the pair itself is different from eukaryotic introns, the evolutionary conserved feature is the parallel strand orientation of the intron ends. Plausibly, this conformation brings the exons together and supports splice junction binding to U5 Loop1 (or DId3 loop). Certainly, the quintessential intron termini pair is central for the two-step splicing mechanism, as it provides a structural link necessary for the transition between the intron branching and the exon ligation. Once the intron breaks off from the 5’ exon and the branch point helix rotates on its’ axis (Somarowthu et al., 2014; Bertram et al., 2017a), the 3’ exon is towed into the reactant site by the intron termini pair (**Figure 14**). In order to be functional at the transition stage, this link must be formed at the pre-catalytic stage.

**Figure 14.**
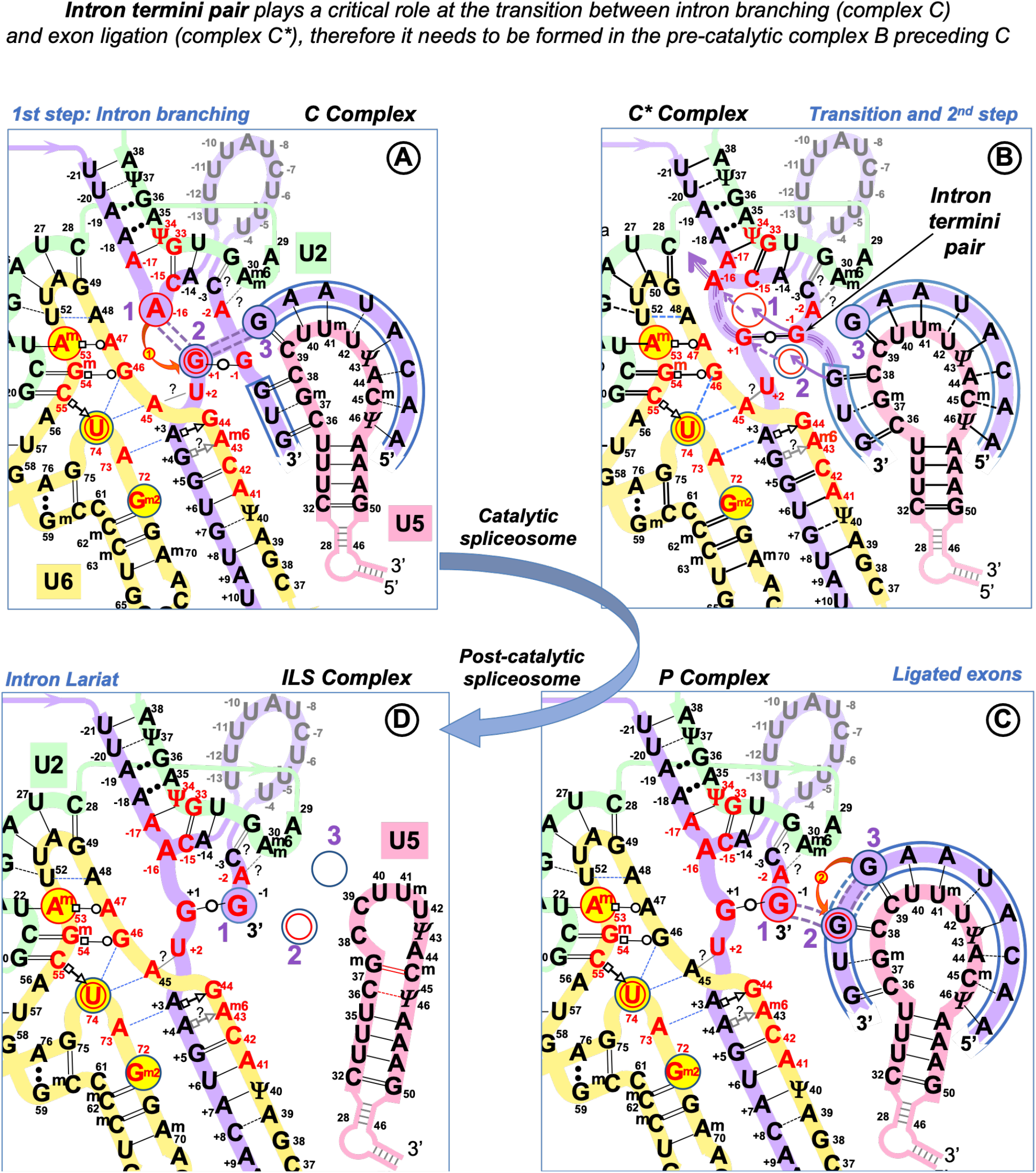
The central role of the intron termini pair in the transition between the two catalytic steps of splicing. The bond between the intron and the 5’exon is broken at the first catalytic step and the formation of the covalent bond between the BP adenosine and the 5’ end of the intron (branching) triggers a rotation of the BP helix on it’s axis (Somarowthu et al. 2014; Bertram et al., 2017a). The correct repositioning of the branched intermediate absolutely requires U6 non-Watson-Crick pair(s) at the start of the intron at position +3 and/or +4 (Konarska et al., 2006). The momentum of the revolving BP helix is transmitted by the intron termini pair and drives the translocation of the 3’ exon. This movement is enabled by the relaxation of the U5 Loop 1 due to the disruption of the covalent bond between the intron and the 5’ exon. The overall configuration of the U6/U2 metal binding site stays unchanged and the reacting residues are transitioned to the fixed <u>Reactant sites</u> (Steitz and Steitz, 1993; Fica et al., 2013; Semlow et al. 2016 – **Comment S2**). **A:** RNA network of the 1^st^ step spliceosome - as in **Figure 10. B**: The RNA-RNA interactions at the moment of transition between the two steps of splicing: The biggest purple arrow indicates the repositioning (rotation) of the BP adenosine after branching and the purple triple dashed line shows the transmission of the motion via the intron termini pair to the 3’ exon. Dashed purple arrows trace the movement of the residues out of the <u>Reactant Sites 1 and 2</u> and the in-coming nucleotides follow the path of continuous purple arrows. The position of the 5’ exon is unchanged in the <u>Reactant Site 3</u>. For the second reaction the metal ions reverse their actions: the Mg^2+^ (2) activates the 3’OH group of the 5’ exon in <u>Reactant Site 3</u>, an attack is launched at the 5’PO_4_ of the 3’ exon in <u>Reactant Site 2</u>, while Mg^2+^ (1) stabilises the leaving 3’OH of the last nucleotide of the intron at <u>Reactant Site 1</u>. The fact that the two metal ions play the activation role in turns enables a single catalytic core to accommodate both steps of splicing and also implies the ease of the reverse process (Fica et al. 2013). Although the second reaction (*curved red arrow 2*) happens in C* complex following the transition, here the emergent covalent bond is shown in the successive P complex. **C:** RNA network after the second catalytic step. Double purple dashed lines indicate the emergent (purple fill) and previous (no fill) covalent bonds. The exons are ligated and are still paired with U5 Loop1. The intron lariat stays paired with U6 and U2 snRNAs (Znang et al., 2019). **D:** The joined exons are disassociated from U5 Loop1 by Prp22 (Wan et al., 2017; Znang et al., 2019); without RNA-partners the loop changes to ‘closed’ (7nt) conformation. The intron lariat stays paired to U2/U6 and associated with U5 snRNP. ILS complex is homologous to Group II intron RNP ready for reverse splicing. (RNA network dynamics of successive spliceosomal complexes is summarised in **Table S1.**)

The formation of the intron termini pair and the earlier involvement of the intron end -3C in the branchpoint helix guarantees the proximity of the 3’ exon to U5 at the pre-catalytic stage. In fact, 3’ exon pairing with U5 Loop1 can be responsible for the transformation of the Loop from the closed 7nt conformation, which it is likely to adopt without RNA partners, to the fully open 11nt form, that can also accommodate the extended 5’ exon helix.

#### 3.6.3 Overall arrangement of pre-mRNA before the catalytic activation of the spliceosome

The intron termini pair and the 3’ exon pairing with U5 in the pre-catalytic spliceosome imply that all the core snRNA interactions with pre-mRNA splice sites are already formed prior to the remodelling by *Brr2* and configuration of the catalytic Mg^2+^ binding site. Collective recognition by snRNAs is the only way to maintain splicing fidelity in the context of a complex genome with divergent exon sequences and variable introns. Our model of U5 interactions with the exons explains that at the 5’ splice site U6 and U5 mutually compensate for the loss of complementarity at their binding sites and similarly at the 3’ splice site for U2 and U5. We propose that the overall threshold stability of all the recognition helices between the substrate and U6, U5 and U2 snRNAs in the pre-catalytic complex B is a fidelity checkpoint for spliceosome activation. (Successive spliceosome complexes detailed in **Table S1.**)

### 3.7 The new U5 model agrees with increasing exon sequence diversity during protein evolution

The evidence of alternative conservation of the intron and exon consensus in higher eukaryotes was presented previously as the evolutionary migration of the splicing signals from exons to introns. Indeed, molecular evolutionists had long identified that ‘old’ introns have a conserved intron consensus and ‘new’ introns on the contrary a conserved exon consensus (Sverdlov et al., 2003). In view of the U5 hypothesis, gradual replacement of Watson-Crick pairs with isosteric mismatches has provided more diversity of the U5 binding sites, relaxing constrains for the sequence of the exons and aiding protein evolution. This process was supported by the conservation of specific Watson-Crick pairs at the U6 and U2 binding sites in the introns, ensuring preservation of the impeccable splicing fidelity.

### 3.8 The need of mutation analyses of the U5 interactions with human exons and the proposed U2 interaction with the 3’ intron end

Experimental validation of the U5 hypothesis requires checking base pairing by double mutagenesis, which aims to introduce covariant pairs between the interacting RNAs. A Watson-Crick pair is geometrically interchangeable for another Watson-Crick pair, however if we account for the energy benefit it is better to swap G and C between the interacting RNAs, rather than introduce an A-U pair. We do not know if changes in the U5 sequence will affect the conformation of Loop 1 or cause a shift in the binding register. Presumably, diverse sequences of Id3 loops in Group IIA introns (**Figure S1A**) follow the same spatial scheme. As a precaution, while subjecting one exon interaction with U5 to double mutagenesis it might be safer to choose another exon complementary to U5 Loop1 to secure the position of the exon junction.

#### 3’exon and U5 Loop1

It is best to start with the 3’ exon binding register to avoid the ambiguity of U1 binding at the 5’ splice site. As we observed that exon +2C/+3G promote inclusion of exons with +1G mutations, we suggest changing nucleotides at exon positions +2 and +3 in minigenes from the study of Fu et al., 2011. For example (**Figure 9B**): will introducing +2C and +3G into exon 25 of *PKHD* gene and removal of +2C, +3G at the cryptic 3’ss suppress the effect of the +1G → *T* mutation and re-activate the normal 3’ss? Further to verify base-pairing changes of exon position +2 and +3 in the minigenes can be combined with mutations at positions 36 and 37 of Loop1 in a U5 expression construct and followed by co-transfection and quantification of the splicing outcome in human cells.

#### 5’exon and U5 Loop1

Mutation analyses for the 5’ exon interaction with U5 is complicated by the initial U1 interaction across the exon/intron boundary. Presumably, nucleotide changes at the exon-end will not block U1 binding if we make sure that complementarity to U1 extends over 5-6 base pairs overall (Ketterling et al., 1999). The second complication is that the 5’ exon binds a stretch of 4 uridines of the U5 loop1, making any 4 nucleotides acceptable at exon positions +2 to +5 as uridine is prone to form isosteric pairs and there is a mechanism of compensation by U6 snRNA for the poor U5 binding affinity. The safest strategy will be to start with the most conserved exon-end G and its partner U5 39C and swap these nucleotides between the interacting RNAs. We include examples of suitable human mutations for the proof-of-principle laboratory testing, some of which are the continuations of the previous studies introduced and discussed above: Juan-Mateu et al., 2013 **Figure S5**; Vincent et al., 2013 **Figure S6;** Carmel et al., 2004; Scalet et al., 2017; Scalet et al., 2018; Breuel et al., 2019 (Experimental designs for these examples are described in supplementary **Section S5**).

#### Intron -3C and U2 snRNA

We suggest that U2 snRNA interacts with the 3’end of the intron: U2 G_31_=C_-3_ (discussed in **3.6.1**). Testing this U2 pair is more straightforward and any human intron in a minigene construct can do, however the *Fas/CD95* intron 5 (Corrionero et al., 2011) is an excellent study to follow by swapping the proposed U2 G_31_=C_-3_ for a double mutant U2 C_31_=G_-3_ (**Figure S7;** described in **Section S5**).

### 3.9 The incentive: re-targeted spliceosomes for therapeutic applications

#### 3.9.1 Small nuclear RNAs targeting splicing mutations

Suppression of splicing mutation by matching modifications of U1 snRNA was discovered by Zuang and Weiner in 1986. In 1989 the same authors demonstrated that modifications of U2 snRNA to increase complementary to the target branch point site in the human β-globin gene was able to suppress a mutation, that created a cryptic 3’ss. In 1996 Hwang and Cohen followed with suppression of the 5’ splice sites mutations by compensatory changes in U6 snRNA, that increased complementarity to intron positions +5 to +9. However, while U5 snRNA with modified Loop1 sequence was shown to promote the use of cryptic splice sites, the base pairing model involved 5’ intron end, which unfortunately jumbled up experimental planning and conclusions (Cortes et al., 1993). The important outcome of these early studies is that exogenous modified snRNAs, that still have their protein-binding sites unchanged, are recognized by spliceosomal protein components and undergo normal assembly process to form functional ribonucleoproteins (spliceosomal snRNPs).

In spite of these previous studies on specific recognition of pre-mRNA by U2, U5 and U6, today snRNA therapeutics is largely limited to U1-based approach (the recent studies include: Balestra et al., 2020; Breuel et al., 2019; Yamazaki et al., 2018; Scalet et al., 2017). The efficiency of splicing correction is variable depending on individual mutations and gene context and a combination of adapted U1 snRNA and antisense oligos, that block cryptic slice sites, is often used to increase the ratio of normal to aberrant products (recent examples include: Breuel et al., 2019; Lee et al., 2019; Balestra et al., 2015). Encouragingly, modified U1 snRNAs proved to be safe *in vivo* (Balestra et al., 2020; Donadon et al., 2019; Lee et al., 2016), possibly because of the competition with the endogenous wt U1 snRNA and due to nonsense mediated decay mechanism that removes any jumbled transcripts of off-target genes.

U6 snRNA modification was again attempted by Carmel et al., 2004. The authors used alternatively U1 and U6 adapted to match a substitution of +6T and achieved partial splicing correction of the human *IKBKAP* gene with U1, but not with U6 snRNA. More recently, Schmid et al., 2013 demonstrated that a combination of modified U1 and U6 snRNAs targeting a substitution of +5G in human cells was more effective than U1 alone to rescue splicing of the BBS1 gene (<u>B</u>ardet-<u>B</u>iedl <u>S</u>yndrome, a ciliopathy associated with severe vision loss in children).

Scalet et al., 2018 provide experimental evidence of the endogenous U5 supporting modified U1 snRNA to achieve correction of aberrant splicing of the *FAH* gene (encodes an enzyme of the tyrosine I catabolism; *FAH* deficiency, **<u>H</u>**ereditary **<u>T</u>**yrosinemia type **<u>I</u>**, HTI is associated with cirrhosis and hepatocellular carcinoma). U1 snRNA modified to be fully complementary to the mutant exon/intron boundary CCG/gtga**<u>a</u>**t (the frequent *FAH*c1062+5G>A mutation in intron 12) failed to rescue normal splicing. However, a compensatory effect of a second mutation at the end of exon 12 -2C>A was discovered in a patient with somatic mosaicism and the *FAH* enzyme present in the liver. Expression of a minigene construct bearing both mutations at the exon/intron boundary C**<u>A</u>**G/gtga**<u>a</u>**t in HepG2 cells produced predominantly aberrant splicing products. However, addition of the U1 complementary to CCG/gtga**<u>a</u>**t yielded mostly correct splicing product, although U1 was not complementary to the A_-2_ change. This effect points at the improved U5 pairing in the pre-catalytic complex, which succeeds U1 snRNA binding in the early complex.

The U5 hypothesis provides the binding register for U5 modification to match the target exon junction and the proposed U2 interaction with the 3’ intron end completes the base pairing scheme for small nuclear RNAs and pre-mRNA splice sites. We propose that re-targeting all snRNA, rather than just U1, as is the current practice, will produce a spliceosome with very high affinity to the target intron and splice junction. We can further limit intermixing with endogenous snRNPs by swapping the strands of helix II between the designer U6 and U2 snRNAs. The use of such designer spliceosome with a full set of modified snRNAs will aid both efficiency and precision. Safe transient delivery of small nuclear RNA molecules (human U1, 164nt; U2, 191nt; U5, 116nt; U6, 107nt), rather than expression constructs, is facilitated by the fact, that their maturation involves a cytoplasmic stage, after which they are transported back to the nucleus (Becker et al., 2019).

#### 3.9.2 Future adaptation of snRNAs to manipulate regulatory alternative splicing switches

Importantly, applications of snRNA therapeutics are not limited to the correction of individual splicing mutations. Targeting alternative splicing switches can be beneficial for patients with common conditions, such as thrombosis. Hemostasis regulation by alternative splicing of coagulation factor V (Vincent et al., 2013, detailed in **Section S5**) is but one example. Such isoform switches are often at the crux of cell fate regulation, providing many clinically important splicing targets. Alternative splicing of the *Fas/CD95* receptor is another example: inclusion of an alternative exon changes a cytoplasmic anti-apoptotic isoform into transmembrane death receptor (Corrionero et al, 2011). Thus, promoting splicing of a pro-apoptotic isoform by a full set of complementary snRNAs can help to develop tumor suppressor drugs.

#### 3.9.3 Future gene repair by reverse splicing

The future for safe genome engineering eliminating the dangers of bacterial endonucleases will be adapting the human U5 snRNA for correction of genomic mutations by specific reverse splicing. Indeed, human snRNAs form a ribozyme identical to that of mobile introns, that are routinely used for genetic engineering in bacteria (Karberg et al., 2001; Mohr et al., 2013). Reverse splicing was previously demonstrated for the spliceosome *in vitro* (Tseng and Cheng, 2008). It is also known that Group II introns reverse splice into DNA and RNA with comparable efficiency, indicating that 2’O of the target does not affect the reverse splicing process (Griffin et al., 1995). The challenges ahead include increasing the U5 snRNA target recognition specificity and exploring the reverse splicing pathway. However, these are challenges worth taking, as spliceosomes are perfectly placed for endogenous gene therapy tools: highly abundant in transcription loci next to vulnerable single stranded DNA.

Development of gene repair by specific insertion is needed for the treatment of Duchenne muscular dystrophy (DMD), a sporadic X-linked fatal condition affecting 1:3500 newborn boys. It is caused predominantly by dystrophin gene deletions, that frequently arise within a region of genomic instability, a common fragile site (CFS) in human populations worldwide (Mitsui et al., 2010). The current therapeutic approach uses antisense oligos for exon skipping to restore the reading frame and slow the disease progression. U5 targeting individual deletion site and insertion of missing exons as a fused cassette can offer personalized dystrophin gene repair to cure DMD.

## 4. Materials and methods

### 4.1 Creating a dataset of base pairs between interacting RNAs *in silico*

We aimed to include approximately 2000 introns (splice junctions) from human genes ranging from well known in medical genetics practice to genes with experimentally confirmed function and expression (1 splice isoform with maximum exons per gene, details and the full list in supplementary materials, **List S1**). The sequences of all exons and all introns were downloaded from ensembl.org; we extracted specifically the 11 nt splice junctions (8nt of the 5’exon end joined to 3nt of the 3’exon start), intron starts (10nt) and intron ends (60nt); introns processed by the minor spliceosome and introns with unusual ends were identified and excluded from the data (excluded introns are detailed in supplementary materials **List S2** and **List S3).** Program splice_sites.py calls functions from U5.py (all the code available via Git from the U5_hypothesis repository https://github.com/oartem01/U5_hypothesis).

### 4.2 Symmetrised Kullback-Leibler divergence (sKL)

+5G_sKL.py detects +5G substtitutions in the major GU_AG introns and sorts exon junctions accordingly into two groups: +5Gsub and +5G (*N_+5Gsub_* =445, *N_+5G_* =1545) and computes U5 base pair distributions for each group (**Figure 4B,C**).

The 11 positions of the splice junction were divided into 4 subsites: 5’exon positions -8 to -6, -5 to - 3, -2 to -1 and 3’exon positions +1 to +3. sKL divergence between the distributions of the base pairs at each of the subsites was calculated as follows:

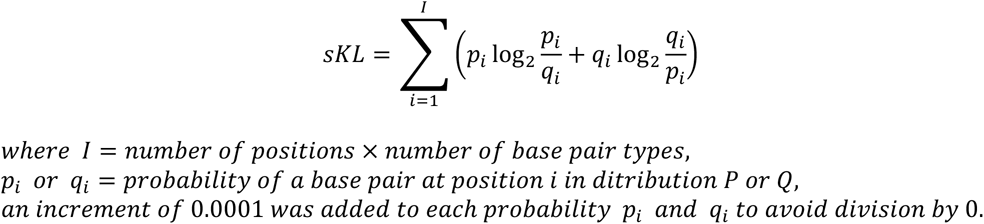

The base line control of 0 divergence is provided by comparing subsets of the larger +5G set to each other, as opposed to the divergence from the smaller +5Gsub. +5G_sKL.py generates 10000 pairs of random non-overlapping and non-redundant sets of 445 (*N_+5Gsub_)* splice junctions from the +5G set and calculates sKL divergence for each pair (+5G/+5G, control) as well as sKL divergence between one of the random +5G sets from each pair and the +5Gsub (+5G/+5Gsub). +5G_sKL.py returns histograms of sKL distributions (**Figure 4D-G**).

File exG_sKL.py performs analogous operations for the effect of the conserved exon-end G (**-1G**) on the U6 bp with the start of the intron: initially it detects exon-end G substitutions and sorts the introns accordingly: exG and exGsub (*N_exGsub_*=392, *N_exG_*=1598); and returns barcharts and histograms **Figure 7B-F**.

### 4.5 Bootstrap procedure

The Bootstrap re-samples data with replacement and enables estimates of standard error in properties of the sample that can be used to test hypotheses of difference (Efron,1979). Comparing any parameter of two datasets with a bootstrap tests the null hypothesis of no change for this parameter, returning *P*(*H*_0_) for conventional ‘statistical significance’ (McShane et al., 2019). In comparing the +5Gsub and +5G datasets we bootstrap individually the frequency difference of every base pair type at each position of the splice junction. The algorithm +5G_boots.py preforms re-sampling with replacement for both datasets: selecting splice junctions at random allowing for chance reprises and generates a re-sampled set of the same size as the original dataset (N=445 for +5Gsub and N=1545 for +5G). The program then computes the bootstrap difference (BD) of the frequency of each base pair at each position between the two re-sampled sets. This procedure is iterated 10000 times to generate a BD distribution for each type of base pairs at each position of the splice junction (**Figure S8**). The probability of the null hypothesis *P*(*H*_0_) of no difference between the +5Gsub and +5G is returned for each individual frequency. The standard definition for *P*(*H*_0_) for the bootstrap hypothesis testing is the proportion of the smaller part of the BD distribution lying beyond 0 line (**Figure 5**). Program +5G_boots.py also summarises all the histograms as 3 sets of violinplots (**Figure 5A-C**). Same program further deals separately with 5’ exon position -3 shared by U1 and U5 binding sites, but with different nucleotides required for Watson-Crick pairs with U1 and U5 snRNA. +5G_boots.py returns stacked barchart of the nucleotide frequencies (rather than base pair type frequency) for this position in +5Gsub and +5G datasets, bootstraps these frequencies and computes *P*(*H*_0_), **Figure 6**.

Script exG_boots.py compares exGsub and exG datasets (*N-_1Gsub_* =392, *N_-1G_* =1598) using the same algorithm to generate BD distribution for each individual bp frequency at each position of the U6/intron interaction, returns violinplots and P-values **(Figure 5D-F**).

File -3C_boots.py first detects substitutions of the conserved -3C in the dataset of introns processed by the major spliceosome, including GC(A)_AG introns along with GU_AG, sorts the exon junctions into two groups: -3Csub and -3C *(N_-3Csub_* =792, *N*-*_3C_* =1211) and computes U5/exons bp frequency (**Figure 8 B,C**). The program then follows the bootstrap procedure algorithm as described above and returns violinplots **Figure 8D-F**.

### 4.6 Bonferroni correction for the multiple significance tests (Dunn’s method)

The danger of testing multiple hypotheses is that some ‘significant’ result may occur by change alone (Bland and Altman, 1995). The simple Bonferroni correction or Dunn’s *α* -splitting (Lee and Lee, 2018) implies that the widely used threshold of statistical significance *α* = 0.05 must be divided by the number of tests *m* performed on each dataset

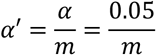

Accordingly, the corrected P-value thresholds for significant changes for base pair frequency tests here are as follows: For the +5G/+5Gsub experiment *m* = 31, *α*’ = 0.0016; for the -1G/-1Gsub experiment *m* = 17, *α*’ = 0.0029; for the -3C/-3Csub experiment *m* = 28, *α*’ = 0.0018; and considering all tests in this study *m* = 76, *α*’ = 0.0007. P-values in **Figures 5 and 8** are marked with triple asterisks or double asterisks if below their respective thresholds for all tests or individual experiments.

Dunn’s application of Bonferroni correction is a stringent method, which is more likely to reject a true positive (Type II error), than to accept a false positive (Type I error) (Lee and Lee, 2018). The application of this method is justified if the outcomes of the hypothesis tests are not related. The comparisons here are independent for the positions of the sites, but strongly correlated for base pair types at each individual position, e.g.: an increase of Watson-Crick pairs means the decrease of isosteric pairs if there are only two pair types or the decrease of either non-isosteric or isosteric pairs (or both) if there are three pair types at any given position. Therefore, we can adjust *m* for the correlated tests (Shi et al., 2012):

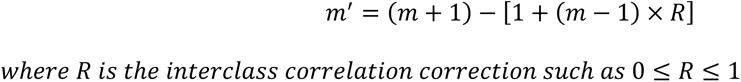

In simple terms, positions with only two pair types account for two perfectly correlated tests, so *R* = 1 and these two tests will count as one. For tests with three types, we can approximate *R* ≈ 0.5 by splitting the correlation between them and *R* ≈ 0.33 when there are four tests. Following this procedure, the +5G/+5Gsub experiment *m*’ = 20, *α*’ = 0.0025; for the -1G/-1Gsub experiment *m*’ = 11, *α*’ = 0.0045; for the -3C/-3Csub experiment *m*’ = 17, *α*’ = 0.0029. P-values below their respective experiment thresholds accounting for correlated tests are marked with a single asterisk in **Figures 5 and 8.**

Here we remark on the current debate on ‘statistical significance’ among the statisticians: McShane et al., 2019 points out that the null-hypothesis significance testing - and generally accepted P-value threshold of 0.05 – is a misleading paradigm for research and instead *P*(*H*_0_) should not be prioritised over other factors, such as plausibility of mechanism and related prior evidence (in this case genomic conservation and mutation data).

## Supporting information

U5_hypothesis_Supplemetary_material

## Author contributions

OI performed the initial sequence alignments of U5 with human exon junctions and *L.l.*LtrB DId3 with bacterial retrotransposition sites, designed the analyses of the relative distributions of base pairs by position in human exon and intron interactions, written the Python code and performed the analysis. OI is responsible for the new U5 interactions model and comparison with the current CryoEM model. OI linked human mutation data to the new U5 model, written the codes in Python and R and performed the tests. AP directed the choice of the journal, and preparation for the experimental verification by exploring the role of U5 in the alternative splicing of the human F5 gene. OI wrote the paper and prepared the figures. AP revised and corrected the manuscript and figures.

## Acknowledgements

OI is most grateful to Professor Eric Westhof for his explanation of the intron termini pair configuration. OI acknowledges the important contributions of Dr Mark Williams (ISMB, Birkbeck, University of London), which include critical review and multiple corrections of the manuscript and supervision of statistical analyses. OI would like to thank him also for conscientious teaching of statistics on the Bioinformatics course at Birkbeck. OI is deeply grateful to Dr Irilenia Nobeli for the life-changing introduction to bioinformatics, sharing her unpublished results, critical review and corrections of the manuscript and diligent teaching of sequence analysis and genomics. OI would like to thank Dr Adrian Shepherd (ISMB, Birkbeck) for his inspirational teaching, that makes coding from scratch easy and for his further help with Python related to this project. OI is most grateful to Dr Susan Brown (RVC, University of London) for the meticulous critical reading of the earlier manuscript. OI would like to thank Dr Rimma Belotserkovskaya (Gurdon Institute, University of Cambridge) for her role in the preparation of the experimental verification of the U5 hypothesis. OI and AP are most grateful to Dr Josefin Ahnström and Prof. Jim Crawley (Centre for Haematology, Imperial College London) for discussions on alternative splicing of human coagulation Factor 5. OI would like to thank Mr David Houldershaw (ISMB, Birkbeck, University of London) and Mr Ketan Kansara (Imperial College Apple Tech Bar) for their expert computer support.

