## Supplementary material for "U5 snRNA interactions with exons ensure splicing precision": U5_hypothesis_Supplemetary_material

#### Contents


##### Section S1: Tables

| | page |
|---|---|
| **Table S1.** Spliceosomal RNA network at successive stages of splicing, defined as distinct RNP complexes | 3 |
| **Table S2.** Diverse views of the U5 snRNA Loop1 interactions with exons were obtained by crosslinking experiments and Cryo electron microscopy by different research groups. | |
| **S2.A.** Budding yeast *(Saccharomyces cerevisiae)* spliceosome | 4 |
| **S2.B.** Human spliceosome with adenovirus pre-mRNA substrates | 5 |
| **Table S3**. Spliceosomal RNA-RNA interactions confirmed by genetic and biochemical studies are not always captured by Cryo electron microscopy | |
| **S3.1** Budding yeast *(Saccharomyces cerevisiae)* spliceosome | 6 |
| **S3.2** Human spliceosome built on the miniature adenovirus pre-mRNA substrate (MINX) | 7 |
| **Table S4 (See Section S4)** | |

##### Section S2: Figures

| | |
|---|---|
| **Figure S1\|** Diverse group II introns recognise their exons mainly by Watson-Crick base pairing with the Id3 loop in a common asymmetrical fashion. | 8 |
| **Figure S2\|** 'Legal' C--U pairs in the U5 interactions prompted by the initial U1 selection | 9 |
| **Figure S3\|** Quick guide to sugar-phosphate backbone orientation relative to the plane of a base | 10 |
| **Figure S4\|** Canonical Watson-Crick pairs with antiparallel strands | 10 |
| **Figure S5** (see **Section S5**) | 39 |
| **Figure S6** (see **Section S5**) | 40 |
| **Figure S7** (see **Section S5**) | 41 |
| **Figure S8** (see **Section S4**) | 45-50 |
| **Figure S9** (see **Section S4**) | 51-53 |
| **Figure S10** (see **Section S4**) | 54-59 |
| **Figure S11 (**see **Section S4)** | 21 |

##### Section S3: Comments to Figures

| | |
|---|---|
| **Comment S1** (to **Figure 12**) The odd feature of the of U12 interaction with the U6atac catalytic triad in the human minor spliceosome: a bulge or a Watson-Crick-like mimic pair? | 11 |
| **Comment S2** (to **Figure 14**) The paradox of the stalled transition in the absence of the Prp16 helicase | 11 |

##### Section S4: Results and Methods

| | |
|---|---|
| **List S1.** List of human genes included in the analysis by sKL divergence and Bootstrap | 12-14 |
| The *ensembl* isoform number is featured in the sequence file names | 14-16 |
| **Unusual introns** (identification and isolation) | 17 |
| **List S2.** Minor introns | 18 |
| **List S3.** Major introns with substitutions of +2U: GC(A)_AG introns | 18 |
| **List S4. I**ntrons missing both conserved exon-end -1G and intron +5G | 19 |
| **Figure S8\|** Histograms of bootstrap difference for U5 bp types at each position of the exon junctions between +5Gsub and +5G datasets (Violinplots of the same **Figure 6A-C**) | 45-50 |
| **Figure S9\|** Histograms of bootstrap difference for U6 bp types at the start of intron position +5 to +10 between -1Gsub and -1G datasets (Violinplots of the same **Figure 6D-F**) | 51-53 |
| **Figure S10\|** Histograms of bootstrap difference for U5 bp types at each position of the splice junction between -3Csub and -3C datasets (Violinplots of the same **Figure 9D-F**) | 54-59 |
| **Human mutation data explained by the U5 hypothesis** | 20 |
| **List S5.** Mutations of exon-start guanine: $G_{+1} \rightarrow T$ or $G_{+1} \rightarrow A$ (Fu et al., 2011) | 20 |
| **Figure S11\|** Inclusion of exon affected by +1G mutations is influenced by multiple *cis* factors | 21 |
| **Table S4. F**actors that promote efficient exon inclusion (PSI 81-100%) in spite of +1G mutations | 22 |
| Kruskal-Wallis rank sum test and Welch's ANOVA (t-test) for **+2C/+3G** | 23 |
| Welch's ANOVA and Kruskal-Wallis rank sum test for **substitute A or T** | 27 |
| Welch's ANOVA and Kruskal-Wallis rank sum test for **-3C** | 30 |
| Boxplots for PSI dependent on +2C/+3G, substitute A or T and -3C | 33 |
| Spearman's correlation for PPS length | 34 |
| Spearman's correlation for branchpoint matches | 35 |
| Scatterplots for PSI dependent on PPS length and branchpoint matches | 36 |

**Section S5: Future work: Molecular cell biology testing of the U5 hypothesis**
Human mutations suitable for the proof-of-principle laboratory testing 37
**Figure S5|** Study design for the correction of the dystrophin gene splicing mutation c9563+5G>C in intron 65 from a Becker muscular dystrophy patient reported by Juan-Mateu et al., 2013. 39
**Figure S6|** Study design targeting the alternative intron (pseudo-intron) splicing of coagulation F5. 40
**Figure S7|** Testing U2 snRNA interaction with the end of the intron (the proposed U2 $G_{31}$=$C_{-3}$ pair) following the study of Corrionero et al., 2011 on Fas/CD95 intron 5. 41

**Supplementary References** 42-44

#### Section S1: Tables

**Table S1.** Spliceosomal RNA network at successive stages of splicing, defined as distinct RNP complexes

| Spicing stage | **RNP complex**[1] | Proteins in action | **RNA network** |
|---|---|---|---|
| Initial 5'splice site (5'ss) selection | **E** Early complex | SF1 binds BP | **5'ss/U1** - binds across exon-intron boundary |
| Branch point (BP) and 3'splice site (3'ss) recognition | **A** Pre-spliceosome | **$U2AF^{65}$ binds the PPT and curves it into a sharp loop**[2] $U2AF^{35}$ binds 3'ss - the intron end AG | **BP/U2** - **BP helix bridges over the PPT bringing the bulged A to the fixed 4nt distance from the intron end**[3] |
|  | **Pre-B** |  | U4/U6•U5 tri-snRNP joins in, all snRNPs are assembled together |
| Spliceosome Assembly | **B** Pre-catalytic spliceosome | Prp28 displaces U1C and initiates 5'ss/U1 unwinding | U1 quits 5'ss **U2, U5 and U6 cooperate in specific sequence recognition to achieve splicing fidelity** **5' intron end pairs with U6-$A_{39}\Psi_{40}A_{41}C_{42}A^{m6}{}_{43}G_{44}A_{45}$ (bases forming conserved non-WC pairs are boxed)** **3' and 5' exons align on the U5-Loop 1. Intron termini form a non-canonical pair.** **Pre-mRNA is completely arranged for the catalysis** |
| Re-arrangement of the catalytic core | **Bact** Activated spliceosome | **Brr2 unwinds U4/U6 stem I helix translocating 3' to 5' along U4** NTC/NTR join in | U4 quits U6 **Formation of the ribozyme catalytic core: U2/U6 triple helix and U6 dinucleotide bulge coordinate the two catalytic $Mg^{2+}$ions.** |
| Catalytic activation | **B*** Catalytically activated spliceosome | Prp2 promotes binding of the step 1 factors Cwc25, Yju2, and Isy1 |  |
| Branching reaction | **C** Catalytic step 1 spliceosome |  | Intermediates: 5' exon cut off the intron, 3' exon with the intron lariat **Both exons stay bound to the U5 Loop 1** |
| Transition between the two steps of splicing | **C*** Catalytic step 2 spliceosome | Prp16 promotes the release of the step 1 factors and binding of Prp22 and the step 2 factors Prp18 and Slu7 | **Branching triggers a rotation of the BP helix on its axis. The intron termini pair pulls on the 3'exon. U5 loop 1 facilitates the translocation of the 3'exon due to the relaxation caused by the snipping of the intron bond from the 5'exon. The $G_{-1}$ and 3' exon position for the step 2 catalysis.** |
| Exon ligation | **P** Post-splicing spliceosome |  | Final products: Ligated exons and the intron lariat |
| Release of the ligated exons | **ILS** Intron-lariat spliceosome | Prp22 promotes the release of step 2 protein factors | Disassociation of the ligated exons |
| Spliceosome disassembly | **U6, U5, U2 snRNPs** | Prp43 | Release of the intron lariat |

[1] Wahl et al. 2009 and 2015, Scheres and Nagai 2017, Bertram et al. 2017[b]; [2] Sickmier et al. 2006, Chen et al. 2010, Kent et al. 2003; [3] based on Corrionero et al. 2011 and Brock et al. 2008; **in blue**: see Discussion.

**Table S2.** Diverse views of the U5 snRNA Loop1 interactions with exons were obtained by crosslinking experiments and Cryo electron microscopy by different research groups.

**S2.A.** Budding yeast *(Saccharomyces cerevisiae)* spliceosome

| Crosslinking with 4-thioU<br>Newman et al.1995;<br>Teigelkamp et al. 1995 | Luhrmann CryoEM<br>Rauhut et al. 2016 | Nagai CryoEM<br>Galej et al. 2016 | Shi CryoEM<br>Wan et al. 2016[a]; Yan et a<br>2016; Bai et al. 2017 |
|---|---|---|---|
| 5' exon<br>***CYH2* 5'ex 4-thioU**<br>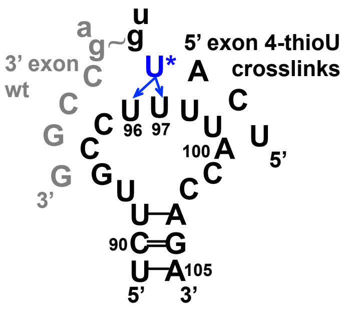<br><br>Detected before and after the 1st step (**Bact** and **C** complexes?).<br>**4-thioU replaced a G** (see wt 5' exon below).<br>The non-canonical intron termini pair is cited in this paper after Parker and Siliciano 1993, Chanfreau et al. 1994, Scadden and Smith 1995. | 5' exon<br>**Actin**<br>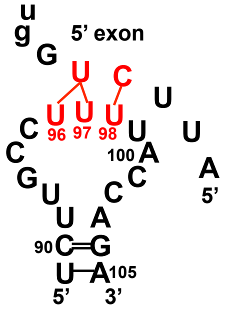<br><br>**Bact** complex | 5' exon<br>***UBC4***<br>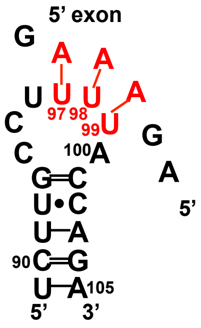<br><br>**C** complex<br>5' exon binding is the same in **C*** **(**Fica et al. 2017) and presumed the same in **Bact** (Scheres and Nagai, 2017) | 5' exon<br>***TUB3* pre-mRNA modelled in**<br>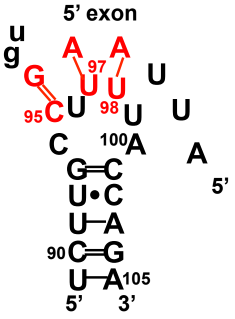<br><br>**B** complex<br><br>**Pre-mRNA modelled in**<br>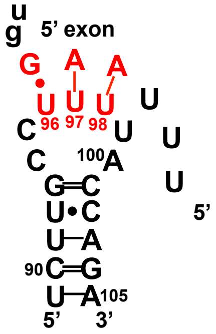<br><br>**Bact** complex<br>5' exon binding is the sam in **C** **(**Wan et al. 2016[b]) an **C***complex (Yan et al. 201 |
| 3' exon<br>***CYH2* 3'ex 4-thioU**<br>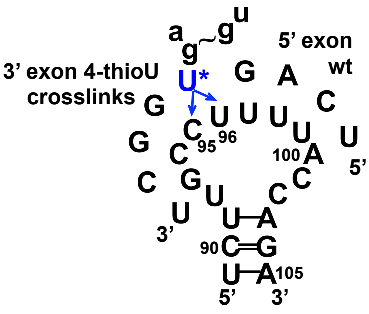<br><br>Detected in the lariat intermediate (**C** complex). 3' exon sequence is not the wt (see above). | | | 3' exon<br>**Pre-mRNA modelled in**<br>**5' exon**<br>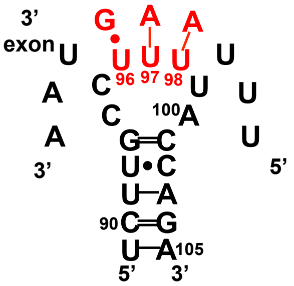<br><br>**P** complex<br>3' exon density is close to U5 Loop 1, but pairing configuration is unclear |

**S2.B.** Human spliceosome with adenovirus pre-mRNA substrates??

| Crosslinking with 4-thioU<br>Sontheimer and Steitz 1993 | Luhrmann CryoEM<br>Bertram et al. 2017a,b | Shi CryoEM<br>Zhang et al. 2017, 2018 |
|---|---|---|

5' exon

***Ad5-1* 5'ex 4-thioU**

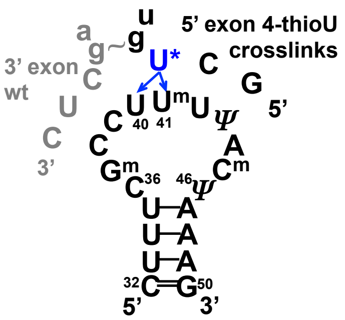


Detected before and after the 1st step (**Bact** and **C** complexes). **4-thioU replaced a G** (the subultimate G was also changed to C to avoid a GU, see wt 5' exon below).
The non-canonical intron termini pair is cited in this paper after Parker and Siliciano 1993.

5' exon

**MINX**

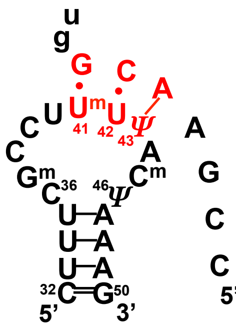

**B** complex

**MINX**

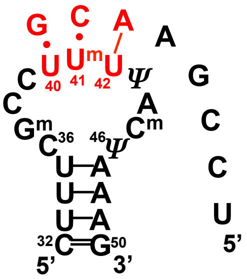

**C*** complex

5' exon

**MINX**

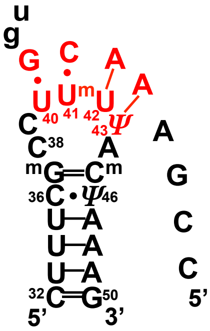

**Bact** complex

**MINX**

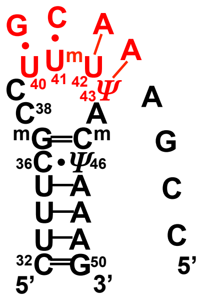

**C*** complex
5' exon binding is presumed the same in **C** (Zhan et al. 2018)

---

3' exon

***Ad3+1* 3'ex 4-thioU**

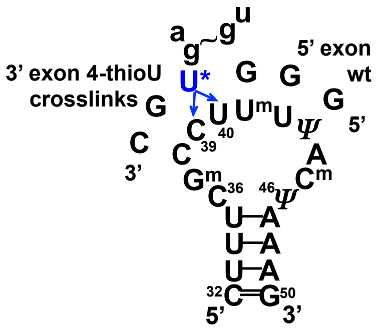


Detected in the lariat intermediate (**C** complex). 3' exon sequence is not the wt (see above).

? See **Table S1** for definitions of the stages of splicing as distinct RNP complexes
?? The pre-mRNA substrates are the derivatives of the Adenovirus-2 major late transcription unit. *Ad5-1* and *Ad3+1* are 4-thioU 5' and 3'ss modifications of the *Adeno* 'standard splicing substrate' (409nt): late exons 1 and 2 separated by the intron 1 with most of it deleted ∆198-982 (Sontheimer and Steitz 1993; Wyatt et al.1992; Solnick 1985).

**Table S3**. Spliceosomal RNA-RNA interactions confirmed by genetic and biochemical studies are not always captured by Cryo electron microscopy. Different stages of splicing are defined as distinct RNP complexes: **B** to **ILS** (see **Table S1**). RNA bp are defined by Westhof geometry (Leontis et al. 2002)

**S3.1** Budding yeast *(Saccharomyces cerevisiae[?])* spliceosome

| **Laboratory**<br>Method used to stall at a particular stage of splicing *in vitro* or to sort the EM images of the isolated total spliceosomes | | **R. Lührmann**<br>prp2-1 heat inactivation | **K. Nagai**<br>3'ss mutations | **Y. Shi**<br>Cef1 affinity[??] - 2D/3D image classification |
|---|---|---|---|---|
| **Spliceosomal complexes**, references, additional methods used before isolation to enrich for spliceosomes at a particular stage of splicing[???] | **B** | | | Wan et al. 2016[a]<br>Prp6 tagged |
| | **Bact** | Rauhut et al. 2016 | | Yan et al. 2016 |
| | **C** | | Galej et al. 2016 | Wan et al. 2016[b] |
| | **C*** | | Fica et al. 2017 | Yan et al. 2017 |
| | **P** | | | Bai et al. 2017<br>Prp22-K512A |
| | **ILS** | | | Wan et al. 2017<br>Yju2 tagged |
| Pre-mRNA substrate | | M3-actin:<br>End of exon 1 - intron 1 - start of exon 2<br>3x MS2 tags | *UBC4* mutants:<br>wt uagAG<br>**C** uacAC<br>**C*** ua($^{d}$g)AG | Unknown fragments, isolated with them |
| **Intron termini pair**<br>**G–○–G** (+1, -1)<br>Confirmed by mutation analysis, interchangeable for a double mutant<br>**A–○–C** (+1, -1)<br>nearly isosteric in this configuration (Parker and Siliciano 1993, Chanfreau et al. 1994) | | **Absent** | **Cannot form due to G$_{-1}$ substitutions** | Captured in **P**<br>**G●–■G** (+1, -1)<br>The strands are **parallel**, but the configuration is impossible for A-C and A-A<br>**Absent in B--Bact-C-C*-ILS** |
| **5' intron end - U6 snRNA**<br>**U—A** (+2, 51)<br>Detected by cross-linking experiments in **Bact** (stalled by prp2-1 heat inactivation) and **C** complexes (Kim and Abelson 1996, Fabrizio and Abelson 1990)<br>**A□–▷G** (+3, 50)<br>Confirmed by mutation analysis (*RPL30* gene), interchangeable with<br>**A□–▷A** (+4, 49)<br>isosteric in this configuration (Konarska et al. 2006) | | **U$_{+2}$** and **A$_{+3}$** are unpaired in **Bact** | Absent in **C**<br>Captured in **C***<br>**U○–□A** (+2, 51)<br>Captured in **C**<br>**A□–▷G** (+3, 50)<br>Absent in **C*** | **U$_{+2}$** and **A$_{+3}$** are unpaired in **B-Bact-C-C*-P--ILS** |

[?] The structure for *Schizosaccharomyces pombe* **ILS** complex is reported by Yan et al., 2015

[??] Cef1 protein is a component of the NTC complex, which joins the spliceosome when the catalytic core is formed, stays part of it during both steps of catalysis and remains with the intron lariat after the release of the ligated exons. Therefore, isolation by Cef1 affinity renders a mixture of **Bact, B*, C, C*, P** and **ILS** complexes.

[???] To facilitate isolation of complexes with U4/U6.U5 tri-snRNP Wan et al. 2016[a] used yeast expressing tagged Prp6, a protein component of the U5 snRNP. To enrich for spliceosomes stalled before exon ligation Bai and co-workers added Prp22-K512A ATPase defective mutant to the yeast cultures. To eliminate the 1st step spliceosomes form the isolated mixture Wan et al. 2016[b] used yeast expressing tagged 1st step factor Yju2.

**S3.2** Human spliceosome built on the miniature adenovirus pre-mRNA substrate (MINX[?])

| Laboratory | | R. Lührmann | Y. Shi |
|---|---|---|---|
| **Spliceosomal complexes**, references, method used to stall at a particular stage of splicing *in vitro* | **B** | Bertram et al. 2017[b]<br>Low $Mg^{2+}$, Brr2 stalled | |
| | **Bact** | | Zhang et al. 2018<br>3’ part deletion |
| | **C** | | Zhan et al. 2018<br>BN82685 inhibition[???] |
| | **C*** | Bertram et al. 2017[a]<br>Low pH 6.4[??] | Zhang et al. 2017<br>3’ss mutation |
| | **P** | | Zhang et al., 2019 |
| | **ILS** | | Zhang et al., 2019 |
| Pre-mRNA substrate | | Uniformly [$^{32}$P]-labeled $m^7$G(5’)ppp(5’)G-capped **MINX** | $m^7$G(5’)ppp(5’)G-capped-**MINX-15**: 3’ part deletion starting 19 nts downstream the branch point (**Bact** complex); **MINX** (**C** complex); **MINX-GG**: ag to gg mutation at the intron end (**C*** complex) |
| **Intron termini pair**<br>**G-o-G** ($G_{+1}$, $G_{-1}$)<br>Confirmed by mutation analysis, interchangeable for double mutants<br>**A-o-C** ($A_{+1}$, $C_{-1}$) **A-o-A** ($A_{+1}$, $A_{-1}$)<br>nearly isosteric in this configuration (Scadden and Smith 1995) | | Absent in **B** and **C*** | Cannot form in **Bact** due to the deletion of the 3’ part of the intron<br><br>Absent in **C** and **C*** |
| **5’ intron end - U6 snRNA**<br>**U—A** ($U_{+2}$, $A_{45}$)<br>Detected by cross-linking experiments in **C** complex (lariat intermediate) and **P** complex (lariat intron product). (Sontheimer and Steitz 1993) | | $\mathbf{U_{+2}}$ is unpaired in **B** and **C*** | $\mathbf{U_{+2}}$ and $\mathbf{A_{+3}}$ are unpaired in **Bact, C and C** |
| **A□–▷G** ($A_{+3}$, $G_{44}$) **A□–▷A$^{m6}$** ($A_{+4}$, $A_{43}$)<br>These pairs can be inferred from the mutation analysis in yeast (*RPL30* gene by Konarska et al. 2006) as both $\mathbf{A_{+3}}$ and $\mathbf{A_{+4}}$ are conserved in humans (78 and 68% of the human dystrophin gene introns respectively). The presence of at least one of these adenines supports the formation of the key non-WC pair. | | $\mathbf{A_{+3}}$ and $\mathbf{A_{+4}}$ are aligned with U6 $\mathbf{G_{44}}$ and $\mathbf{A^{m6}_{43}}$, respectively at the start of the $\mathbf{A_{39}\Psi_{40}A_{41}C_{42}}$ helix in **B** and **C*,** but base-pairing configuration is not specified. | $\mathbf{A_{+4}}$ is near $\mathbf{A^{m6}_{43}}$ in **Bact, C** and **C***, but base-pairing configuration is not specified. |

[?] MINX - **Mini**ature substrate, a derivative of the Adenovirus-2 major late pre-mRNA. pMINX (220bp) contains 3’ and 5’ parts of exon 2 separated by a small composite intron consisting of the start of intron 2 (5’ss) followed by a short stretch of the plasmid vector and the end of intron 1 with the branch point and 3’ss (Zillmann et al. 1988; Padgett et al. 1984)

[??] *In vitro* splicing proceeds normally at pH7.9 in the presence of HeLa nuclear extracts

#### Section S2: Figures

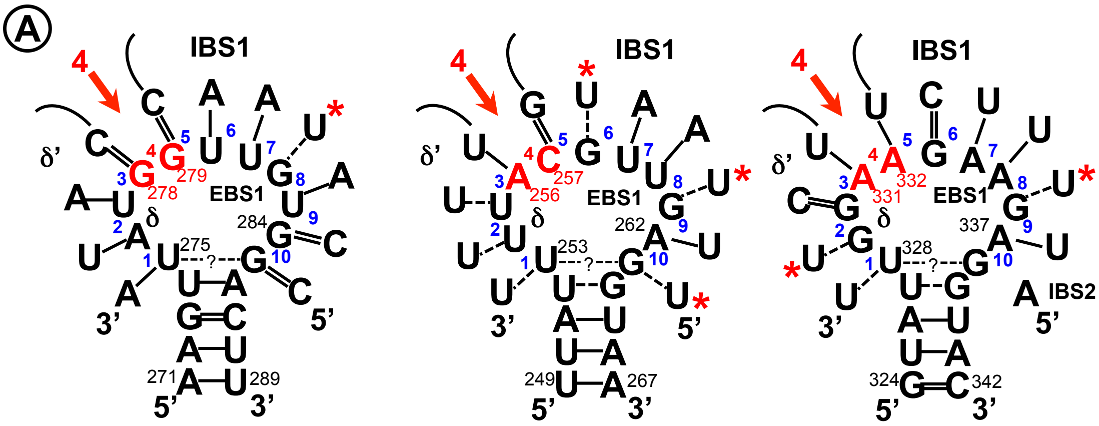


**Ll.LtrB** (L.l.I1)
intron of the *ltrB* gene encoding a relaxase found in the conjugative elements of *Lactococcus lactis*

**aI1** (cox1.I1 ) **aI2** (cox1.I2)
introns of the mitochondrial gene *cox1* encoding cytochrome c oxidase subunit I in *Saccharomyces cerevisiae*

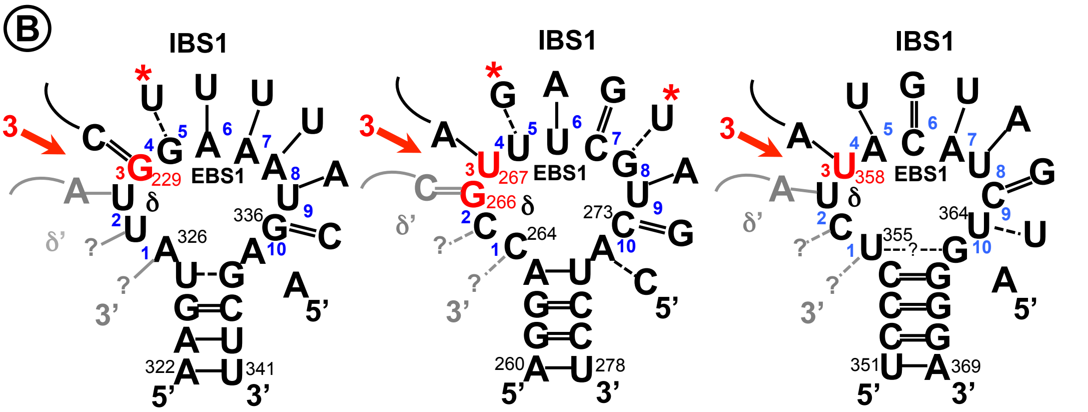


**ai5γ**
intron of the mitochondrial gene *cox1* encoding cytochrome *c* oxidase subunit I in *Saccharomyces cerevisiae*

**RmInt1** (Sr.me.I1)
intron found within the insertion sequence IS *Rm2011-2* in bacterium *Sinorhizobium meliloti,* a nitrogen-fixing symbiont of alfalfa *(Medicago sativa)*

**Avi.groEL** (A.v.I1)
intron of an essential housekeeping gene required for protein folding, chaperonin *groEL*, found in *Azotobacter vinelandii*

**Figure S1**| Diverse group II introns recognise their exons mainly by Watson-Crick base pairing with the Id3 loop in a common asymmetrical fashion. The arows indicate positions for the exon junction **3** or **4** referring to all 'possible' positions numbered in blue.

**A**. Id3 loops of these **Group IIA** introns bind both exons: 6-7nt at the end of the 5' exon and 1-4nt at the start of the 3' exon (Plante and Cousineau 2006, Zhong and Lambowitz 2003, Ichiyanagi et al. 2002, Eskes et al. 1997, 2000).

IBS1, Intron Binding Sequence 1 is the end of the 5'exon, δ' sequence is start of the 3'exon. EBS1, Exon Binding Sequence 1 and δ sequence of the Id3 loops bind the 5' and 3' exons respectively.

**B**. Id3 loop of **Group IIB** introns typically interacts with 7-8nt of the 5' exon and secures the 3' exon via a coordination loop and a further EBS3 – IBS3 interaction (Somarowthu et al. 2014, Ferat et al. 2003, Chillón et al. 2014). EBS3 of the coordination loop binds IBS3 of the 3' exon. δ' sequence of the coordination loop is shown in grey.

* G-U and U-G pairs are indicated with red asterisks

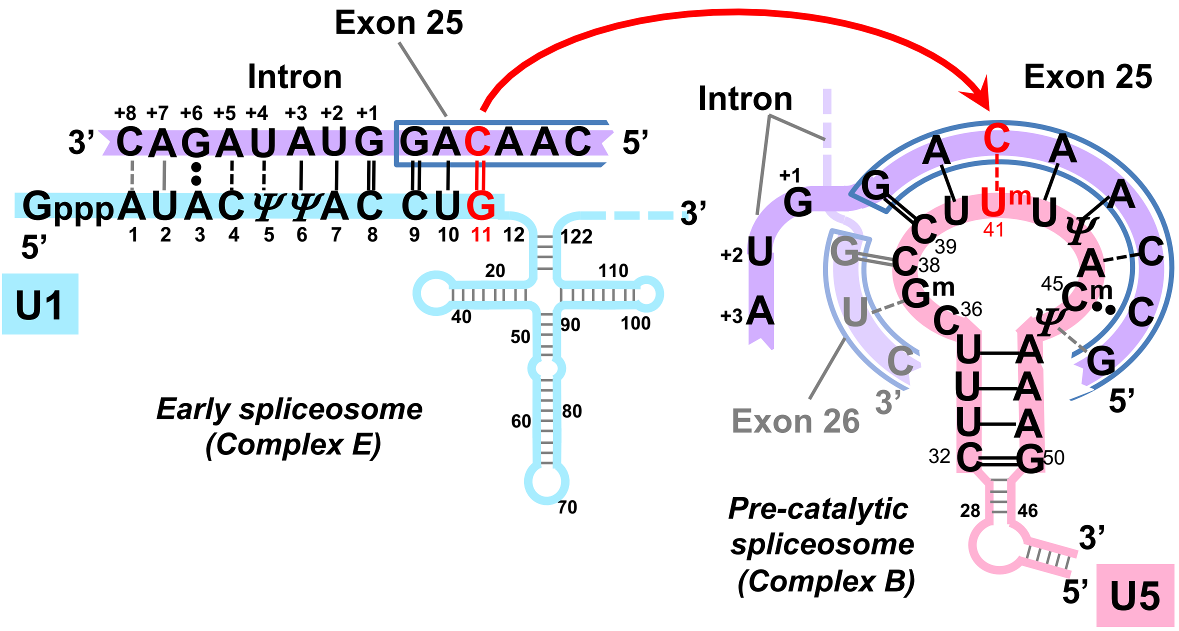


**Figure S2| 'Legal' C--U pairs in the U5 interactions prompted by the initial U1 selection** (Human dystrophin exon/intron 25 as an example).
Base modifications as in **Figure 1** caption.

***Sugar-phosphate backbone orientation (+/o) relative to the plane of a base***

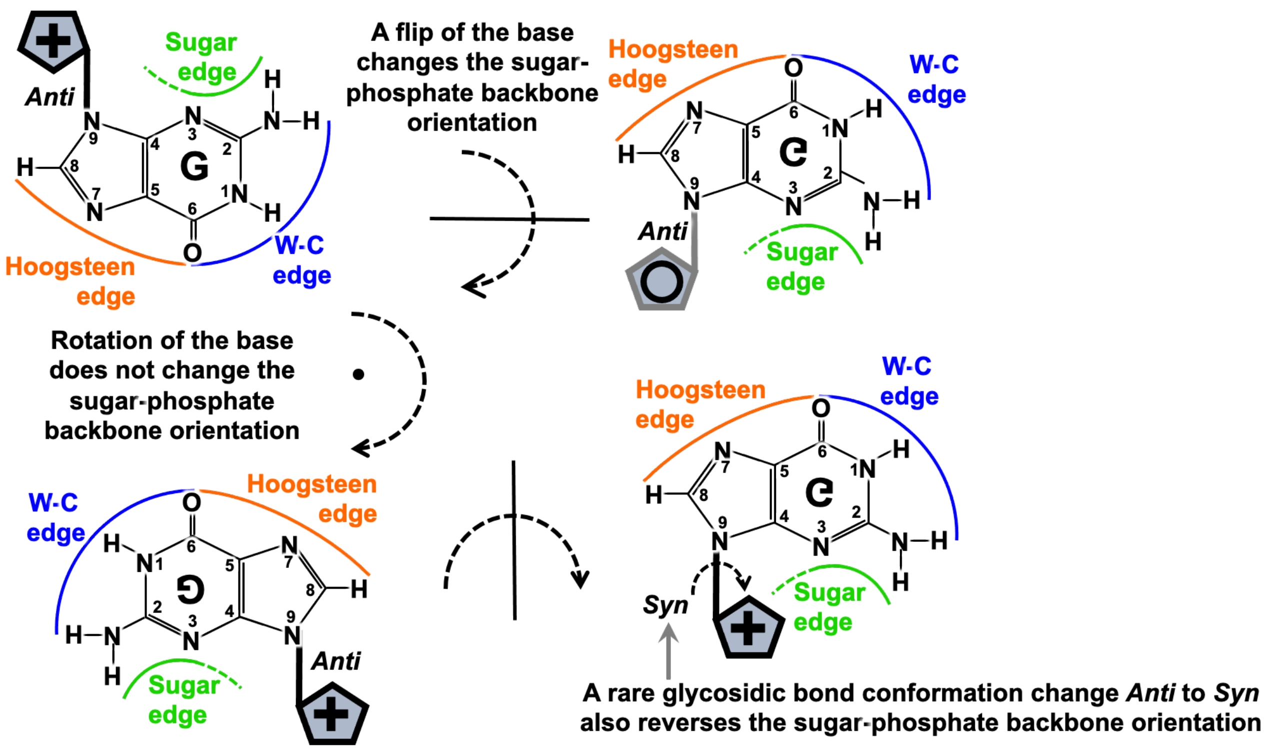


**Figure S3| Quick guide to sugar-phosphate backbone orientation relative to the plane of a base (explanatory schematics to accompany Figure 13).** Ribose is located on the perpendicular plane and is shown as a schematic blue pentagon with **+** and **o** indicating the opposite directions of the sugar-phosphate backbone.

***Canonical Watson-Crick, antiparallel strands, 1st Westhof geometric family***

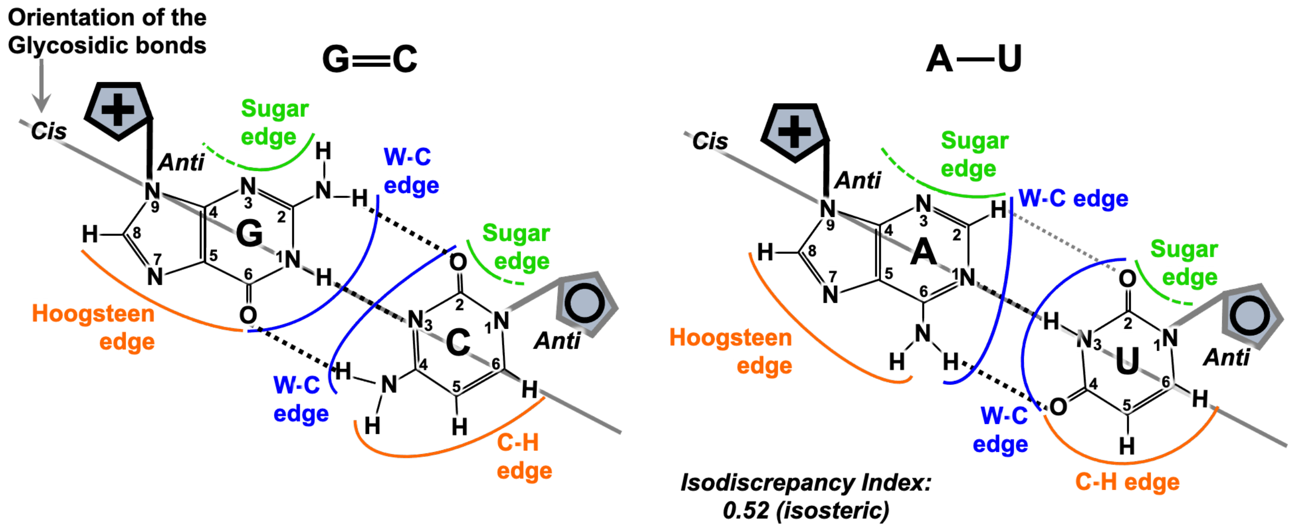


**Figure S4| Canonical Watson-Crick pairs with antiparallel strands.** Isodiscrepancy index is a numerical measure of geometric similarity (isostericity) of base pairs (on-line RNA base pair catalogue http://ndbserver.rutgers.edu/ndbmodule/services/BPCatalog/bpCatalog.html).

#### Section S3: Comments to Figures

##### Comment S1 (to Figure 12)
##### The odd feature of the of U12 interaction with the U6atac catalytic triad in the human minor spliceosome: a bulge or a Watson-Crick-like mimic pair?

According to the review of Turunen et al. 2013, which faithfully reproduces the original Tarn and Steitz 1996[b], U12 $C_4$ is bulging out of the helix U12 forms with the minor spliceosome catalytic triad U6atac $A_{26}G_{27}C_{28}$ (compare Helix Ib in **Figure 10C** and **10A**). This is a surprising feature, as we assume the same spatial organisation of the key catalytic triple helix in the paralogous spliceosomes. In the minor spliceosome of *Arabidopsis* there is no bulging inside Helix Ib (Shukla and Padget 1999), although the binding register is currently disputed (Ciavarella et al., 2020).
There is a possibility, hitherto unexplored, that bulging inside Helix Ib can be avoided by formation of a Watson-Crick-like pair of $A_{26}$--$C_5$ followed by $G_{27}$=$C_4$ (residue numbers as in **Figure 10C**).
To test this, we propose alternative substitutions in the sequence of U12 snRNA:

1) $U_6$ for A or G to abolish U6atac $A_{26}$-$U_6$: this mutant can only be functional if the binding register includes $A_{26}$--$C_5$ and $G_{27}$=$C_4$
2) $C_5$ for U, to introduce $A_{26}$-$U_5$, which should be functionally equivalent to $A_{26}$--$C_5$
3) $C_4$ for A or G, which maybe partly functional if the proposed new binding register is wrong and $C_4$ is indeed a bulge

##### Comment S2 (to Figure 14)
##### The stalled transition between 2 steps of splicing in the absence of the Prp16 helicase

The conclusion that the transition between the two steps of splicing is triggered and transacted by the RNA component is consistent with the conservation of the lariat intermediate between the spliceosome and Group II introns, but it seems to disagree with the fact that spliceosome is stalled after the first reaction in the absence of Prp16. But let us consider the dual role of helicases: their role in remodelling of the spliceosome and alternative splicing. Prp16 uses its helicase activity to dissociate suboptimal branch point sites from U2 snRNA, as it is conclusively proven in the experiment with deoxy A substitutions at the branch point (Semlow et al. 2016), but there is no evidence that it translocates along the RNA strand and unwinds RNA duplexes in order to mediate the transition between the 1$^{st}$ and 2$^{nd}$ steps. In fact, there is a proof that it does not translocate across the branch point interaction (Semlow et al. 2016). What is known is that the function of Prp16 is blocked if the nucleotides -4 to -11 at the end of the intron are substituted by their deoxy counterparts. This is the region analogous to the mammalian PPT (**Figure 10A**), the least conserved of the 3' intron end sequences and likely to be involved in RNA-protein interactions. The example of Prp28 highlights, that the helicase activity as such might be not employed in the spliceosome remodelling. Instead, Prp28 initiates the dissociation of the 5' splice site from the U1snRNA by displacing another protein, U1C from the RNA duplex to destabilise it (Chen et al. 2001). Could it be that Prp16 also has a protein target and dissociation of this target facilitates the conformational change of the BP adenosine after branching? This would be a logical explanation, as the preserved ancient mechanism of ribozyme activity would need to be adjusted to maintain the complexity of the spliceosomal proteome, which becomes largely influential for the regulation of alternative splicing in the metazoan genome.

#### Section S4: Results and Methods - Corelated base pair variation analyses

##### **List S1.** List of human genes included in the analysis by sKL divergence and Bootstrap

Human genes ranging from well known in medical genetics practice to genes with experimentally confirmed function and expression were chosen at random avoiding paralogues (for example: dystrophin is included, but utrophin is not). Aiming for approximately 2000 introns (splice junctions), 132 genes were included altogether.

(SJ=number of splice junctions, Ch=chromosome)

| # | *Gene name* | *SJ* | *Ch* |
|---|---|---|---|
| 1 | *DMD*, dystrophin | 78 | X |
| 2 | *F5,* coagulation factor V | 24 | 1 |
| 3 | *CFTR*, cystic fibrosis transmembrane conductance regulator | 26 | 7 |
| 4 | *F8*, coagulation factor VIII | 25 | X |
| 5 | *F9*, coagulation factor IX | 7 | X |
| 6 | *VWF*, von Willebrand factor | 51 | 12 |
| 7 | *SMN1*, survival of motor neuron 1 | 8 | 5 |
| 8 | *PAH*, phenylalanine hydroxylase (PKU, phenylketonuria) | 12 | 12 |
| 9 | *APOE*, apolipoprotein E | 3 | 19 |
| 10 | *BRCA1*, breast cancer 1, DNA repair associated | 23 | 17 |
| 11 | *HPRT*, hypoxanthine phosphoribosyltransferase 1 | 8 | X |
| 12 | *HBB*, haemoglobin subunit beta | 2 | 11 |
| 13 | *Fas/CD95*, Fas cell surface death receptor | 8 | 10 |
| 14 | *TP53*, tumor protein p53 | 11 | 17 |
| 15 | *TNF*, tumour necrosis factor | 3 | 6 |
| 16 | *UBC*, ubiquitin C | 1 | 12 |
| 17 | *EGFR*, epidermal growth factor receptor | 27 | 7 |
| 18 | *VEGFA*, vascular endothelial growth factor | 6 | 6 |
| 19 | *IL6*, interleukin 6 | 5 | 7 |
| 20 | *TGFB1*, transforming growth factor, beta | 6 | 19 |
| 21 | *ESR1*, oestrogen receptor 1 | 9 | 6 |
| 22 | HLA-DRB1, major histocompatibility complex, class II, DR beta 1 | 5 | 6 |
| 23 | *NFKB1*, nuclear factor of kappa | 23 | 4 |
| 24 | *IL10*, interleukin 10 | 4 | 1 |
| 25 | *AKT1*, AKT serine/threonine kinase 1 | 12 | 14 |
| 26 | *CD4*, cluster of differentiation 4 glycoprotein | 9 | 12 |
| 27 | *GRB2*, growth factor receptor-bound protein 2 | 5 | 17 |
| 28 | *ELP1/IKBKAP,* elongator complex protein 1 | 36 | 9 |
| 29 | *NOVA1*, alternative splicing regulator 1 Neuro-Oncological Ventral Antigen 1 | 4 | 14 |
| 30 | *TOP1*, topoisomerase I | 20 | 20 |
| 31 | *INS*, insulin | 1 | 11 |
| 32 | *FHL5/ACT*, four and a half LIM domains 5 | 6 | 6 |
| 33 | *TPM1*, tropomyosin | 8 | 15 |
| 34 | *DAG1*, dystroglycan 1 | 5 | 3 |
| 35 | *POLR2A*, RNA polymerase II subunit A isoform 201 | 9 | 17 |
| 36 | *PRPF8*, Prp8, pre-mRNA processing factor 8 | 42 | 17 |
| 37 | *DDX8*, Prp22, pre-mRNA processing factor 22 | 22 | 17 |
| 38 | *SNRNP200*, sn ribonucleoprotein U5 subunit 200, Brr2 helicase, | 44 | 2 |
| 39 | *MYC*, proto-oncogene, bHLH transcription factor | 2 | 8 |
| 40 | *TBP*, TATA-box binding protein | 7 | 6 |
| 41 | *LAMB1*, laminin subunit beta 1 | 33 | 7 |
| 42 | *NOTCH1*, notch receptor 1 | 33 | 9 |
| 43 | *SGCA*, sarcoglycan alpha | 9 | 17 |
| 44 | *TUBA1A,* tubulin | 3 | 12 |
| 45 | *MYO7A*, myosin VIIA | 48 | 11 |
| 46 | *NUP155*, nucleoporin 155 | 34 | 5 |
| 47 | *PTBP1*, polypyrimidine tract binding protein 1 | 14 | 19 |
| 48 | *U2AF2*, U2AF65, U2 small nuclear RNA auxiliary factor 2 | 11 | 19 |
| 49 | *GAPDH*, glyceraldehyde-3-phosphate dehydrogenase | 8 | 12 |
| 50 | *TK1*, thymidine kinase 1 | 6 | 17 |
| 51 | *ATM*, serine/threonine kinase (ataxia telangiectasia mutated) | 62 | 11 |
| 52 | *TERT*, telomerase reverse transcriptase | 15 | 5 |
| 53 | *ADAMTS13*, metallopeptidase with thrombospondin type 1 motif 13 | 28 | 9 |

| | | | |
|---|---|---|---|
| 54 | *NEU1*, neuraminidase 1 | 5 | 6 |
| 55 | *GLB1*, beta-galactosidase | 15 | 3 |
| 56 | *CTSA*, cathepsin A | 14 | 20 |
| 57 | *ACE*, angiotensin I converting enzyme | 24 | 17 |
| 58 | *ERN1*, endoplasmic reticulum to nucleus signaling 1 | 21 | 17 |
| 59 | *BMP2*, bone morphogenetic protein 2 | 2 | 20 |
| 60 | *CDK6*, cyclin dependent kinase 6 | 7 | 7 |
| 61 | *CCND1*, cyclin D1 | 4 | 11 |
| 62 | *EZH1,* enhancer of zeste 1 polycomb repressive complex 2 subunit | 20 | 17 |
| 63 | *HAT1*, histone acetyltransferase 1 | 10 | 2 |
| 64 | *ITGB3*, integrin subunit beta 3 | 14 | 17 |
| 65 | *IFNG*, interferon gamma | 3 | 12 |
| 66 | *ERBB2*, erb-b2 receptor tyrosine kinase 2 | 30 | 17 |
| 67 | *APP*, amyloid beta precursor protein | 17 | 21 |
| 68 | *EGF*, epidermal growth factor | 23 | 4 |
| 69 | *CTNNB1*, catenin beta 1 | 14 | 3 |
| 70 | *IGF1*, insulin like growth factor 1 | 5 | 12 |
| 71 | *MAPK1*, mitogen-activated protein kinase 1 | 8 | 22 |
| 72 | *IGHG1*, immunoglobulin heavy constant gamma 1 | 5 | 14 |
| 73 | *TFRC*, transferrin receptor | 18 | 3 |
| 74 | *CRP*, C-reactive protein | 1 | 1 |
| 75 | *MET*, proto-oncogene, receptor tyrosine kinase | 20 | 7 |
| 76 | *RGS6*, regulator of G protein signaling 6 | 17 | 14 |
| 77 | *BCL2*, apoptosis regulator | 1 | 18 |
| 78 | *XRCC6*, X-ray repair cross complementing 6, Ku70 | 12 | 22 |
| 79 | *RAD52,* RAD52 homolog, DNA repair protein | 11 | 12 |
| 80 | *MSTN*, myostatin (GDF8, growth and differentiation factor 8) | 2 | 2 |
| 81 | *PNMT,* phenylethanolamine N-methyltransferase | 2 | 17 |
| 82 | *CELF1*, CUGBP Elav-like family member 1 (splicing regulator) | 15 | 11 |
| 83 | *BLM,* RecQ like helicase (Bloom syndrome) | 21 | 15 |
| 84 | *PSEN1*, FAD, presenilin 1 ($\gamma$-secretase subunit, processing of APP) | 11 | 14 |
| 85 | *AMY1A*, amylase alpha 1A (salivary) | 10 | 1 |
| 86 | *SPSB1*, splA/ryanodine receptor domain, SOCS box containing-1 (SSB1, ssDNA binding-1) | 2 | 1 |
| 87 | *G6PD*, glucose-6-phosphate dehydrogenase | 12 | X |
| 88 | *USB1*, U6 snRNA biogenesis phosphodiesterase 1 | 6 | 16 |
| 89 | *ERCC6*, excision repair 6, chromatin remodeling factor | 20 | 10 |
| 90 | *ATP2B1*, ATPase plasma membrane Ca2+ transporting 1 | 19 | 12 |
| 91 | *ETV4*, ETS variant transcription factor 4 | 12 | 17 |
| 92 | *HNF1A*, HNF1 homeobox A | 9 | 12 |
| 93 | *NAT2*, N-acetyltransferase 2, AAC2 | 1 | 8 |
| 94 | *LPL*, lipoprotein lipase | 9 | 8 |
| 95 | *FADS1*, fatty acid desaturase 1 | 11 | 11 |
| 96 | *GDF5*, growth differentiation factor 5 | 1 | 20 |
| 97 | *ACADM*, acyl-CoA dehydrogenase medium chain | 11 | 1 |
| 98 | *GATA1,* GATA binding protein 1 | 5 | X |
| 99 | *GSTM1*, glutathione S-transferase mu 1 | 7 | 1 |
| 100 | *CDH1*, cadherin 1 | 15 | 16 |
| 101 | *FMR1*, fragile X mental | 16 | X |
| 102 | *POLD1*, DNA polymerase delta 1, catalytic subunit | 26 | 19 |
| 103 | *RPA*, replication protein A | 16 | 17 |
| 104 | *WRN*, WRN RecQ like helicase | 34 | 8 |
| 105 | *SP1*, Sp1 transcription factor | 5 | 12 |
| 106 | *TWIST1*, twist family bHLH transcription factor 1 | 1 | 7 |
| 107 | *TAF1*, TATA-box binding protein associated factor 1 | 37 | X |
| 108 | *POLB,* DNA polymerase beta | 13 | 8 |
| 109 | *TERF2*, telomeric repeat binding factor 2 | 9 | 16 |
| 110 | *RFC1*, replication factor C, subunit 1 | 24 | 4 |
| 111 | *MAPT*, microtubule associated protein tau | 14 | 17 |
| 112 | *ACTA2, actin alpha 2, smooth muscle* | 8 | 10 |
| 113 | *PCGF2*, polycomb group ring finger 2 | 10 | 17 |

| | | | |
|---|---|---|---|
| 114 | *SSBP3*, single stranded DNA binding protein 3 | 17 | 1 |
| 115 | *CHEK1,* checkpoint kinase 1 | 12 | 11 |
| 116 | *BRIP1,* BRCA1 interacting protein C-terminal helicase 1 | 19 | 17 |
| 117 | *NFATC2*, nuclear factor of activated T cells 2 | 10 | 20 |
| 118 | *PRKDC*, protein kinase, DNA-activated, catalytic subunit, DNA-PKC | 85 | 8 |
| 119 | *OPTN, optineurin* | 14 | 10 |
| 120 | *ZBP1,* Z-DNA binding protein 1 | 7 | 20 |
| 121 | *LDHA,* lactate dehydrogenase A | 7 | 11 |
| 122 | *RBPJ,* recombination signal binding protein for immunoglobulin κ J region | 11 | 4 |
| 123 | *FKTN,* fukutin | 9 | 9 |
| 124 | *SFPQ*, splicing factor proline and glutamine rich | 9 | 1 |
| 125 | *MAD1L1,* mitotic arrest deficient 1 like 1 | 16 | 7 |
| 126 | *RAD51,* recombinase, RecA | 9 | 15 |
| 127 | *POLA1,* DNA polymerase alpha 1, catalytic subunit | 36 | X |
| 128 | *PCNA,* proliferating cell nuclear antigen | 5 | 20 |
| 129 | *PRIM2,* DNA primase subunit 2 | 4 | 6 |
| 130 | *ITPR1,* inositol 1,4,5-trisphosphate receptor type 1 | 58 | 3 |
| 131 | *EFNA5,* ephrin A5 | 4 | 5 |
| 132 | *OPA1,* mitochondrial dynamin like GTPase | 29 | 3 |
| | | Total SJ: **2007** | |

```
>>>
= RESTART: /Users/olga/Documents/Bioinformatics MSc Birkbeck/Codes for U5 paper/splice_sites.py
```

Human gene sequences were downloaded from *ensembl* with two separate files for exons and introns. The splice isoforms were selected to have:

- Experimental evidence (RefSeq, **TSL1**=transcript support level 1)
- Maximum splice junctions for a particular gene

**The *ensembl* isoform number is featured in the sequence file names.**

Extracting slice junctions from 132 human genes

```
001_DMD_203_exons.txt
Splice junctions in this gene:  78
002_F5_202_exons.txt
Splice junctions in this gene:  24
003_CFTR_201_exons.txt
Splice junctions in this gene:  26
004_F8_202_exons.txt
Splice junctions in this gene:  25
005_F9_201_exons.txt
Splice junctions in this gene:  7
006_VWF_201_exons.txt
Splice junctions in this gene:  51
007_SMN1_202_exons.txt
Splice junctions in this gene:  8
008_PAH_215_exons.txt
Splice junctions in this gene:  12
009_APOE_201_exons.txt
Splice junctions in this gene:  3
010_BRCA1_210_exons.txt
Splice junctions in this gene:  23
011_HPRT1_201_exons.txt
Splice junctions in this gene:  8
012_HBB_206_exons.txt
Splice junctions in this gene:  2
013_FAS_221_exons.txt
Splice junctions in this gene:  8
014_TP53_223_exons.txt
Splice junctions in this gene:  11
015_TNF_208_exons.txt
Splice junctions in this gene:  3
016_UBC_201_exons.txt
Splice junctions in this gene:  1
017_EGFR_201_exons.txt
Splice junctions in this gene:  27
018_VEGFA_205_exons.txt
Splice junctions in this gene:  6
019_IL6_204_exons.txt
Splice junctions in this gene:  5
020_TGFB1_201_exons.txt
Splice junctions in this gene:  6
021_ESR1_207_exons.txt
Splice junctions in this gene:  9
022_HLA_DRB1_201_exons.txt
Splice junctions in this gene:  5
023_NFKB1_201_exons.txt
Splice junctions in this gene:  23
024_IL10_202_exons.txt
Splice junctions in this gene:  4
025_AKT1_208_exons.txt
Splice junctions in this gene:  12
026_CD4_201_exons.txt
Splice junctions in this gene:  9
027_GRB2_203_exons.txt
Splice junctions in this gene:  5
028_ELP1_201_exons.txt
Splice junctions in this gene:  36
029_NOVA1_206_exons.txt
Splice junctions in this gene:  4
030_TOP1_201_exons.txt
Splice junctions in this gene:  20
031_INS_203_exons.txt
Splice junctions in this gene:  1
032_FHL5_201_exons.txt
Splice junctions in this gene:  6
033_TPM1_206_exons.txt
Splice junctions in this gene:  8
034_DAG1_222_exons.txt
Splice junctions in this gene:  5
035_POLR2A_201_exons.txt
Splice junctions in this gene:  9
036_PRPF8_201_exons.txt
Splice junctions in this gene:  42
037_DHX8_201_exons.txt
Splice junctions in this gene:  22
038_SNRNP200_201_exons.txt
Splice junctions in this gene:  44
039_MYC_207_exons.txt
Splice junctions in this gene:  2
040_TBP_202_exons.txt
Splice junctions in this gene:  7
041_LAMB1_201_exons.txt
Splice junctions in this gene:  33
042_NOTCH1_205_exons.txt
Splice junctions in this gene:  33
043_SGCA_201_exons.txt
Splice junctions in this gene:  9
044_TUBA1A_202_exons.txt
Splice junctions in this gene:  3
045_MYO7A_202_exons.txt
Splice junctions in this gene:  48
046_NUP155_201_exons.txt
Splice junctions in this gene:  34
047_PTBP1_203_exons.txt
Splice junctions in this gene:  14
048_U2AF2_202_exons.txt
Splice junctions in this gene:  11
049_GAPDH_201_exons.txt
Splice junctions in this gene:  8
050_TK1_201_exons.txt
Splice junctions in this gene:  6
051_ATM_201_exons.txt
Splice junctions in this gene:  62
052_TERT_201_exons.txt
Splice junctions in this gene:  15
053_ADAMTS13_206_exons.txt
Splice junctions in this gene:  28
054_NEU1_206_exons.txt
Splice junctions in this gene:  5
```

055_GLB1_201_exons.txt
Splice junctions in this gene: 15
056_CTSA_204_exons.txt
Splice junctions in this gene: 14
057_ACE_202_exons.txt
Splice junctions in this gene: 24
058_ERN1_201_exons.txt
Splice junctions in this gene: 21
059_BMP2_201_exons.txt
Splice junctions in this gene: 2
060_CDK6_201_exons.txt
Splice junctions in this gene: 7
061_CCND1_201_exons.txt
Splice junctions in this gene: 4
062_EZH1_202_exons.txt
Splice junctions in this gene: 20
063_HAT1_201_exons.txt
Splice junctions in this gene: 10
064_ITGB3_201_exons.txt
Splice junctions in this gene: 14
065_IFNG_201_exons.txt
Splice junctions in this gene: 3
066_ERBB2_219_exons.txt
Splice junctions in this gene: 30
067_APP_201_exons.txt
Splice junctions in this gene: 17
068_EGF_201_exons.txt
Splice junctions in this gene: 23
069_CTNNB1_201_exons.txt
Splice junctions in this gene: 14
070_IGF1_203_exons.txt
Splice junctions in this gene: 5
071_MAPK1_201_exons.txt
Splice junctions in this gene: 8
072_IGHG1_202_exons.txt
Splice junctions in this gene: 5
073_TFRC_201_exons.txt
Splice junctions in this gene: 18
074_CRP_201_exons.txt
Splice junctions in this gene: 1
075_MET_201_exons.txt
Splice junctions in this gene: 20
076_RGS6_209_exons.txt
Splice junctions in this gene: 17
077_BCL2_202_exons.txt
Splice junctions in this gene: 1
078_XRCC6_202_exons.txt
Splice junctions in this gene: 12
079_RAD52_202_exons.txt
Splice junctions in this gene: 11
080_MSTN_201_exons.txt
Splice junctions in this gene: 2
081_PNMT_201_exons.txt
Splice junctions in this gene: 2
082_CELF1_214_exons.txt
Splice junctions in this gene: 15
083_BLM_201_exons.txt
Splice junctions in this gene: 21
084_PSEN1_201_exons.txt
Splice junctions in this gene: 11
085_AMY1A_201_exons.txt
Splice junctions in this gene: 10
086_SPSB1_201_exons.txt
Splice junctions in this gene: 2
087_G6PD_202_exons.txt
Splice junctions in this gene: 12
088_USB1_201_exons.txt
Splice junctions in this gene: 6
089_ERCC6_201_exons.txt
Splice junctions in this gene: 20
090_ATP2B1_201_exons.txt
Splice junctions in this gene: 19
091_ETV4_201_exons.txt
Splice junctions in this gene: 12
092_HNF1A_201_exons.txt
Splice junctions in this gene: 9
093_NAT2_201_exons.txt
Splice junctions in this gene: 1
094_LPL_207_exons.txt
Splice junctions in this gene: 9
095_FADS1_201_exons.txt
Splice junctions in this gene: 11
096_GDF5_201_exons.txt
Splice junctions in this gene: 1
097_ACADM_202_exons.txt
Splice junctions in this gene: 11
098_GATA1_202_exons.txt
Splice junctions in this gene: 5
099_GSTM1_201_exons.txt
Splice junctions in this gene: 7
100_CDH1_201_exons.txt
Splice junctions in this gene: 15
101_FMR1_205_exons.txt
Splice junctions in this gene: 16
102_POLD1_205_exons.txt
Splice junctions in this gene: 26
103_RPA1_201_exons.txt
Splice junctions in this gene: 16
104_WRN_201_exons.txt
Splice junctions in this gene: 34
105_SP1_201_exons.txt
Splice junctions in this gene: 5
106_TWIST1_201_exons.txt
Splice junctions in this gene: 1
107_TAF1_203_exons.txt
Splice junctions in this gene: 37
108_POLB_201_exons.txt
Splice junctions in this gene: 13
109_TERF2_201_exons.txt
Splice junctions in this gene: 9
110_RFC1_201_exons.txt
Splice junctions in this gene: 24
111_MAPT_204_exons.txt
Splice junctions in this gene: 14
112_ACTA2_201_exons.txt
Splice junctions in this gene: 8
113_PCGF2_207_exons.txt
Splice junctions in this gene: 10
114_SSBP3_204_exons.txt
Splice junctions in this gene: 17
115_CHEK1_213_exons.txt
Splice junctions in this gene: 12
116_BRIP1_201_exons.txt
Splice junctions in this gene: 19
117_NFATC2_201_exons.txt
Splice junctions in this gene: 10
118_PRKDC_201_exons.txt
Splice junctions in this gene: 85
119_OPTN_202_exons.txt
Splice junctions in this gene: 14
120_ZBP1_201_exons.txt
Splice junctions in this gene: 7
121_LDHA_205_exons.txt
Splice junctions in this gene: 7
122_RBPJ_205_exons.txt
Splice junctions in this gene: 11
123_FKTN_201_exons.txt
Splice junctions in this gene: 9
124_SFPQ_201_exons.txt
Splice junctions in this gene: 9
125_MAD1L1_201_exons.txt
Splice junctions in this gene: 16
126_RAD51_201_exons.txt
Splice junctions in this gene: 9
127_POLA1_201_exons.txt
Splice junctions in this gene: 36
128_PCNA_201_exons.txt
Splice junctions in this gene: 5
129_PRIM2_201_exons.txt
Splice junctions in this gene: 4
130_ITPR1_203_exons.txt
Splice junctions in this gene: 58
131_EFNA5_201_exons.txt
Splice junctions in this gene: 4
132_OPA1_205_exons.txt
Splice junctions in this gene: 29

**Splice junctions total: 2007**

#### Extracting introns (starts and ends) from 132 human genes

001_DMD_203_introns.txt
Introns in this gene: 78
002_F5_202_introns.txt
Introns in this gene: 24
003_CFTR_201_introns.txt
Introns in this gene: 26
004_F8_202_introns.txt
Introns in this gene: 25
005_F9_201_introns.txt
Introns in this gene: 7
006_VWF_201_introns.txt
Introns in this gene: 51
007_SMN1_202_introns.txt
Introns in this gene: 8
008_PAH_215_introns.txt
Introns in this gene: 12
009_APOE_201_introns.txt
Introns in this gene: 3
010_BRCA1_210_introns.txt
Introns in this gene: 23
011_HPRT1_201_introns.txt
Introns in this gene: 8
012_HBB_206_introns.txt
Introns in this gene: 2
013_FAS_221_introns.txt
Introns in this gene: 8
014_TP53_223_introns.txt
Introns in this gene: 11
015_TNF_208_introns.txt
Introns in this gene: 3
016_UBC_201_introns.txt
Introns in this gene: 1
017_EGFR_201_introns.txt
Introns in this gene: 27
018_VEGFA_205_introns.txt
Introns in this gene: 6
019_IL6_204_introns.txt
Introns in this gene: 5
020_TGFB1_201_introns.txt
Introns in this gene: 6
021_ESR1_207_introns.txt
Introns in this gene: 9
022_HLA_DRB1_201_introns.txt
Introns in this gene: 5
023_NFKB1_201_introns.txt
Introns in this gene: 23
024_IL10_202_introns.txt
Introns in this gene: 4
025_AKT1_208_introns.txt
Introns in this gene: 12
026_CD4_introns.txt
Introns in this gene: 9
027_GRB2_203_introns.txt
Introns in this gene: 5
028_ELP1_201_introns.txt
Introns in this gene: 36
029_NOVA1_206_introns.txt

Introns in this gene: 4
030_TOP1_201_introns.txt
Introns in this gene: 20
031_INS_203_introns.txt
Introns in this gene: 1
032_FHL5_201_introns.txt
Introns in this gene: 6
033_TPM1_206_introns.txt
Introns in this gene: 8
034_DAG1_222_introns.txt
Introns in this gene: 5
035_POLR2A_201_introns.txt
Introns in this gene: 9
036_PRPF8_201_introns.txt
Introns in this gene: 42
037_DHX8_201_introns.txt
Introns in this gene: 22
038_SNRNP200_201_introns.txt
Introns in this gene: 44
039_MYC_207_introns.txt
Introns in this gene: 2
040_TBP_202_introns.txt
Introns in this gene: 7
041_LAMB1_201_introns.txt
Introns in this gene: 33
042_NOTCH1_205_introns.txt
Introns in this gene: 33
043_SGCA_201_introns.txt
Introns in this gene: 9
044_TUBA1A_202_introns.txt
Introns in this gene: 3
045_MYO7A_202_introns.txt
Introns in this gene: 48
046_NUP155_201_introns.txt
Introns in this gene: 34
047_PTBP1_203_introns.txt
Introns in this gene: 14
048_U2AF2_202_introns.txt
Introns in this gene: 11
049_GAPDH_201_introns.txt
Introns in this gene: 8
050_TK1_201_introns.txt
Introns in this gene: 6
051_ATM_201_introns.txt
Introns in this gene: 62
052_TERT_201_introns.txt
Introns in this gene: 15
053_ADAMTS13_206_introns.txt
Introns in this gene: 28
054_NEU1_206_introns.txt
Introns in this gene: 5
055_GLB1_201_introns.txt
Introns in this gene: 15
056_CTSA_204_introns.txt
Introns in this gene: 14
057_ACE_202_introns.txt
Introns in this gene: 24
058_ERN1_201_introns.txt
Introns in this gene: 21
059_BMP2_201_introns.txt
Introns in this gene: 2
060_CDK6_201_introns.txt
Introns in this gene: 7
061_CCND1_201_introns.txt
Introns in this gene: 4
062_EZH1_202_introns.txt
Introns in this gene: 20
063_HAT1_201_introns.txt
Introns in this gene: 10
064_ITGB3_201_intons.txt
Introns in this gene: 14
065_IFNG_201_introns.txt
Introns in this gene: 3
066_ERBB2_219_introns.txt
Introns in this gene: 30
067_APP_201_introns.txt
Introns in this gene: 17
068_EGF_201_introns.txt
Introns in this gene: 23
069_CTNNB1_201_introns.txt
Introns in this gene: 14
070_IGF1_203_introns.txt
Introns in this gene: 5
071_MAPK1_201_introns.txt
Introns in this gene: 8
072_IGHG1_202_introns.txt
Introns in this gene: 5
073_TFRC_201_introns.txt
Introns in this gene: 18
074_CRP_201_introns.txt
Introns in this gene: 1
075_MET_201_introns.txt
Introns in this gene: 20
076_RGS6_209_introns.txt
Introns in this gene: 17
077_BCL2_202_introns.txt
Introns in this gene: 1
078_XRCC6_202_introns.txt
Introns in this gene: 12
079_RAD52_202_introns.txt
Introns in this gene: 11
080_MSTN_201_introns.txt
Introns in this gene: 2
081_PNMT_201_introns.txt
Introns in this gene: 2
082_CELF1_214_introns.txt
Introns in this gene: 15
083_BLM_201_introns.txt
Introns in this gene: 21
084_PSEN1_201_introns.txt
Introns in this gene: 11
085_AMY1A_201_introns.txt
Introns in this gene: 10
086_SPSB1_201_introns.txt
Introns in this gene: 2
087_G6PD_202_introns.txt
Introns in this gene: 12
088_USB1_201_introns.txt
Introns in this gene: 6
089_ERCC6_201_introns.txt
Introns in this gene: 20
090_ATP2B1_201_introns.txt
Introns in this gene: 19
091_ETV4_201_introns.txt
Introns in this gene: 12
092_HNF1A_201_introns.txt
Introns in this gene: 9
093_NAT2_201_introns.txt
Introns in this gene: 1
094_LPL_207_introns.txt
Introns in this gene: 9
095_FADS1_201_introns.txt
Introns in this gene: 11
096_GDF5_201_introns.txt
Introns in this gene: 1
097_ACADM_202_introns.txt
Introns in this gene: 11
098_GATA1_202_introns.txt
Introns in this gene: 5
099_GSTM1_201_introns.txt
Introns in this gene: 7
100_CDH1_201_introns.txt
Introns in this gene: 15
101_FMR1_205_introns.txt
Introns in this gene: 16
102_POLD1_205_introns.txt
Introns in this gene: 26
103_RPA1_201_introns.txt
Introns in this gene: 16
104_WRN_201_introns.txt
Introns in this gene: 34
105_SP1_201_introns.txt
Introns in this gene: 5
106_TWIST1_201_introns.txt
Introns in this gene: 1
107_TAF1_203_introns.txt
Introns in this gene: 37
108_POLB_201_introns.txt
Introns in this gene: 13
109_TERF2_201_introns.txt
Introns in this gene: 9
110_RFC1_201_introns.txt
Introns in this gene: 24
111_MAPT_204_introns.txt
Introns in this gene: 14
112_ACTA2_201_introns.txt
Introns in this gene: 8
113_PCGF2_207_introns.txt
Introns in this gene: 10
114_SSBP3_204_introns.txt
Introns in this gene: 17
115_CHEK1_213_introns.txt
Introns in this gene: 12
116_BRIP1_201_introns.txt
Introns in this gene: 19
117_NFATC2_201_introns.txt
Introns in this gene: 10
118_PRKDC_201_introns.txt
Introns in this gene: 85
119_OPTN_202_introns.txt
Introns in this gene: 14
120_ZBP1_201_introns.txt
Introns in this gene: 7
121_LDHA_205_introns.txt
Introns in this gene: 7
122_RBPJ_205_introns.txt
Introns in this gene: 11
123_FKTN_201_introns.txt
Introns in this gene: 9
124_SFPQ_201_introns.txt
Introns in this gene: 9
125_MAD1L1_201_introns.txt
Introns in this gene: 16
126_RAD51_201_introns.txt
Introns in this gene: 9
127_POLA1_201_introns.txt
Introns in this gene: 36
128_PCNA_201_introns.txt
Introns in this gene: 5
129_PRIM2_201_introns.txt
Introns in this gene: 4
130_ITPR1_203_introns.txt
Introns in this gene: 58
131_EFNA5_201_introns.txt
Introns in this gene: 4
132_OPA1_205_introns.txt
Introns in this gene: 2

**Introns total: 2007**

### Unusual introns

Order for all sequences printed out below: splice junction (8+3nt), intron start (10nt), intron end (60nt)

#### Isolating all minor (U12*) introns

Looking for U6atac* binding site motif **AT(+5C)C**

723 CTTCATTG/CTG ATATCCGTGC U6atac/ U12
TCCGGACCCCAGGCCAGTGCTCCCTCTATTCGGCACAAGCCCTTCCTTGACAGTCCCCAG unusual intron ends combination: AT-AG
735 GTTTGGAC/ATA GTATCCTATT U6atac/ U12(?) or U2 CTTAC
GAAACAAGTTAAAGAAGAATAAGATTCACCTCTGTCTGCCTTACATCCACACTACCTCAG
1141 AAAGATGT/ATA GTATCCTTTT U6atac/ U12
TTCAATGCTTTGCTAAATGTGACCAGCTAATTGGTGTTTTACCTTAACCTGTGCAAACAG
1940 TACTCCAG/TTT GTATCCACTA U6atac/ U12(?) or U2 TAAC
CAGAGTCAAGATGGTATGTTAATGTAACCAATCTGTTTCCATTTCCACTTGAATCTTTAG
[722, 734, 1140, 1939]

**Total U12 introns: 4 0.20 %**

*Looking for U12 binding site*

520 TATGTCAG/GTG GTGGGCTGGA U6/ U2, major class intron
AAACTCTGATTTTAATTGGGCTTCCTTAACAAGTCTTAATCTCTCCATGTTTTCCTTCAG
833 TTTTCAGG/GGA GTATGTACAT U6/ U2, major class intron
CTGGTCTATGAACAAAACTTTTTAAAACGATGACTGTATTTTTTCCCTTAACTCTGTTAG
1141 AAAGATGT/ATA GTATCCTTTT U6atac/ U12
TTCAATGCTTTGCTAAATGTGACCAGCTAATTGGTGTTTTACCTTAACCTGTGCAAACAG

*Caution: U12 motif CCTTAAC and U2 motif TAAC are not easy to separate, U6atac motif (see above) is a safer option*

---

**U6atac and U4atac are the names originally given to these snRNA homologues of U6 and U4, as the first minor introns identified had AT_AC ends. It later appeared that most of minor introns still have GT_AC ends and occasionally major introns can also have AT_AC ends. U2 and U1 minor homologues were always named U12 and U11.*

##### Isolating AU-AC introns

##### Testing for the first intron G

723 CTTCATTG/CTG ATATCCGTGC U6atac/ U12
TCCGGACCCCAGGCCAGTGCTCCCTCTATTCGGCACAAGCCCTTCCTTGACAGTCCCCAG unusual intron ends combination: AT-AG

##### Testing for the last intron G

All the examined introns end with a G

#### List S2. Minor introns

*Minor (U12) introns*

| # | File | Gene, (ensembl isoform) | Intron | In. bp | Ch |
|---|---|---|---|---|---|
| 723 | 045_MYO7A_202_introns | *MYO7A*, myosin VIIA (202) | 38 of 48 | 858 | 11 |
| 735 | 046_NUP155_201_introns | *NUP155*, nucleoporin 155 (201) | 2 of 34 | 262 | 5 |
| 1141 | 071_MAPK1_201_introns | *MAPK1*, mitogen-activated protein kinase 1 (201) | 2 of 8 | 604 | 22 |
| 1940 | 130_ITPR1_203_introns | *ITPR1,* inositol 1,4,5-trisphosphate receptor type 1 (203) | 24 of 58 | 3751 | 3 |

*GA-AG intron*

| | | | | | |
|---|---|---|---|---|---|
| 1942 | 130_ITPR1_203_introns | *ITPR1,* inositol 1,4,5-trisphosphate receptor type 1(203) | 26 of 58 | 1967 | 3 |

**Excluding the minor spliceosome (U12) introns from the analysis**
**Total splice junctions of major introns only: 2003**

**Isolating GC-AG introns**

#### List S3. Major introns with substitutions of +2U: GC(A)_AG introns

Order for all sequences printed out below: splice junction (8+3nt), intron start (10nt), intron end (60nt)

30 AAGCCCAGAAA GCAAGTACAT
CTTGGAAAGTTAGTTGTTCTTTGTAGAGCATGCTGACTAATAATGCTATCCTCCCAACAG
84 TTTGCAAGCTG GCAAGAAACT
TTTCAATCAACCATAATTTCTTCTCTTGAGTTATTTCATTGTCTTTCTGTCCTAACTCAG
489 GCTATAAGGTG GCACGTAGCA
TGTCCTTTTATTCTTAGATACCTCCTTCACTGAGACCTTTTCCTTACCTCACCTCTCTAG
645 GTGGACAGGTC GCGCGTATAC
GGGGACAGGGAGCTCAGGGAGTGGCAGCTGCCCGGGGCCGACAGCTCCTGTTCCCTGCAG
674 CCTCGTGGTTC GCAAGTTGGG
TTATCCCCTGCCCAGGACTGAGGCTGGCCTGTGTGTTTGGGACTTGTGGGGTCCCCACAG
856 TGATGAAGAGA GCAAGTGTTA
AGATGTGAGAATATTTGAAATACCTTGTTTCTTAATTTTGTGTCTTTTTTTTAATGGTAG
883 CAGGACAGCCC GCAAGTGTGG
AGGGTTGGAGGATGCCACCTCTGGCCTCTTCTGGAACGGAGTCTGATTTTGGCCCCGCAG
1016 AAGCACAGATT GCAAGAAGGG
CCTGCATTTGCAGAGCTCCCTAGGGCCGTGACTGAGACCTGTCCTTCCTGTCTTTTTCAG
1320 AATTACAGGTT GCAAGTGCTC
GCAGTTGATGACTGTATATGCTCTTTCATGGTTGTTTTCTTCTTGGACGTTTGGATGCAG
1643 GGTGCCAGGGA GCAAGTCTGG
CCTCCTGGGTCTCCTGGTTCTTCCTGGGTCTTCCTAGCTTCATTCTTTCTCCGTCCCCAG
1676 CCAGACAGTTA GCAAGTCGGA
GTTCTCTGGTTTTCTTTGTAAGGCGTGTCTCAATATTTGACATTTTCATTTCTGTTGTAG
1942 GGAGGAAGTGA GAAAGTATAT **(?)**
GGGGCGTGAGAGGAGGCATTTGTCATTCATTTGGCCTTTCCCCACCTTGTGCTCCTTTAG
1961 AACTGAAGAAC GCAAGTAGGA
TTCAGGCCGGGCCCCCAAGCAGCCCCTCTACATTTCCTCTCCCTTTGTTTCTCCAACTAG
[29, 83, 488, 644, 673, 855, 882, 1015, 1319, 1642, 1675, 1941, 1960]

**Total GC(A)_AG introns: 13 0.65 %**
**All GC(A)-AC introns have multiple W-C pairs in U5 and U6 helices at the ex/in boundary.**

#### Isolating the exon/intron boundaries missing both conserved Gs: Exon-end -1Gsub and intron +5G sub

**List S4. Introns missing both conserved exon-end -1G and intron +5G**
Sequences of exon junctions (8nt/3nt), intron starts (10nt) and intron ends (60nt). Substitutions of the conserved guanines are in **red**. **Bold and underlined** are positions that form Watson-Crick pairs with U5 snRNA (exon junctions) and U6 snRNA (intron starts). In **bold not underlined** are the conserved adenines at intron positions +3 and +4, that form non-canonical pairs with U6 positions 43 and 44. ID numbers refer to lists `splice_junctions_u6, intron_starts_u6, introns_ends_u6` (numbers in brackets refer to lists `splice_junctions, intron_starts, introns_ends` before the data cleaning, which allow to trace the gene and intron number).

899 (908) GCAGTCCA/**G**T**G** GTT**A**T**G**TC**C**T *compensation with +9C=G pair?*
GAGCCTGCCCTGCTGGGAATCGGGGAAGCAC**T**G**CT**T**AC**CTGTCTCCTGCTCCCTTTT**C**AG *[very strong 3' end interactions]*

*Compensation with multiple other W-C pairs of the U5 and U6 binding sites*
984 (993) **A**AAG**AAA**A/**G**T**G** GT**AA**C**T**TCGC
AGGAAGGCTTCTGGGCTGTCTGATTGCACTTTCTTCTTATCCTCCCGTCTCCTCCTTTAG
1220 (1231) **A**T**TAA**TCA/AGA GT**AA**A**TA**C**C**T
TGCTTTCTGTGAGAATACTTCATTATTTGCTGATGACGCACTTTTCCTTGACCTCTGTAG
1278 (1289) TACTT**AA**A/**G**AA GT**AA**A**TA**AAT
AGTTACTCTCAAACTATTGTGAAATGATACATCAACGTATATCTTATGTTTCAAAAATAG
1370 (1382) CAG**AAAA**A/**G**GT GT**AA**T**TA**AAT
TTTGTGAACAGTGCTTTTGATTGTTCTACATGGCATATTCACATCCATTTTCTTCCACAG
1714 (1728) **A**AACT**AA**A/**G**TT GT**A**CT**T**C**T**AT
AATATTCTCTTGTCATCAACGTGATAACTTAAACATTGCTGACCTCCTGGTCTGTTTTAG

**Total exon/intron boundaries missing both conserved Gs: 6 0.30 %**
>>>

### Human mutation data explained by the U5 hypothesis

Fu et al., 2011 Mutations of exon-start guanine: $G_{+1}$ → *T* or $G_{+1}$ → *A*

**List S5. Mutations of exon-start guanine: $G_{+1}$ → *T* or $G_{+1}$ → *A*** (Fu et al., 2011)

| Gene | Ex | PSI | U5 base pairs: **Watson-Crick** Isosteric **Non-isosteric** | U5 model support | U2 base pairs: **Watson-Crick (BP, BP+$C_{-3}$)** Isosteric **Non-isosteric** | PPT Stretch (PPS) |
|---|---|---|---|---|---|---|
| *GH1* | 3 | 1 | *T*AA | Yes | cgtagA$_{-26}$cc… t$_{-3}$ (1,1) | 10 |
| *FECH* | 9 | 17 | *T*TT | Yes | cacttA$_{-25}$cg…t$_{-3}$ (4,4) | 6 |
| *EYA1* | 10 | 66 | *T*AT | Yes | tcttcA$_{-22}$cc…t$_{-3}$ (3,3) | 5 |
| *LPL* | 5 | 100 | *T*CC | Yes | aatttA$_{-34}$ca…a$_{-3}$ (4,4) | 14 |
| *HEXA* | 13 | 100 | *A*CC | Yes | gcccaA$_{-27}$tc…c$_{-3}$ (2,3) | 13 |
| *PKHD1* | 25 | 99<br>0* | upstream **cryptic** ag**ACG**<br>instead of *T*TC | Yes | aaaaaA$_{-20}$ct…t$_{-3}$ (3,3)<br>atgtaA$_{-10}$cc…c$_{-3}$ (3,4) | 6<br>3 |
| *COL1A2* | 37 | 41 | *T*GT | Yes | ctgtaA$_{-18}$ct…c$_{-3}$ (3,4) | 5 |
| *CLCN2* | 19 | 81 | *A*GA | No⟶ | tgggaA$_{-6}$ca…c$_{-3}$ (4,5) | 4 |
| *LAMA2* | 24 | 100 | *A*TG | Yes | tataaA$_{-28}$ct…c$_{-3}$ (4,5) | 9 |
| *CAPN3* | 10 | 91 | *T*CT | Yes⟶ | ctgtgA$_{-31}$cc…a$_{-3}$ (2,2) | 10 |
| *CAPN3* | 17 | 0 | *T*TT | Yes | cattcA$_{-36}$ca…a$_{-3}$ (4,4) | 9 |
| *NEU1* | 2 | 100 | *A*TG | Yes | tgttgA$_{-22}$cc…c$_{-3}$ (3,4) | 14 |
| *COL6A2* | 8 | 100 | *T*GC | No⟶ | cactaA$_{-26}$tg…c$_{-3}$ (4,5) | 15 |
| *COL1A1* | 23 | 100 | *A*GC | No⟶ | *cct*ctA$_{-27}$tc…t$_{-3}$ (0,0)** | 16 |

* 0% PSI for *PKHD1* exon 25. Longer exon with cryptic 3'ss is included with PSI 99% and exon skipping amounts for 1%. We used alternatively PSI 0 or PSI 99 for the plots, Welch's ANOVA and Kruskal-Wallis test.

>PKHD1-202 intron 24:protein_coding

GTATGTATAGTATCCCTTCTAGTGGCATGGATTTGCTGCCCTGGAGAAATTCTGCAAAAAAAAAAATCAGA
ATTATACAAATGGAAACTGTCATGAATTATTAATTTTGTTGGAATATTTGAGTAAAGGGATTCTATGAACTTT
CTTTCTGGAAGTTTTCAACACATATTTTGTATATCATATCCCTGAAAAAGCAATTTAAACCCAACATCTCCC
TTCCTAACTCAGTGTAGGTAGTGAAGTTATTGGTCAGTGTCCACCATGCAATAATTCTACATCTGTTATTG
GATAATTTAAGCAGATGTGATTGACTTAGATGATTGTGGAAGCAGATGGCATAGCAGAAGGAATTTTAGAA
AAGAATTTGCTGTCTTTGGGTAATGACAAAACCAGATGCTCTTGCATATTTAGGGTCTTAAAAAATAATCTA
ATGATCAACTAGCATTTGGTTATTATAAAATGAAGCCCAAGATGAAAATTTTTCCTAGTTGTATGGTGTTGT
GGGTTATCTGGTTTTCAATCGAAATTCTAGAAAGGAGATGGAGGAAGAATGGGAGACCTTGCTCTTATCT
CATGGCACTGTTTCTTTCCACTTTATTTTAATATTTTGGATTGCAAACATATAAAATGGAAAAGCATGGAGA
AGCAAAGTGTCTCAAGACTTAGAGAGAAGATTGATTTATTGTACCTCTTTTGGCAATCCACTTGCAAATGA
ATATTTCTCTAATCTACTGGGACTTTCCTCAGCCATTATTTACTGTTACGTTTCTATATTTCAATTCTCTGTT
ACTCTGCTTTCACTTTCTAGCTGTCTGATTATCTGAAACACAATGTG**AAAAAC**TCTTTA**T**TATTTGATTAG|
BP PPS Cryptic 3'ss

**ACG**AGATTAGATTTCGGTTCCATGACAGAATTTACCAGAA**ATGTAAC**CATCT**C**AG

** *COL1A1* intron 22 is only 113bp long, it has only 1 conserved major intron position (+5G), however, it has 5 conserved minor intron positions: +6C and +7T and branchpoint helix CCT. There is also a super perfect PPT: an uninterrupted stretch of 16 uridines. As it is likely a U12 intron we also tested excluding this data point.

>COL1A2-201 intron 22:protein_coding

GTAA**G***CT*GTCTATCACTTACTTCCTAGAAAGGGGCTTGCTGCTTCTGGTGGTGGGTGTGTCATTAGCTTTA
GCATCCTCCT*CCT*CT**A**TCTGTTTTTTTTTTTTTTTTGAATAG
BP PPS

Can we state that the presence of +2C or/and +3G that form Watson-Crick pairs with U5 Loop1 according to our new model is a strong factor that promotes exon inclusion in spite of +1G mutations? We explore the other factors that are expected to influence PSI:

*The effect of the exon-start +1G mutations on the exon inclusion efficiency (PSI)*

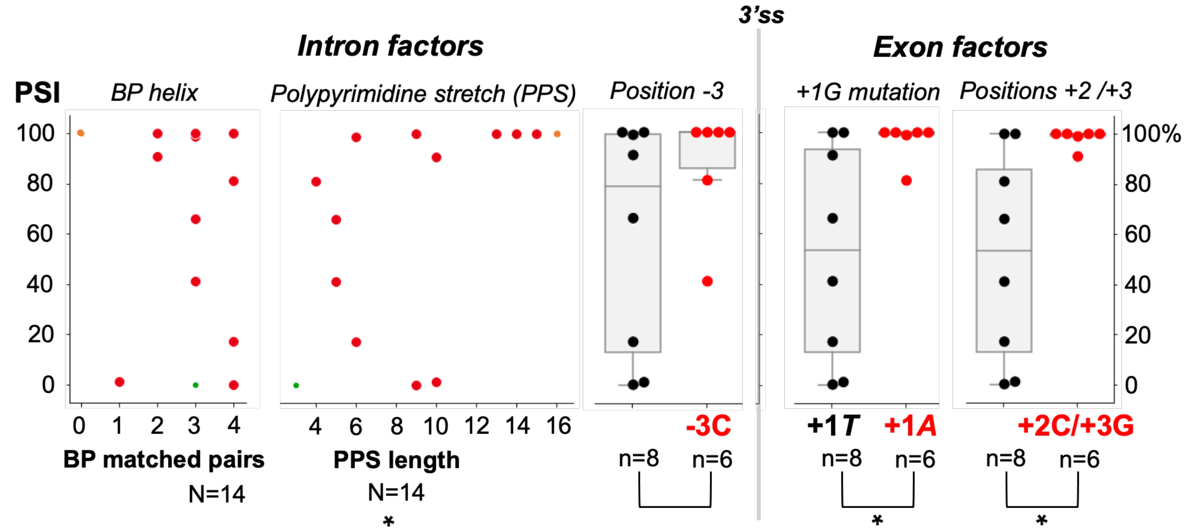


| | BP matched pairs | PPS length | | -3C | +1T vs +1A | +2C/+3G |
|---|---|---|---|---|---|---|
| *Spearman's correlation* | ρ=–0.05, P=8.7e-01 | * ρ=0.64, **P=1.4e-02** | *Welch's ANOVA* | $F_{0.09}$=2.2, P = 1.7e-01 | * $F_{0.13}$=8.3, **P = 2.2e-02** | * $F_{0.14}$=10.2, **P = 1.5e-02** |
| | | | *Kruskal-Wallis* | $H_1$=2.18, P = 1.4e-01 | $H_1$=4.06, **P = 4.4e-02** | $H_1$=4.62, **P = 3.2e-02** |
| **Excluding U12 datapoint •** | | | | | | |
| *Spearman's correlation* | ρ=0.09, P=7.6e-01 | ρ=0.58, **P=3.9e-02** | *Welch's ANOVA* | $F_{0.11}$= 2.8, P = 1.3e-01 | $F_{0.13}$=7.9, **P = 2.3e-02** | $F_{0.16}$=13.0, **P = 1.1e-02** |
| | | | *Kruskal-Wallis* | $H_1$=3.11, P = 7.8e-02 | $H_1$=3.27, P = 7.1e-02 | $H_1$=6.24, **P = 1.2e-02** |

**Figure S11|** Inclusion of exon affected by +1G mutation (data from Fu et al., 2011) is influenced by multiple *cis* factors. In agreement with Fu and colleagues PPS length comes up as a significant intronic factor (PPS, polypyrimidine stretch, uninterrupted by purines, as opposed to PPT, polypyrimidine tract, used generally in this article, for variable sequence, not exclusively pyrimidines, between the branchpoint and intron position -3). We add that the exon-start sequence is also a strong factor of exon inclusion. Nine of these 14 mutations were +1G → *T* with PSI ranging 0-100% and five were +1G → *A* with consistently high PSI 81-100%. Given that exon-start +1G is conserved at 50% in humans, we can expect that a change for purine is better tolerated (generally +1A is twice more likely than +1T). Specifically, we looked for cytosines in exon position +2 and guanines in exon position +3 that form Watson-Crick pairs with U5 Loop1 according to our new model. Indeed, all +1G mutations followed by +2C/+3G have PSI above 90%, which suggests a compensatory mechanism in agreement with our new U5 binding register. **PSI:** percent spliced-in

**Excluding U12 datapoint** • in scatterplots refers to +1G mutation in *COL1A1* exon 23

Unusual *COL1A1* intron 22 is only 113bp long, it has only 1 conserved major intron position (+5G), however, it has 5 conserved minor intron positions: +6C and +7T and branchpoint motif CCT. There is also a super perfect PPT: an uninterrupted stretch of 16 uridines. As it is likely a U12 intron we also tested excluding this data point.

>COL1A2-201 intron 22:protein_coding

GTAA**G***CT*GTCTATCACTTACTTCCTAGAAAGGGGCTTGCTGCTTCTGGTGGTGGGTGTGTCATTAGCTTTA

GCATCCTCCT*CCT*CT**A**TCTGTTTTTTTTTTTTTTTTGAATAG

BP PPS

• datapoint in scatterplots refers to *PKHD* exon 25 PSI 0, not included in the boxplots. *PKHD* cryptic 3'ss (longer exon 25) with PSI 99 appears both in scatterplots and in the boxplots and was used exclusively for statistical tests presented here (see below pages 23-32 for all alternative calculations).

**Table S4.** Factors that promote efficient exon inclusion (PSI 81-100%) in spite of exon-start (+1G) mutations (red dots on the boxplots **Figure S11**). Exon inclusion for five +1G mutations (grey background) are supported by at least two factors out of +2C/+3G, +1G→*A* and -3C. *COL1A1* exon 23 is associated with U12 intron 22 and can be excluded from the analysis (see legend to **Figure S**). Therefore, +1G→*A* does not come up as an independent factor. *LPL* exon 5 and *COL6A2* exon 8 are both additionally supported by 4 matched (Watson-Crick) pairs in the branchpoint helix and a long PPS. The high rate of *CAPN3* exon 10 inclusion is left to be explained solely by +2C/+3G. (Key: *GENE* exon: BP matched pairs PPS length)

| **Exon +2C/+3G** | **Exon +1G→*A*** | **Intron -3C** |
|---|---|---|
| **PSI 100** | | |
| *HEXA* exon 13: 2 **13** | *HEXA* exon 13: 2 **13** | *HEXA* exon 13: 2 **13** |
| *LAMA* exon 24: **4** 9 | *LAMA* exon 24: **4** 9 | *LAMA* exon 24: **4** 9 |
| *NEU1* exon 10: 3 **14** | *NEU1* exon 10: 3 **14** | *NEU1* exon 10: 3 **14** |
| *LPL* exon 5: **4** **14** | | |
| | *COL1A1* exon 23: **0*** **16** | |
| | | *COL6A2* exon 8: **4** **15** |
| **PSI 99** | | |
| *PKHD* **cryptic** ex. 25: 3 6 | *PKHD* **cryptic** ex. 25: 3 6 | |
| **PSI 91** | | |
| *CAPN3* exon 10: 2 10 | | |
| | **PSI 81** | |
| | *CLNC2* exon 19: **4** 4 | *CLNC2* exon 19: **4** 4 |

Kruskal-Wallis rank sum test and **Welch's** ANOVA (t-test) for +2C/+3G

1. *Include COL1A1 and PKHD normal 3'ss PSI 0 (N=14)*

| | PSI |
|---|---|
| U5 Watson-Crick pairs | *LPL:* 100, *HEXA:* 100, *LAMA2:* 100, *CAPN3*ex10: 91, *NEU1:* 100 |
| No U5 Watson-Crick pairs | *GH1:* 1, *FECH:* 17, *EYA1:* 66, *COL1A2:* 41, *CLCN2:* 81, *COL6A2:* 100, *COL1A1:* 100, *CAPN3*ex17: 0, *PKHD1:* 0 |

```
U5_WatsonCrick=[100,100,100,91,100]#n=5
U5_noWatsonCrick=[1,17,66,41,81,100,100,0,0]#n=9
from scipy import stats
###Kruskal-Wallis rank sum test and Welch's t-test for +2C/+3G
print(stats.kruskal(U5_WatsonCrick, U5_noWatsonCrick))
print(stats.ttest_ind(U5_WatsonCrick, U5_noWatsonCrick, equal_var = False))
KruskalResult(statistic=5.255847255369932, pvalue=0.021873149625322268)
Ttest_indResult(statistic=3.7021033170416677, pvalue=0.005702540534091117)
```

2. *Include COL1A1 and PKHD cryptic 3'ss PSI 99 (N=14)*

| | PSI |
|---|---|
| U5 Watson-Crick pairs | *LPL:* 100, *HEXA:* 100, *LAMA2:* 100, *CAPN3*ex10: 91, *NEU1:* 100, *PKHD1:* 99 |
| No U5 Watson-Crick pairs | *GH1:* 1, *FECH:* 17, *EYA1:* 66, *COL1A2:* 41, *CLCN2:* 81, *COL6A2:* 100, *COL1A1:* 100, *CAPN3*ex17: 0 |

```
U5_WatsonCrick=[100,100,100,91,100,99]#n=6
U5_noWatsonCrick=[1,17,66,41,81,100,100,0]#n=8
KruskalResult(statistic=4.622222222222221, pvalue=0.03156032503831624)
Ttest_indResult(statistic=3.196999122936383, pvalue=0.014734923368321556)
```

3. *Include PKHD normal 3'ss PSI 0 and exclude COL1A1 (N=13)*

| | PSI |
|---|---|
| U5 Watson-Crick pairs | *LPL:* 100, *HEXA:* 100, *LAMA2:* 100, *CAPN3*ex10: 91, *NEU1:* 100 |
| No U5 Watson-Crick pairs | *GH1:* 1, *FECH:* 17, *EYA1:* 66, *COL1A2:* 41, *CLCN2:* 81, *COL6A2:* 100, *CAPN3*ex17: 0, *PKHD1:* 0 |

```
U5_WatsonCrick=[100,100,100,91,100]#n=5
U5_noWatsonCrick=[1,17,66,41,81,100,0,0] #n=8
KruskalResult(statistic=6.57201166180758, pvalue=0.01035947924583537)
Ttest_indResult(statistic=4.208231542954459, pvalue=0.0037207957451188445)
```

4. *Include PKHD cryptic 3'ss PSI 99 and exclude COL1A1 (N=13)*

| | PSI |
|---|---|
| U5 Watson-Crick pairs | *LPL:* 100, *HEXA:* 100, *LAMA2:* 100, *CAPN3*ex10: 91, *NEU1:* 100, *PKHD1:* 99 |
| No U5 Watson-Crick pairs | *GH1:* 1, *FECH:* 17, *EYA1:* 66, *COL1A2:* 41, *CLCN2:* 81, *COL6A2:* 100, *CAPN3*ex17: 0 |

```
U5_WatsonCrick=[100,100,100,91,100,99]#n=6
U5_noWatsonCrick=[1,17,66,41,81,100,0] #n=7
KruskalResult(statistic=6.240863787375417, pvalue=0.012483557594188328)
Ttest_indResult(statistic=3.6121657705778767, pvalue=0.010840625573149486)
```

**R:** Kruskal-Wallis rank sum test and **Welch's** ANOVA (t-test) for **+2C/+3G**

##### 1. *Include COL1A1 and PKHD normal 3'ss PSI 0 (N=14)*

```
### Calculations from lists (unequal sizes are acceptable)
U5_WatsonCrick=c(100,100,100,91,100)#n=5
U5_noWatsonCrick=c(1,17,66,41,81,100,100,0,0)#n=9

> kruskal.test(list(U5_WatsonCrick,U5_noWatsonCrick))#uses vectors without NaNs

	Kruskal-Wallis rank sum test

data:  list(U5_WatsonCrick, U5_noWatsonCrick)
Kruskal-Wallis chi-squared = 5.2558, df = 1, p-value = 0.02187

> shapiro.test(U5_WatsonCrick)$p.value #uses vector without NaNs
[1] 0.0001309782
> shapiro.test(U5_noWatsonCrick)$p.value #uses vector without NaNs
[1] 0.09080469
> sd(U5_WatsonCrick) #uses vector without NaNs
[1] 4.024922
> sd(U5_noWatsonCrick) #uses vector without NaNs
[1] 42.68034
> (sd(U5_WatsonCrick))^2 #uses vector without NaNs
[1] 16.2
> (sd(U5_noWatsonCrick))^2 #uses vector without NaNs
[1] 1821.611

> t.test(U5_WatsonCrick,U5_noWatsonCrick,var.equal = FALSE)# vectors without NaNs

	Welch Two Sample t-test

data:  U5_WatsonCrick and U5_noWatsonCrick
t = 3.7021, df = 8.2539, p-value = 0.005703
alternative hypothesis: true difference in means is not equal to 0
95 percent confidence interval:
 20.19659 85.98118
sample estimates:
mean of x mean of y
 98.20000  45.11111

###Dataframe (requires Nans for unequal size groups)
U5_WatsonCrick=c(100,100,100,91,100,NaN,NaN,NaN,NaN)#n=5
U5_noWatsonCrick=c(1,17,66,41,81,100,100,0,0)#n=9
U5_table=data.frame(cbind(U5_WatsonCrick,U5_noWatsonCrick))# vectors with NaNs
U5_table_stacked=stack(U5_table)

#kruskal.test and t.test produce the same result as with lists without NaNs
kruskal.test(values~ind,U5_table_stacked)#uses df with NaNs
t.test(values~ind,U5_table_stacked, var.equal = FALSE)#uses df with NaNs

> oneway.test(values~ind,U5_table_stacked, var.equal = FALSE)#uses df with NaNs

	One-way analysis of means (not assuming equal variances)

data:  values and ind
```

**F = 13.706**, num df = 1.0000, denom df = 8.2539, **p-value = 0.005703**

2. ***Include COL1A1 and PKHD cryptic 3'ss PSI 99 (N=14)***

```
### Calculations from lists (unequal sizes are acceptable)
U5_WatsonCrick=c(100,100,100,91,100,99)#n=6
U5_noWatsonCrick=c(1,17,66,41,81,100,100,0)#n=8

> kruskal.test(list(U5_WatsonCrick,U5_noWatsonCrick))#uses vectors without NaNs

	Kruskal-Wallis rank sum test

data:  list(U5_WatsonCrick, U5_noWatsonCrick)
Kruskal-Wallis chi-squared = 4.6222, df = 1, p-value = 0.03156

> shapiro.test(U5_WatsonCrick)$p.value #uses vector without NaNs
[1] 0.0001621368
> shapiro.test(U5_noWatsonCrick)$p.value #uses vector without NaNs
[1] 0.2241062
> sd(U5_WatsonCrick) #uses vector without NaNs
[1] 3.614784
> sd(U5_noWatsonCrick) #uses vector without NaNs
[1] 41.89016
> (sd(U5_WatsonCrick))^2 #uses vector without NaNs
[1] 13.06667
> (sd(U5_noWatsonCrick))^2 #uses vector without NaNs
[1] 1754.786
> t.test(U5_WatsonCrick,U5_noWatsonCrick,var.equal = FALSE)# vectors without NaNs

	Welch Two Sample t-test

data:  U5_WatsonCrick and U5_noWatsonCrick
t = 3.197, df = 7.1387, p-value = 0.01473
alternative hypothesis: true difference in means is not equal to 0
95 percent confidence interval:
 12.52701 82.63966
sample estimates:
mean of x mean of y 
 98.33333  50.75000 

###Dataframe (requires Nans for unequal size groups)
U5_WatsonCrick=c(100,100,100,91,100,99,NaN,NaN)#n=6
U5_noWatsonCrick=c(1,17,66,41,81,100,100,0)#n=8
U5_table=data.frame(cbind(U5_WatsonCrick,U5_noWatsonCrick))# vectors with NaNs
U5_table_stacked=stack(U5_table)

#kruskal.test and t.test produce the same result as with lists without NaNs
kruskal.test(values~ind,U5_table_stacked)#uses df with NaNs
t.test(values~ind,U5_table_stacked, var.equal = FALSE)#uses df with NaNs

> oneway.test(values~ind,U5_table_stacked, var.equal = FALSE)#uses df with NaNs

	One-way analysis of means (not assuming equal variances)

data:  values and ind
F = 10.221, num df = 1.0000, denom df = 7.1387, p-value = 0.01473
```

##### *4.Include COL1A1 and PKHD cryptic 3’ss PSI 99 (N=14)*

```
### Calculations from lists (unequal sizes are acceptable)
U5_WatsonCrick=c(100,100,100,91,100,99)#n=6
U5_noWatsonCrick=c(1,17,66,41,81,100,0) #n=7

> kruskal.test(list(U5_WatsonCrick,U5_noWatsonCrick))#uses vectors without NaNs

      Kruskal-Wallis rank sum test

data:  list(U5_WatsonCrick, U5_noWatsonCrick)
Kruskal-Wallis chi-squared = 6.2409, df = 1, p-value = 0.01248

> shapiro.test(U5_WatsonCrick)$p.value #uses vector without NaNs
[1] 0.0001621368
> shapiro.test(U5_noWatsonCrick)$p.value #uses vector without NaNs
[1] 0.4577796
> sd(U5_WatsonCrick) #uses vector without NaNs
[1] 3.614784
> sd(U5_noWatsonCrick) #uses vector without NaNs
[1] 39.81505
> (sd(U5_WatsonCrick))^2 #uses vector without NaNs
[1] 13.06667
> (sd(U5_noWatsonCrick))^2 #uses vector without NaNs
[1] 1585.238
> t.test(U5_WatsonCrick,U5_noWatsonCrick,var.equal = FALSE)#uses vectors without
NaNs

      Welch Two Sample t-test

data:  U5_WatsonCrick and U5_noWatsonCrick
t = 3.6122, df = 6.1153, p-value = 0.01084
alternative hypothesis: true difference in means is not equal to 0
95 percent confidence interval:
 17.78805 91.45005
sample estimates:
mean of x mean of y
 98.33333  43.71429

###Dataframe (requires Nans for unequal size groups)
U5_WatsonCrick=c(100,100,100,91,100,99,NaN)#n=6
U5_noWatsonCrick=c(1,17,66,41,81,100,0) #n=7
U5_table=data.frame(cbind(U5_WatsonCrick,U5_noWatsonCrick))# vectors with NaNs
U5_table_stacked=stack(U5_table)

#kruskal.test and t.test produce the same result as with lists without NaNs
kruskal.test(values~ind,U5_table_stacked)#uses df with NaNs
t.test(values~ind,U5_table_stacked, var.equal = FALSE)#uses df with NaNs

> oneway.test(values~ind,U5_table_stacked, var.equal = FALSE)#uses df with NaNs

      One-way analysis of means (not assuming equal variances)

data:  values and ind
F = 13.048, num df = 1.0000, denom df = 6.1153, p-value = 0.01084
```

Welch's ANOVA and Kruskal-Wallis rank sum test for substitute *A* or *T*

1. *Include COL1A1 and PKHD normal 3'ss PSI 0 (N=14)*

| | PSI |
|---|---|
| *A* | *HEXA:* 100, *LAMA2:* 100, *CLCN2:* 81, *NEU1:* 100, *COL1A1:* 100 |
| *T* | *GH1:* 1, *FECH:* 17, *EYA1:* 66, *LPL:* 100, *COL1A2:* 41, *CAPN3*ex10: 91, *COL6A2:* 100, *CAPN3*ex17: 0, *PKHD1:* 0 |

```
a=[100,100,81,100,100]#n=5
t=[1,17,66,100,41,91,100,0,0]#n=9
KruskalResult(statistic=4.638080084858127, pvalue=0.031269966652780735)
Ttest_indResult(statistic=3.3095330554652915, pvalue=0.009041709556861968)
```

2. *Include COL1A1 and PKHD cryptic 3'ss PSI 99 (N=14)*

| | PSI |
|---|---|
| *A* | *HEXA:* 100, *LAMA2:* 100, *CLCN2:* 81, *NEU1:* 100, *COL1A1:* 100, *PKHD1:* 99 |
| *T* | *GH1:* 1, *FECH:* 17, *EYA1:* 66, *LPL:* 100, *COL1A2:* 41, *CAPN3*ex10: 91, *COL6A2:* 100, *CAPN3*ex17: 0 |

```
a=[100,100,81,100,100,99]#n=6
t=[1,17,66,100,41,91,100,0]#n=8
KruskalResult(statistic=4.0625, pvalue=0.04384554166017646)
Ttest_indResult(statistic=2.873903456290757, pvalue=0.02188473533338844)
```

3. *Include PKHD normal 3'ss PSI 0 and exclude COL1A1 (N=13)*

| | PSI |
|---|---|
| *A* | *HEXA:* 100, *LAMA2:* 100, *CLCN2:* 81, *NEU1:* 100 |
| *T* | *GH1:* 1, *FECH:* 17, *EYA1:* 66, *LPL:* 100, *COL1A2:* 41, *CAPN3*ex10: 91, *COL6A2:* 100, *CAPN3*ex17: 0, *PKHD1:* 0 |

```
a=[100,100,81,100]#n=4
t=[1,17,66,100,41,91,100,0,0]#n=9
KruskalResult(statistic=3.6384839650145793, pvalue=0.05645832236429441)
Ttest_indResult(statistic=3.1903049635301564, pvalue=0.010284290272625068)
```

4. *Include PKHD cryptic 3'ss PSI 99 and exclude COL1A1 (N=13)*

| | PSI |
|---|---|
| *A* | *HEXA:* 100, *LAMA2:* 100, *CLCN2:* 81, *NEU1:* 100, *PKHD1:* 99 |
| *T* | *GH1:* 1, *FECH:* 17, *EYA1:* 66, *LPL:* 100, *COL1A2:* 41, *CAPN3*ex10: 91, *COL6A2:* 100, *CAPN3*ex17: 0 |

```
a=[100,100,81,100,99]#n=5
t=[1,17,66,100,41,91,100,0]#n=8
KruskalResult(statistic=3.2651162790697636, pvalue=0.07076800545138776)
Ttest_indResult(statistic=2.806397062740887, pvalue=0.02347286415692591З)
```

2. *Include COL1A1 and PKHD cryptic 3'ss PSI 99 (N=14)*

```
### Calculations from lists (unequal sizes are acceptable)
a=c(100,100,81,100,100,99)#n=6
t=c(1,17,66,100,41,91,100,0)#n=8

> kruskal.test(list(a,t))#uses vectors without NaNs

	Kruskal-Wallis rank sum test

data:  list(a, t)
Kruskal-Wallis chi-squared = 4.0625, df = 1, p-value = 0.04385

> shapiro.test(a)$p.value
[1] 5.761099e-05
> shapiro.test(t)$p.value
[1] 0.1474304
> sd(a)
[1] 7.685484
> sd(t)
[1] 43.05478
> (sd(a))^2
[1] 59.06667
> (sd(t))^2
[1] 1853.714

> t.test(a,t,var.equal = FALSE)# vectors without NaNs

	Welch Two Sample t-test

data:  a and t
t = 2.8739, df = 7.5883, p-value = 0.02188
alternative hypothesis: true difference in means is not equal to 0
95 percent confidence interval:
  8.485037 80.848297
sample estimates:
mean of x mean of y
 96.66667  52.00000

###Dataframe (requires NaNs added to smaller groups to make sizes equal)
a=c(100,100,81,100,100,99,NaN,NaN)#n=6
t=c(1,17,66,100,41,91,100,0)#n=8
U5_table=data.frame(cbind(a,t))
U5_table_stacked=stack(U5_table)

#kruskal.test and t.test produce the same result as with lists without NaNs
kruskal.test(values~ind,U5_table_stacked)#uses df with NaNs
t.test(values~ind,U5_table_stacked, var.equal = FALSE)#uses df with NaNs

> oneway.test(values~ind,U5_table_stacked, var.equal = FALSE)#uses df with NaNs

	One-way analysis of means (not assuming equal variances)

data:  values and ind
F = 8.2593, num df = 1.0000, denom df = 7.5883, p-value =0.02188
```

4. *Include PKHD cryptic 3'ss PSI 99 and exclude COL1A1 (N=13)*

```
### Calculations from lists (unequal sizes are acceptable)
a=c(100,100,81,100,99)#n=5
t=c(1,17,66,100,41,91,100,0) #n=8

> kruskal.test(list(a,t))#uses vectors without NaNs

	Kruskal-Wallis rank sum test

data:  list(a, t)
Kruskal-Wallis chi-squared = 3.2651, df = 1, p-value = 0.07077

> shapiro.test(a)$p.value
[1] 0.0004215357
> shapiro.test(t)$p.value
[1] 0.1474304
> sd(a)
[1] 8.396428
> sd(t)
[1] 43.05478
> (sd(a))^2
[1] 70.5
> (sd(t))^2
[1] 1853.714

> t.test(a,t,var.equal = FALSE)#uses vectors without NaNs

	Welch Two Sample t-test

data:  a and t
t = 2.8064, df = 7.8271, p-value = 0.02347
alternative hypothesis: true difference in means is not equal to 0
95 percent confidence interval:
  7.705914 80.294086
sample estimates:
mean of x mean of y 
       96        52 

###Dataframe (requires NaNs added to smaller groups to make sizes equal)
a=c(100,100,81,100,99,NaN,NaN,NaN)#n=5
t=c(1,17,66,100,41,91,100,0) #n=8
U5_table=data.frame(cbind(a,t))
U5_table_stacked=stack(U5_table)

#kruskal.test and t.test produce the same result as with lists without NaNs
kruskal.test(values~ind,U5_table_stacked)#uses df with NaNs
t.test(values~ind,U5_table_stacked, var.equal = FALSE)#uses df with NaNs

> oneway.test(values~ind,U5_table_stacked, var.equal = FALSE)#uses df with NaNs

	One-way analysis of means (not assuming equal variances)

data:  values and ind
F = 7.8759, num df = 1.0000, denom df = 7.8271, p-value = 0.02347
```

Welch's ANOVA and Kruskal-Wallis rank sum test for -3C

1. *Include COL1A1 and PKHD normal 3'ss PSI 0 (N=14)*

| | PSI |
|---|---|
| -3C | *HEXA:* 100, *COL1A2:* 41, *CLCN2:* 81, *LAMA2:* 100, *NEU1:* 100, *COL6A2:* 100, *PKHD1:* 0 |
| -3Csub | *GH1:* 1, *FECH:* 17, *EYA1:* 66, *LPL:* 100, *CAPN3*ex10: 91, *COL1A1:* 100, *CAPN3*ex17: 0, |

```
pos3C=[100,41,81,100,100,100,0]#n=7
pos3Csub=[1,17,66,100,91,100,0]#n=7
KruskalResult(statistic=0.8687350835322164, pvalue=0.351305729128759)
Ttest_indResult(statistic=0.9143731510776307, pvalue=0.37897292811768657)
```

2. *Include COL1A1 and PKHD cryptic 3'ss PSI 99 (N=14)*

| | PSI |
|---|---|
| -3C | *HEXA:* 100, *COL1A2:* 41, *CLCN2:* 81, *LAMA2:* 100, *NEU1:* 100, *COL6A2:* 100 |
| -3Csub | *GH1:* 1, *FECH:* 17, *EYA1:* 66, *LPL:* 100, *CAPN3*ex10: 91, *COL1A1:* 100, *CAPN3*ex17: 0, *PKHD1:* 99 |

```
pos3C=[100,41,81,100,100,100]#n=6
pos3Csub=[1,17,66,100,91,100,0,99]#n=8
KruskalResult(statistic=2.184722222222221, pvalue=0.13938620946554242)
Ttest_indResult(statistic=1.4712477438172564, pvalue=0.16933922490210684)
```

3. *Include PKHD normal 3'ss PSI 0 and exclude COL1A1 (N=13)*

| | PSI |
|---|---|
| -3C | *HEXA:* 100, *COL1A2:* 41, *CLCN2:* 81, *LAMA2:* 100, *NEU1:* 100, *COL6A2:* 100, *PKHD1:* 0 |
| -3Csub | *GH1:* 1, *FECH:* 17, *EYA1:* 66, *LPL:* 100, *CAPN3*ex10: 91, *CAPN3*ex17: 0, |

```
pos3C=[100,41,81,100,100,100,0]#n=7
pos3Csub=[1,17,66,100,91,0]#n=6
KruskalResult(statistic=1.5647646813827558, pvalue=0.21096894075885705)
Ttest_indResult(statistic=1.2080619908622303, pvalue=0.2546947882170792)
```

4. *Include PKHD cryptic 3'ss PSI 99 and exclude COL1A1 (N=13)*

| | PSI |
|---|---|
| -3C | *HEXA:* 100, *COL1A2:* 41, *CLCN2:* 81, *LAMA2:* 100, *NEU1:* 100, *COL6A2:* 100 |
| -3Csub | *GH1:* 1, *FECH:* 17, *EYA1:* 66, *LPL:* 100, *CAPN3*ex10: 91, *CAPN3*ex17: 0, *PKHD1:* 99 |

```
pos3C=[100,41,81,100,100,100]#n=6
pos3Csub=[1,17,66,100,91,0,99]#n=7
KruskalResult(statistic=3.1096345514950166, pvalue=0.0778304219058807)
Ttest_indResult(statistic=1.6833419314017677, pvalue=0.12573231405797983)
```

*2.Include COL1A1 and PKHD cryptic 3'ss PSI 99 (N=14)*

```
## Calculations from lists (unequal sizes are acceptable)
pos3C=c(100,41,81,100,100,100)#n=6
pos3Csub=c(1,17,66,100,91,100,0,99)#n=8

> kruskal.test(list(pos3C,pos3Csub))#uses vectors without NaNs

	Kruskal-Wallis rank sum test

data:  list(pos3C, pos3Csub)
Kruskal-Wallis chi-squared = 2.1847, df = 1, p-value = 0.1394

> shapiro.test(pos3Csub)$p.value
[1] 0.02263669
> shapiro.test(pos3C)$p.value
[1] 0.002399249
> sd(pos3Csub)
[1] 45.73761
> sd(pos3C)
[1] 23.78235
> (sd(pos3Csub))^2
[1] 2091.929
> (sd(pos3C))^2
[1] 565.6

> t.test(pos3C,pos3Csub,var.equal = FALSE)#uses vectors without NaNs

	Welch Two Sample t-test

data:  pos3C and pos3Csub
t = 1.4712, df = 10.962, p-value = 0.1693
alternative hypothesis: true difference in means is not equal to 0
95 percent confidence interval:
 -13.78145  69.28145
sample estimates:
mean of x mean of y 
    87.00     59.25 

###Dataframe (requires NaNs added to smaller groups to make sizes equal)
pos3C=c(100,41,81,100,100,100,NaN,NaN)#n=6
pos3Csub=c(1,17,66,100,91,100,0,99)#n=8
U5_table=data.frame(cbind(pos3C,pos3Csub))
U5_table_stacked=stack(U5_table)
#kruskal.test and t.test produce the same result as with lists without NaNs
kruskal.test(values~ind,U5_table_stacked)#uses df with NaNs

> oneway.test(values~ind,U5_table_stacked, var.equal = FALSE)#uses df with NaNs

	One-way analysis of means (not assuming equal variances)

data:  values and ind
F = 2.1646, num df = 1.000, denom df = 10.962, p-value = 0.1693
```

*4.Include PKHD cryptic 3'ss PSI 99 and exclude COL1A1 (N=13)*

```
### Calculations from lists (unequal sizes are acceptable)
pos3C=c(100,41,81,100,100,100)#n=6
pos3Csub=c(1,17,66,100,91,0,99) #n=7

> kruskal.test(list(pos3C,pos3Csub))#uses vectors without NaNs

     Kruskal-Wallis rank sum test

data:  list(pos3C, pos3Csub)
Kruskal-Wallis chi-squared = 3.1096, df = 1, p-value = 0.07783

> shapiro.test(pos3Csub)$p.value
[1] 0.06937585
> shapiro.test(pos3C)$p.value
[1] 0.002399249
> sd(pos3Csub)
[1] 46.08997
> sd(pos3C)
[1] 23.78235
> (sd(pos3Csub))^2
[1] 2124.286
> (sd(pos3C))^2
[1] 565.6

> t.test(pos3C,pos3Csub,var.equal = FALSE)#uses vectors without NaNs

     Welch Two Sample t-test

data:  pos3C and pos3Csub
t = 1.6833, df = 9.237, p-value = 0.1257
alternative hypothesis: true difference in means is not equal to 0
95 percent confidence interval:
 -11.36769  78.51054
sample estimates:
mean of x mean of y
 87.00000  53.42857

###Dataframe (requires NaNs added to smaller groups to make sizes equal)#For
pos3C=c(100,41,81,100,100,100,NaN)#n=6
pos3Csub=c(1,17,66,100,91,0,99) #n=7
U5_table=data.frame(cbind(pos3C,pos3Csub))
U5_table_stacked=stack(U5_table)
#kruskal.test and t.test produce the same result as with lists without NaNs
kruskal.test(values~ind,U5_table_stacked)#uses df with NaNs
t.test(values~ind,U5_table_stacked, var.equal = FALSE)#uses df with NaNs

> oneway.test(values~ind,U5_table_stacked, var.equal = FALSE)#uses df with NaNs

     One-way analysis of means (not assuming equal variances)

data:  values and ind
F = 2.8336, num df = 1.000, denom df = 9.237, p-value = 0.1257
```

Boxplots for PSI dependent on +2C/+3G, substitute *A* or *T* and -3C

1. *Include COL1A1 and PKHD normal 3’ss PSI 0 (N=14)*

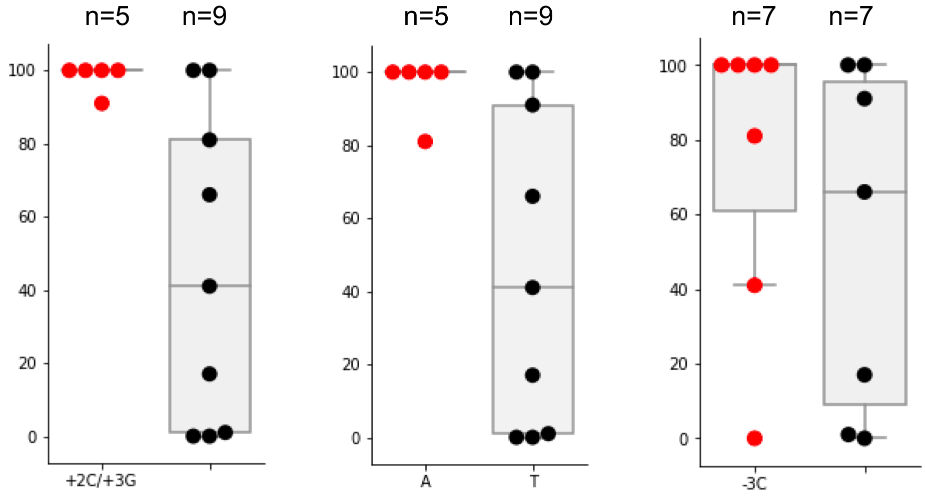


2. *Include COL1A1 and PKHD cryptic 3’ss PSI 99 (N=14)*

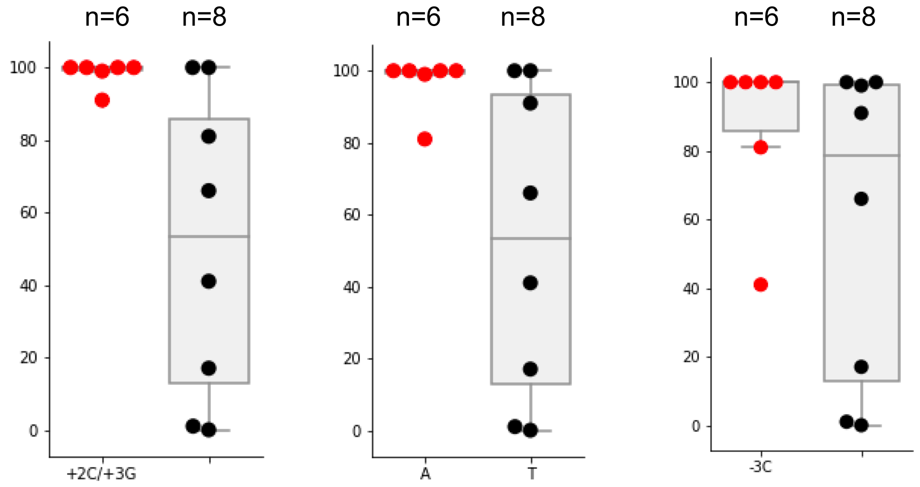


3. *Include PKHD normal 3’ss PSI 0 and exclude COL1A1 (N=13)*

4. *Include PKHD cryptic 3’ss PSI 99 and exclude COL1A1 (N=13)*

Spearman's correlation for PPS length

| Gene | Ex | PSI | PPT Stretch (PPS) |
|---|---|---|---|
| *GH1* | 3 | 1 | 10 |
| *FECH* | 9 | 17 | 6 |
| *EYA1* | 10 | 66 | 5 |
| *LPL* | 5 | 100 | 14 |
| *HEXA* | 13 | 100 | 13 |
| *PKHD1* | 25 | 99<br>**0** • | 6<br>3 • |
| *COL1A2* | 37 | 41 | 5 |
| *CLCN2* | 19 | 81 | 4 |
| *LAMA2* | 24 | 100 | 9 |
| *CAPN3* | 10 | 91 | 10 |
| *CAPN3* | 17 | 0 | 9 |
| *NEU1* | 2 | 100 | 14 |
| *COL6A2* | 8 | 100 | 15 |
| *COL1A1* | 23 | 100 • | 16 • |

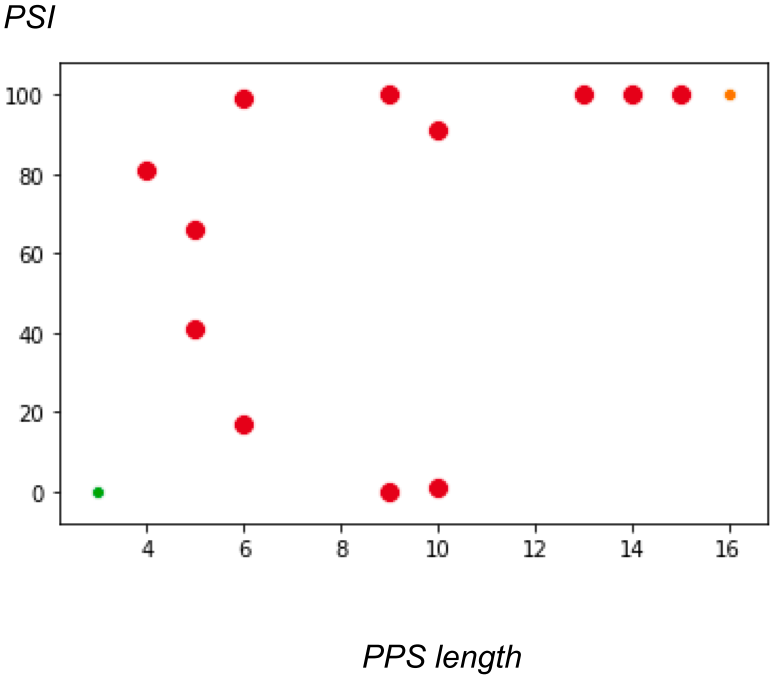


1. *Include COL1A1 and PKHD normal 3'ss PSI 0 (N=14)*

```
X=c(10,6, 5, 14, 13, 3,5, 4, 9,  10,9,14, 15, 16)#PPS length
Y=c(1, 17,66,100,100,0,41,81,100,91,0,100,100,100)#PSI
SpearmanrResult(correlation=0.7108258237527484, pvalue=0.004375165814040274)
```

2. *Include COL1A1 and PKHD cryptic 3'ss PSI 99 (N=14)*

```
X=c(10,6, 5, 14, 13, 6, 5, 4, 9,  10,9,14, 15, 16)#PPS length
Y=c(1, 17,66,100,100,99,41,81,100,91,0,100,100,100)#PSI
SpearmanrResult(correlation=0.6360104238605363, pvalue=0.014485699211541149)
```

3. *Include PKHD normal 3'ss PSI 0 and exclude COL1A1 (N=13)*

```
X=c(10,6, 5, 14, 13, 3,5, 4, 9,  10,9,14, 15) #PPS length
Y=c(1, 17,66,100,100,0,41,81,100,91,0,100,100) #PSI
SpearmanrResult(correlation=0.6701817559482136, pvalue=0.012192348615293494)
```

4. *Include PKHD cryptic 3'ss PSI 99 and exclude COL1A1 (N=13)*

```
X=c(10,6, 5, 14, 13, 6, 5, 4, 9,  10,9,14, 15) #PPS length
Y=c(1, 17,66,100,100,99,41,81,100,91,0,100,100) #PSI
SpearmanrResult(correlation=0.5776564039581658, pvalue=0.03868651748677648)
```

Spearman's correlation for branchpoint matches

| Gene | Ex | PSI | BP matches |
|---|---|---|---|
| *GH1* | 3 | 1 | 1 |
| *FECH* | 9 | 17 | 4 |
| *EYA1* | 10 | 66 | 3 |
| *LPL* | 5 | 100 | 4 |
| *HEXA* | 13 | 100 | 2 |
| *PKHD1* | 25 | 99<br>**0** | 3<br>3 |
| *COL1A2* | 37 | 41 | 3 |
| *CLCN2* | 19 | 81 | 4 |
| *LAMA2* | 24 | 100 | 4 |
| *CAPN3* | 10 | 91 | 2 |
| *CAPN3* | 17 | 0 | 4 |
| *NEU1* | 2 | 100 | 3 |
| *COL6A2* | 8 | 100 | 4 |
| *COL1A1* | 23 | 100 | 0 |

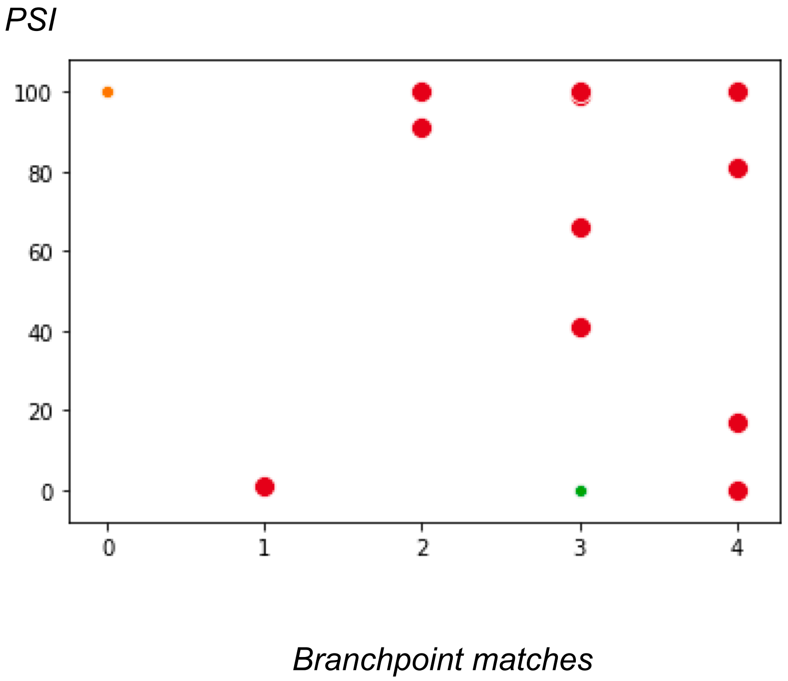


1. *Include COL1A1 and PKHD normal 3'ss PSI 0 (N=14)*
```
X=c(1,4, 3, 4,  2,  3,3, 4, 4,  2, 4,3,  4,  0)
Y=c(1,17,66,100,100,0,41,81,100,91,0,100,100,100)
```
**SpearmanrResult(correlation=-0.025**36416873014165, **pvalue=0.931**4119671811683)

2. *Include COL1A1 and PKHD cryptic 3'ss PSI 99 (N=14)*
```
X=c(1,4, 3, 4,  2,  3, 3, 4, 4,  2, 4,3,  4,  0)
Y=c(1,17,66,100,100,99,41,81,100,91,0,100,100,100)
```
**SpearmanrResult(correlation=-0.049**46153176891343, **pvalue=0.866**6504276289523)

3. *Include PKHD normal 3'ss PSI 0 and exclude COL1A1 (N=13)*
```
X=c(1,4, 3, 4,  2,  3,3, 4, 4,  2, 4,3,  4)
Y=c(1,17,66,100,100,0,41,81,100,91,0,100,100)
```
**SpearmanrResult(correlation=0.124**14328599550853, **pvalue=0.686**1588126520046)

4. *Include PKHD cryptic 3'ss PSI 99 and exclude COL1A1 (N=13)*
```
X=c(1,4, 3, 4,  2,  3, 3, 4, 4,  2, 4,3,  4)
Y=c(1,17,66,100,100,99,41,81,100,91,0,100,100)
```
**SpearmanrResult(correlation=0.093**72790559206708, **pvalue=0.760**7010691102144)

Scatterplots for PSI dependent on PPS length and branchpoint matches

1. *Include COL1A1 and PKHD normal 3'ss PSI 0 (N=14)*

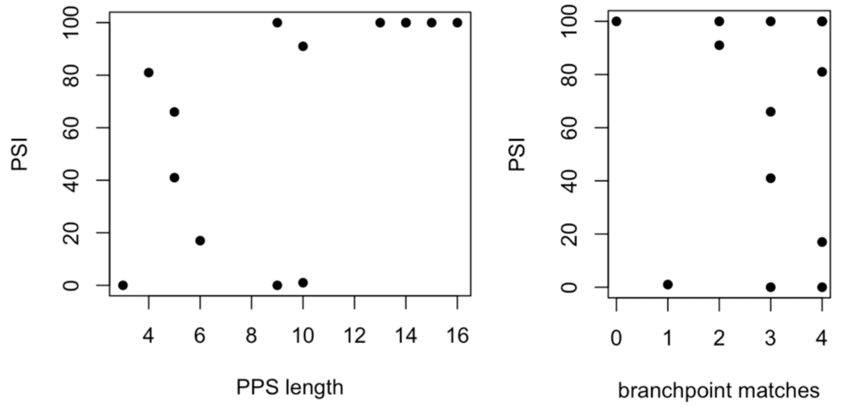


2. *Include COL1A1 and PKHD cryptic 3'ss PSI 99 (N=14)*

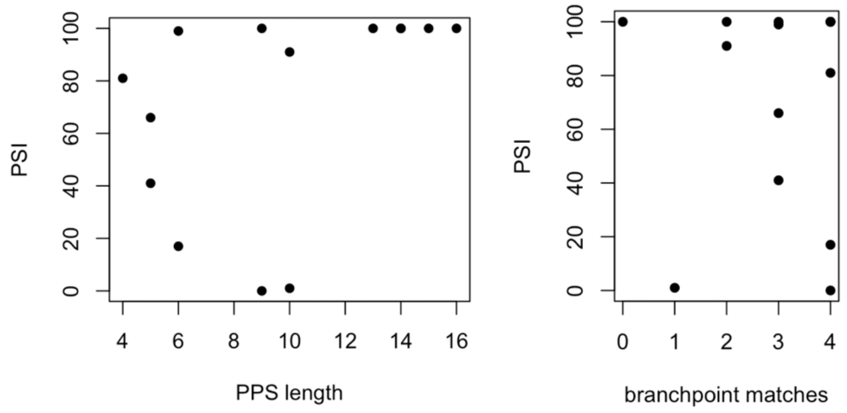


3. *Include PKHD normal 3'ss PSI 0 and exclude COL1A1 (N=13)*

4. *Include PKHD cryptic 3'ss PSI 99 and exclude COL1A1 (N=13)*

5. *All-in-one*

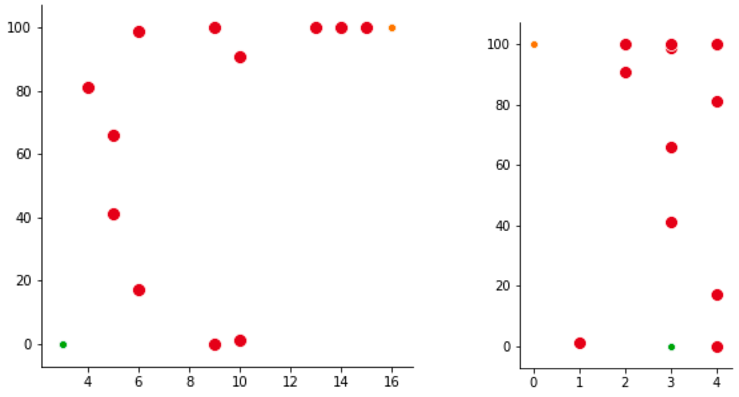

**Section S5: Future work: Molecular cell biology testing of the U5 hypothesis**

Below, we include suitable human mutations for the proof-of-principle laboratory testing:

1. A **Becker muscular dystrophy** patient with c9563+5G>C mutation in intron 65 (Juan-Mateu et al., 2013) has a dystrophin deficiency due to the activation of two alternative cryptic 5' splice sites. Wild-type dystrophin exon 65 unusually ends with a C (c9563), while the cryptic sites both have Gs for the exon-end. We can attempt to restore normal splicing in a minigene construct in cultured cells with a second compensatory mutation of exon 65 -1C>G (c9563C>G) to match the wild type U5 39C. Reciprocal experiment, a swap of G and C between the interacting RNAs, can involve modified U5 39C>G to match the wt exon 65 -1C (**Figure S5)**. U5 should be checked out independently and in combination with the matching U1 9C>G.

2. A rare A>G substitution in the **coagulation factor V** gene (*F5-Texas*, Vincent et al., 2013), defines an end of an exon and activates splicing of an alternative intron, causing a bleeding phenotype. Interestingly, a wt *F5-short* was later identified in normal individuals as a rare splice isoform that regulates anticoagulation. *F5-Texas* mutant guanine at the end of the exon preceding the alternative intron activates splicing by the factor of $10^2$ compared to the wt A in this position. Again, an experiment with a minigene or full length F5 cDNA construct can serve to swap the G and C between the interacting RNAs: F5 alternative intron preceded by -1A>C mutation and modified U5 39C>G. Again, modified U1 9C>G can be considered in parallel (**Figure S6**). The future clinical application involves regulation of anticoagulation for the prevention of common venous thrombosis by targeting F5 alternative intron with a full set of complementary snRNAs.

Follow up of the previous experiments that have already determined that a change in the exon sequence at the splice junction can suppress a different splicing mutation:

3. Mutation study of the ***IKBKAP*** gene (Introduction; Carmel et al., 2004) indicate that substitution of a rare exon-end A for G fully restores normal splicing, blocked by +6T>C mutation. Some restoration of normal splicing is achieved using U1 9C>U modification to match the wild-type end of exon A. In case of U5 snRNA a change for one more U in the uracil-rich Loop1 might be not very useful, so, we suggest starting with U5 39C>G modification combined with the exon-end A for C substitution to introduce the swap G=C pair. U1 9C>G can be tried in parallel.

4. The study of the mutant **ATR** gene (Introduction; Scalet et al., 2017) can be adapted to change the end of exon 9 A>C co-transfected with modified U5 C39>G gene to check if the swap G=C pair will also correct spicing independently or in combination with the matching U1 9C>G snRNA. At the wt 3'ss exon 9 conveniently starts with a C, which will match U5 C38>G modification.

5. The compensatory mutation study of the ***FAH*** c1062+5G>A allele (Discussion above; Scalet et al., 2018) can be followed by testing the effect of U5 37U>C modification to create U5 $C_{37}$=$G_{-2}$ pair with wt exon 12 in combination with the modified U1 version from the previous study.

Follow up of an exon-end G>A mutation:

6. Finally, in a very recent study (Breuel et al., 2019) from Prof Neidhardt laboratory, who pioneered combining both modified U1 and U6 for splicing correction (Schmid et al., 2013), Breuel and colleagues use adapted U1 snRNA together with antisense oligos to correct the G>A substitution at the end of **BBS** exon 5. In order to verify if a combination of modified U1 and U5 will be effective to correct this mutation, we suggest first changing it in the minigene for G>C and co-transfecting with the matching U5 C39>G and U1 9C>G genes (alternatively and combined).

Testing U2 snRNA interaction with the end of the intron (the proposed U2 $G_{31}$=$C_{-3}$ pair) is more straightforward and any human intron can do, however the **Fas/CD95** intron 5 (Corrionero et al, 2011) is an excellent study to follow. The inclusion of Fas receptor exon 6 is an apoptotic switch, which is disabled by -3C>G mutation in intron 5 leading to accumulation of T-cells in autoimmune limphoproliferative syndrome (ALPS). We can attempt to rescue normal splicing of intron 5 in a minigene construct using modified U2 31G>C (**Figure S7**).

Therapeutic application for every case outlined above will involve the assessment of the efficacy of a full set of complementary snRNAs (U1, U6, U5 and U2) specific to the target intron and splice junction and with a swapped U2/U6 helix II, which will decrease the intermixing with the endogenous snRNAs.

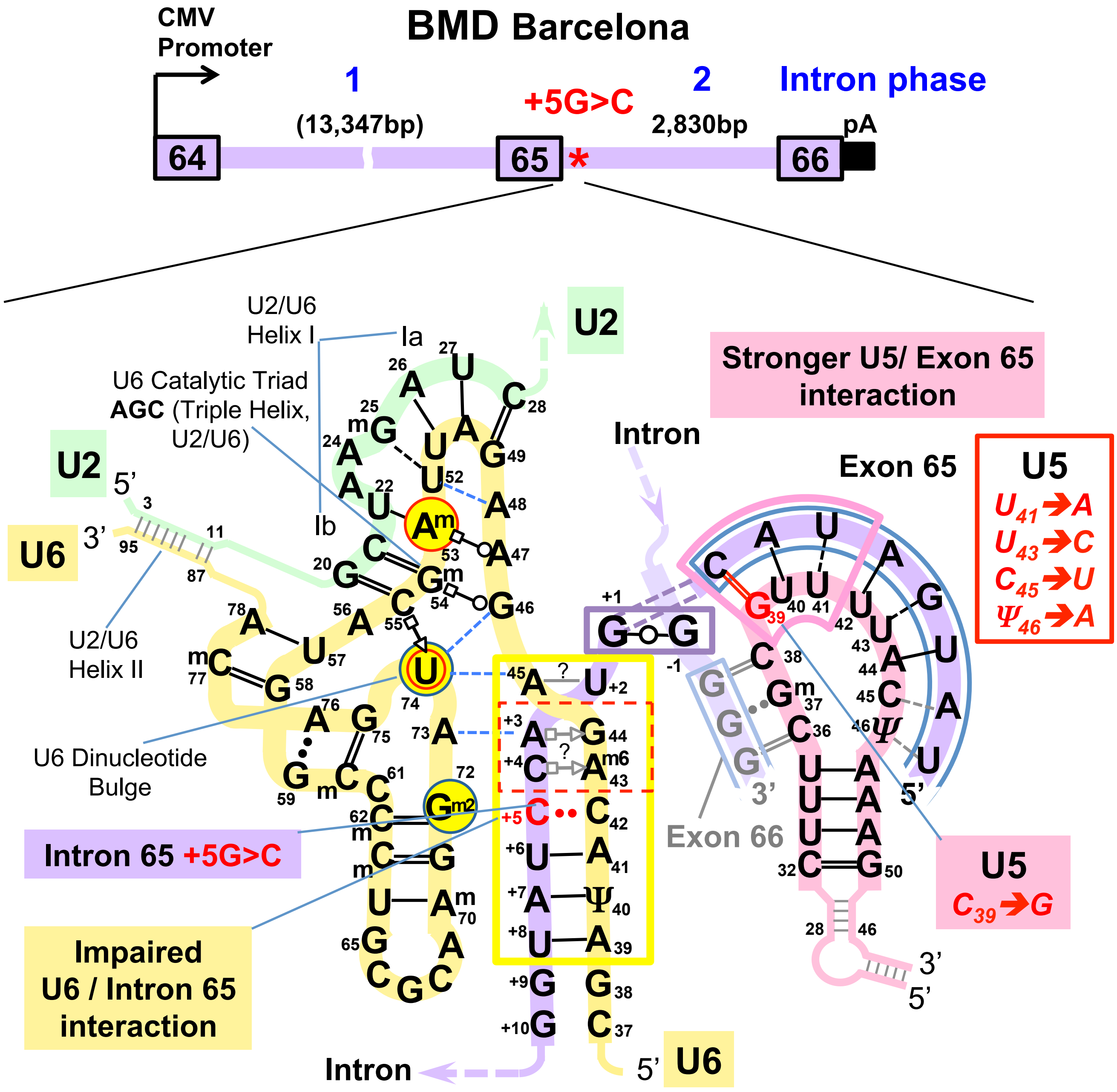


**Figure S5|** Study design for the correction of the dystrophin gene splicing mutation c9563+5G>C in intron 65 from a Becker muscular dystrophy patient reported by Juan-Mateu et al., 2013. The U5 snRNA modifications, matching wt exon 65 are shown here, as we are specifically interested if U5 can compensate for the loss of the most conserved Watson-Crick pair at the U6 binding site. For therapeutic purpose all spliceosomal snRNAs can be modified to increase complementarity to the dystrophin intron 65. U1 and U6 are obvious candidates for modifications matching the mutant +5C.

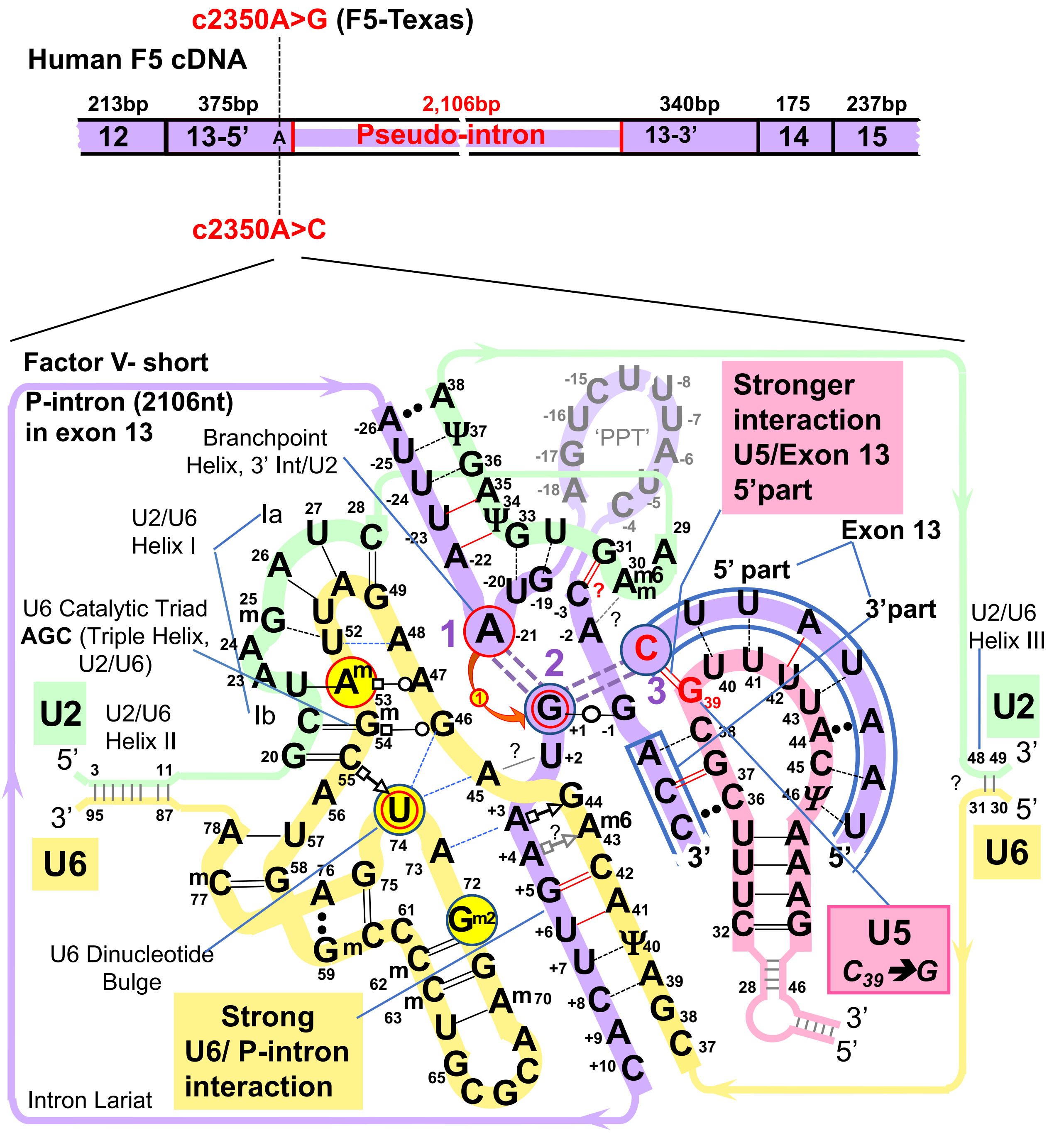


**Figure S6|** Study design targeting the alternative intron (pseudo-intron) splicing of coagulation F5. Normally, 1% of the circulating F5 protein is the short isoform, lacking the part that corresponds to the alternative intron in exon 13. F5-Texas mutation (Vincent et al., 2013) leads to exclusive production of the F5-short isoform and bleeding disorder. To explore U5 snRNA base pairing with the conserved exon-end guanine we can try to introduce a co-variant pair and swap the G and C between the interacting RNAs: F5 c2350A>C mutation and modified U5 39C>G. Modified U1 9C>G can be considered in parallel. U1 9C>U and a full set of snRNAs complementary to the wt alternative intron might help to reduce coagulation in patients with thrombosis.

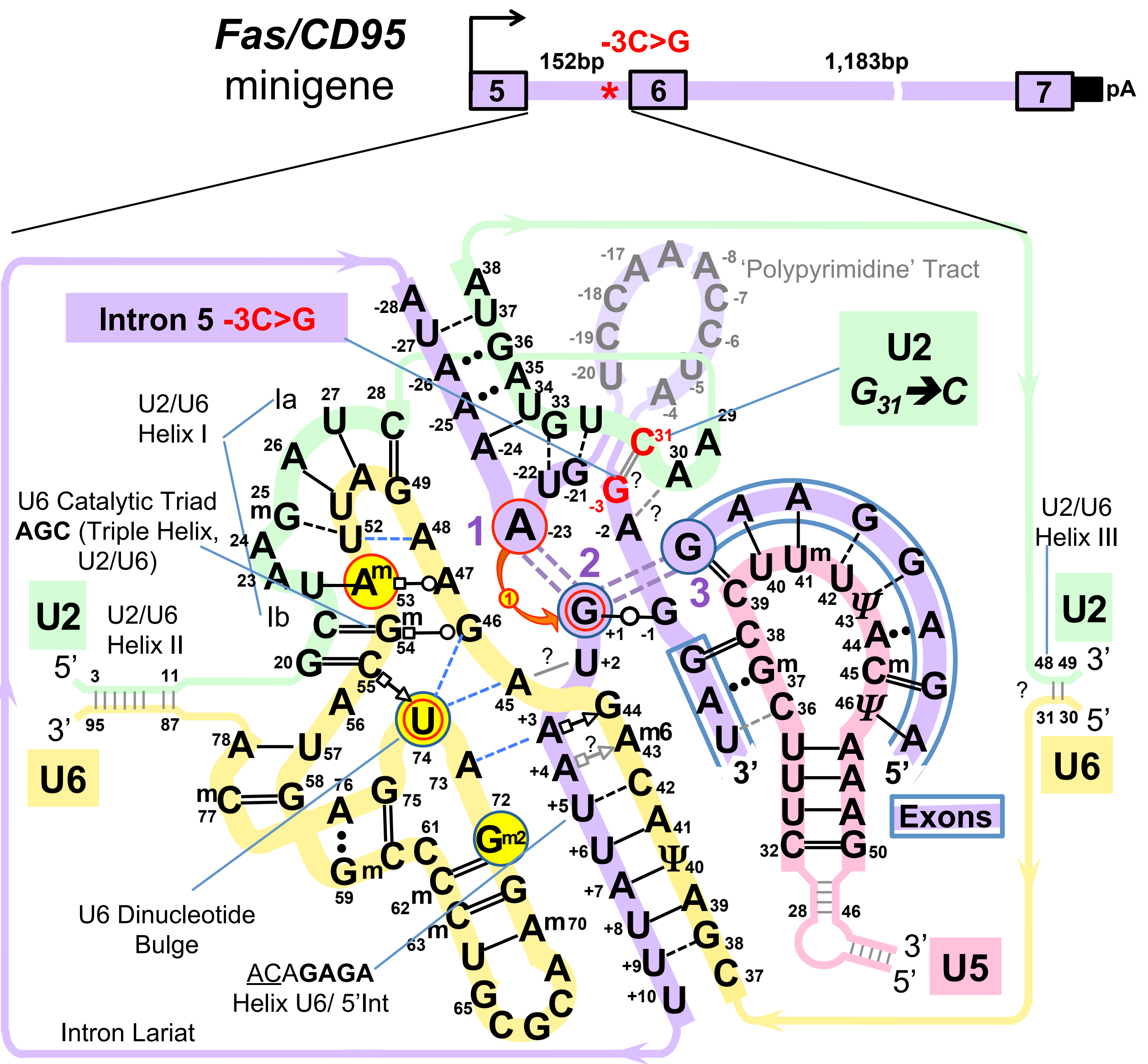


**Figure S7|** Testing U2 snRNA interaction with the end of the intron (the proposed U2 $G_{31}$=$C_{-3}$ pair) following the study of Corrionero et al., 2011 on Fas/CD95 intron 5. The inclusion of Fas receptor exon 6 is an apoptotic switch, which is disabled by -3C>G mutation in intron 5 leading to accumulation of T-cells in autoimmune limphoproliferative syndrome (ALPS). We can attempt to rescue normal splicing of intron 5 in a minigene construct using modified U2 31G>C.

#### Section S4: Results and Methods (continued)

*Histograms of bootstrap difference for U5 bp geometry at exon junctions of +5Gsub and +5G introns*

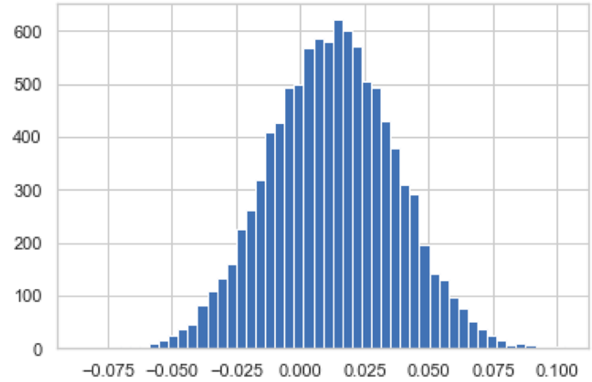


BD_5pr8_WC
P_H0_5pr8_WC 0.31

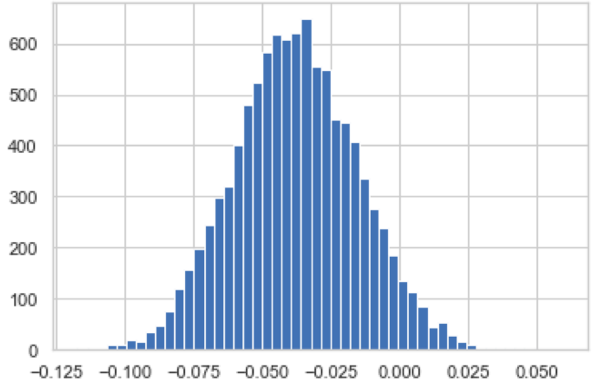


BD_5pr7_WC
P_H0_5pr7_WC 0.0486

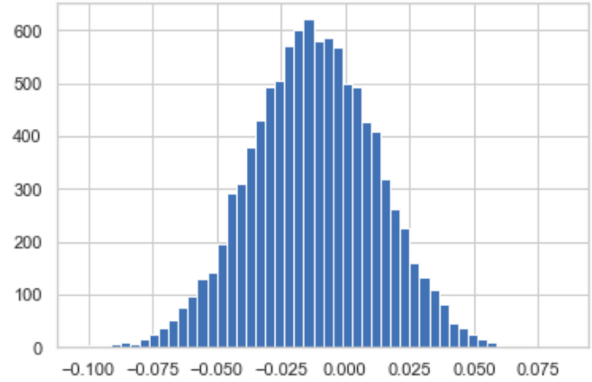


BD_5pr8_iso
P_H0_5pr8_iso 0.31

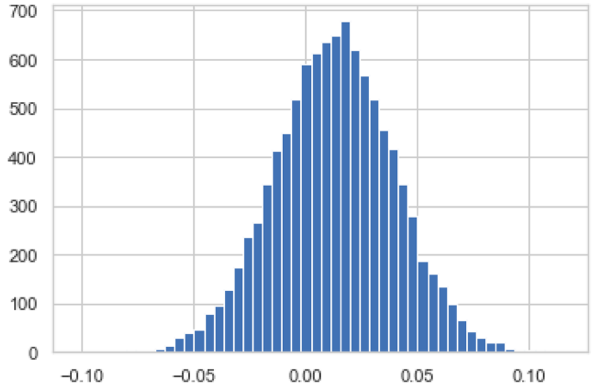


BD_5pr7_iso
P_H0_5pr7_iso 0.3104

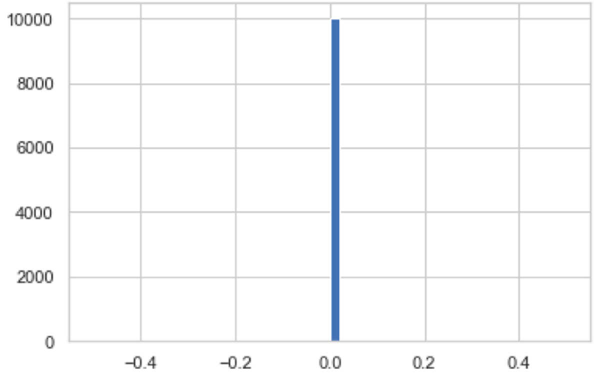


BD_5pr8_dif

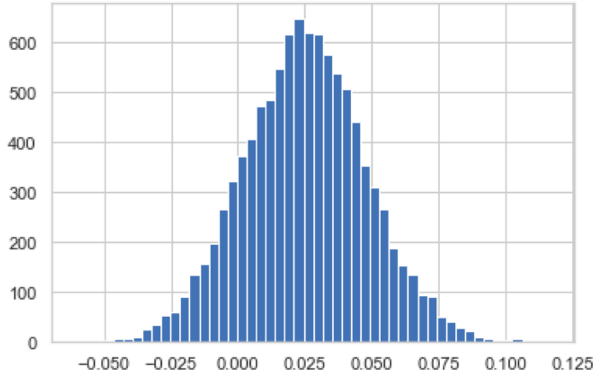


BD_5pr7_dif
P_H0_5pr7_dif 0.1414

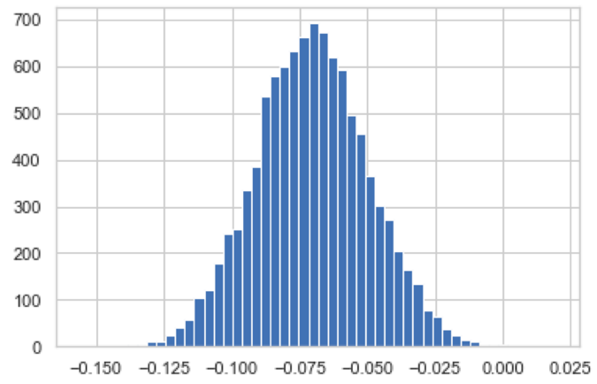


BD_5pr6_WC
P_H0_5pr6_WC 0.0007

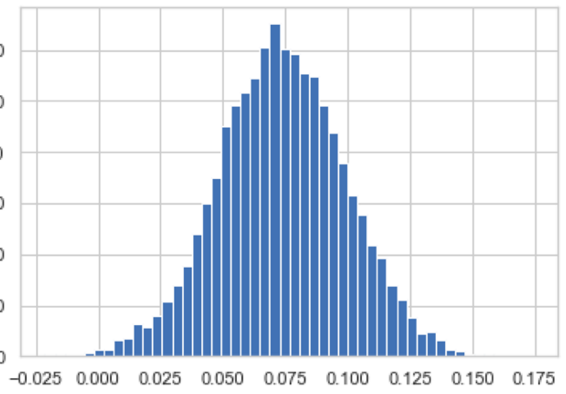


BD_5pr5_WC
P_H0_5pr5_WC 0.0025

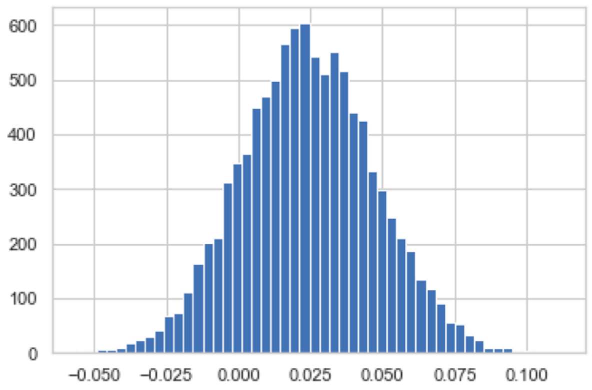


BD_5pr6_iso
P_H0_5pr6_iso 0.151

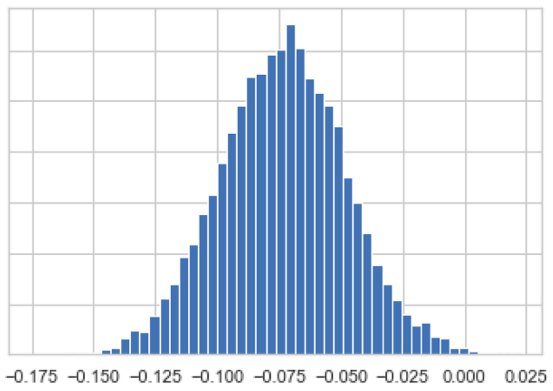


BD_5pr5_iso
P_H0_5pr5_iso 0.0025

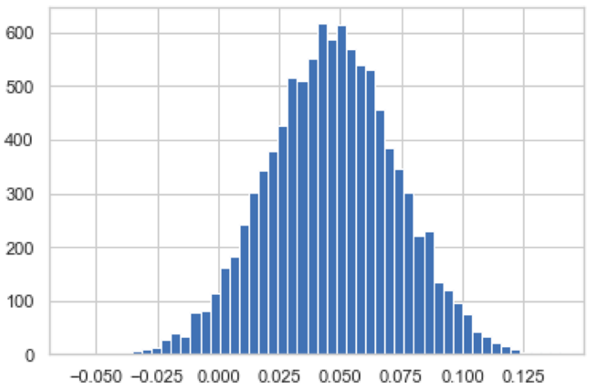


BD_5pr6_dif
P_H0_5pr6_dif 0.0394

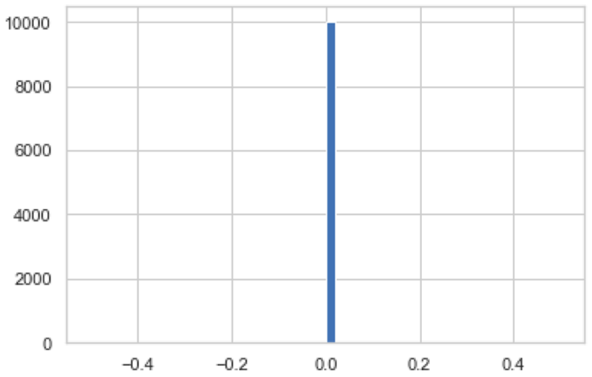


BD_5pr5_dif

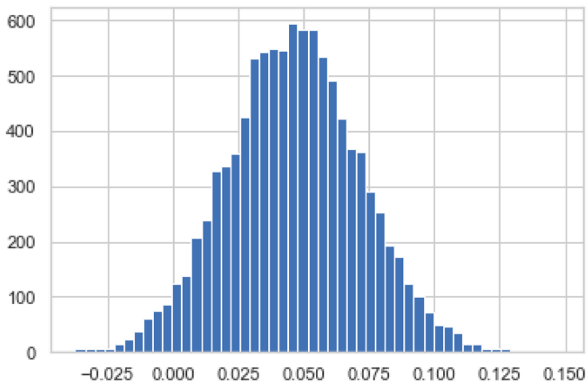


BD_5pr4_WC
P_H0_5pr4_WC 0.0336

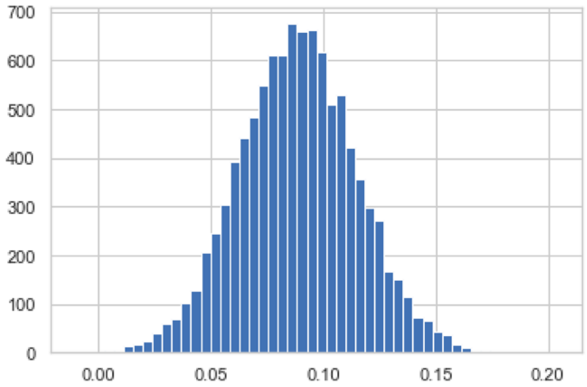


BD_5pr3_WC
P_H0_5pr3_WC 0.0003

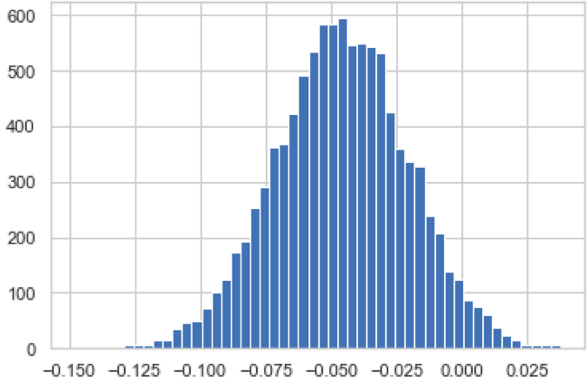


BD_5pr4_iso
P_H0_5pr4_iso 0.0336

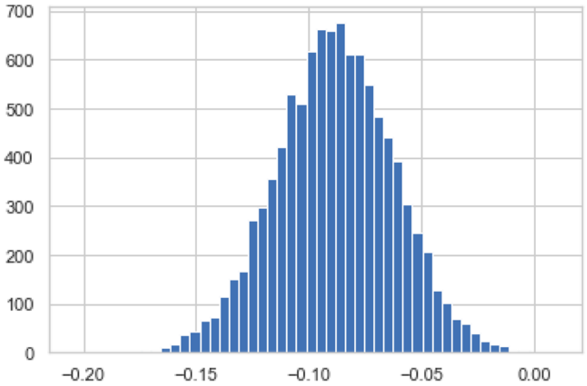


BD_5pr3_iso
P_H0_5pr3_iso 0.0003

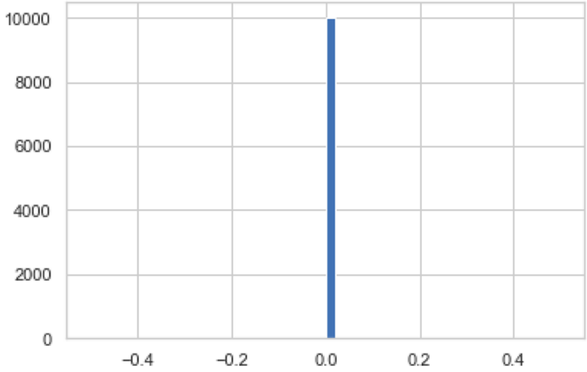


BD_5pr4_dif

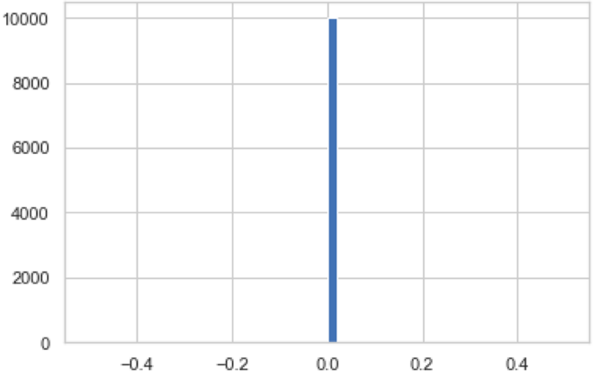


BD_5pr3_dif

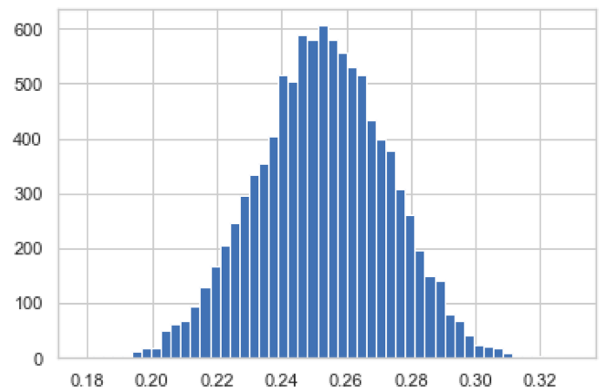

BD_5pr2_WC
P_H0_5pr2_WC 0.0

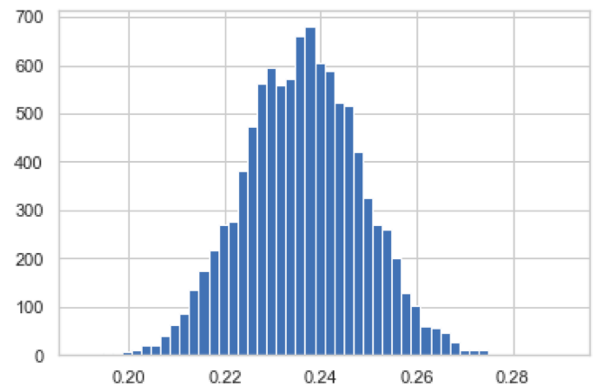

BD_5pr1_WC
P_H0_5pr1_WC 0.0

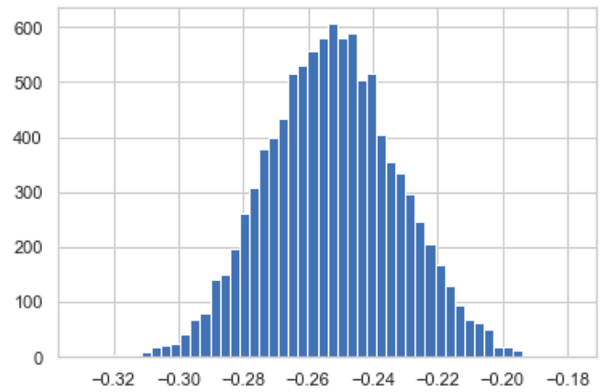

BD_5pr2_iso
P_H0_5pr2_iso 0.0

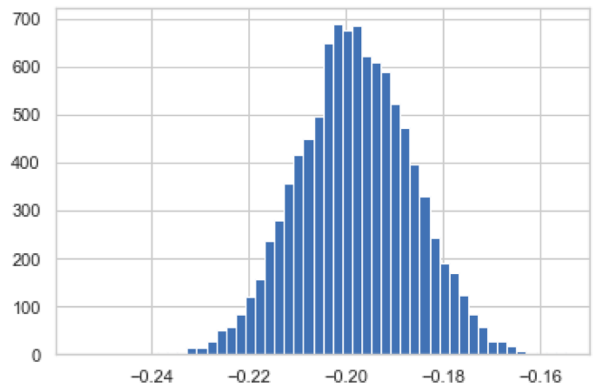

BD_5pr1_iso
P_H0_5pr1_iso 0.0

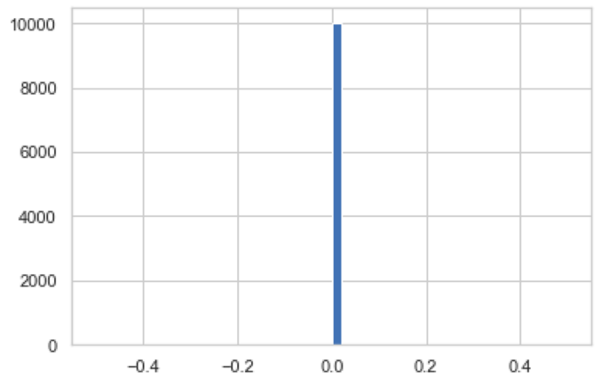

BD_5pr2_dif

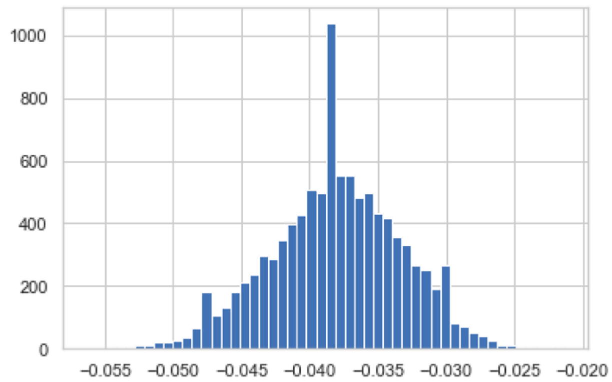

BD_5pr1_dif
P_H0_5pr1_dif 0.0

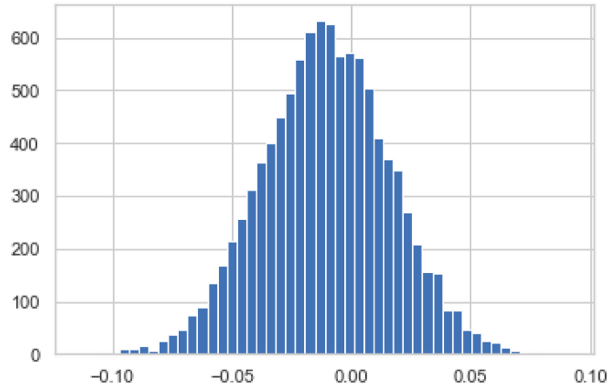


BD_3pr1_WC
P_H0_3pr1_WC 0.3473

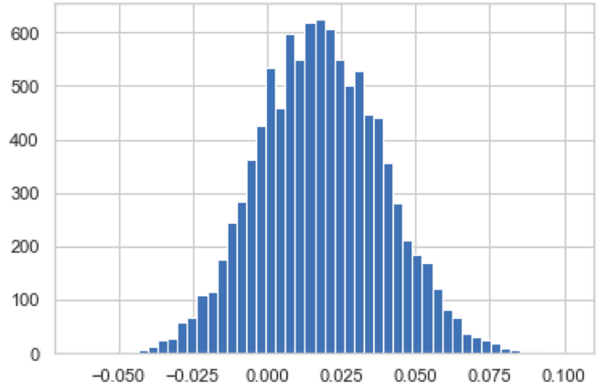


BD_3pr2_WC
P_H0_3pr2_WC 0.2026

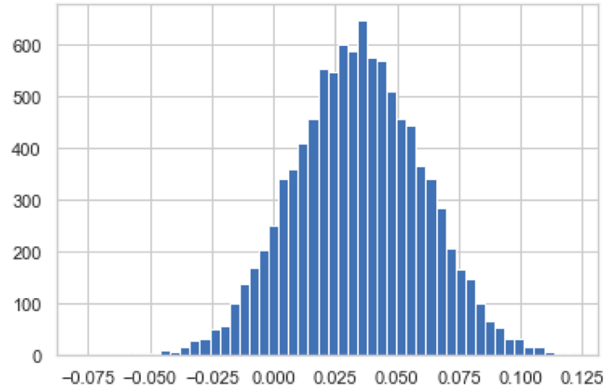


BD_3pr1_iso
P_H0_3pr1_iso 0.0951

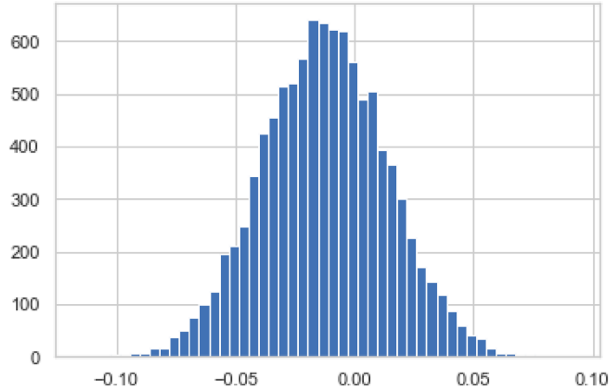


BD_3pr2_iso
P_H0_3pr2_iso 0.3149

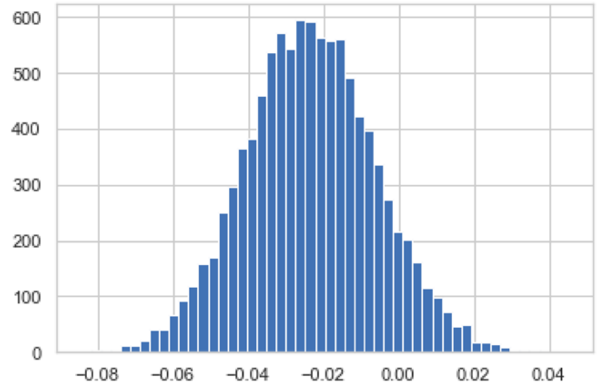


BD_3pr1_dif
P_H0_3pr1_dif 0.0886

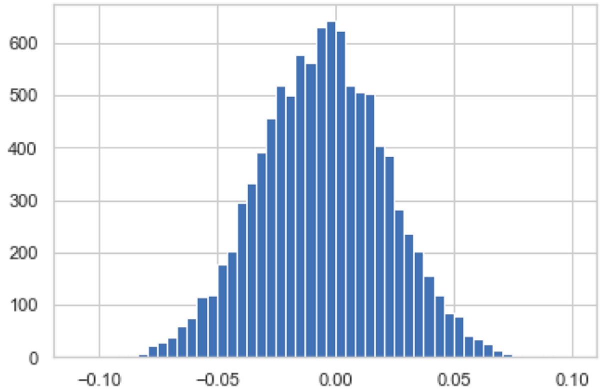


BD_3pr2_dif
P_H0_3pr2_dif 0.4189

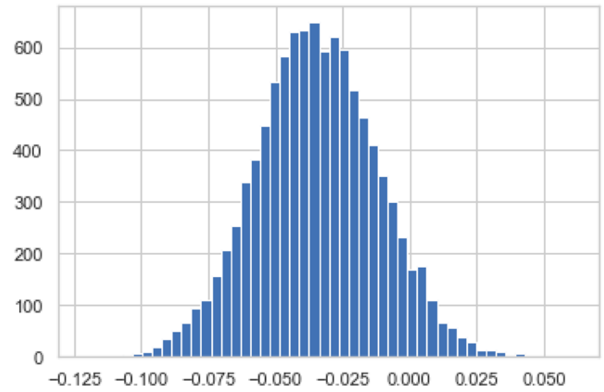


BD_3pr3_WC
P_H0_3pr3_WC 0.0656

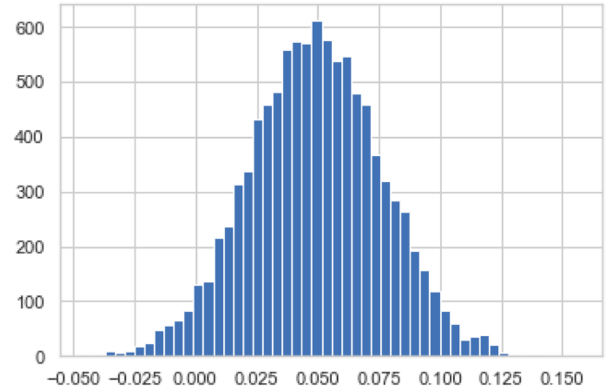


BD_3pr3_iso
P_H0_3pr3_iso 0.0357

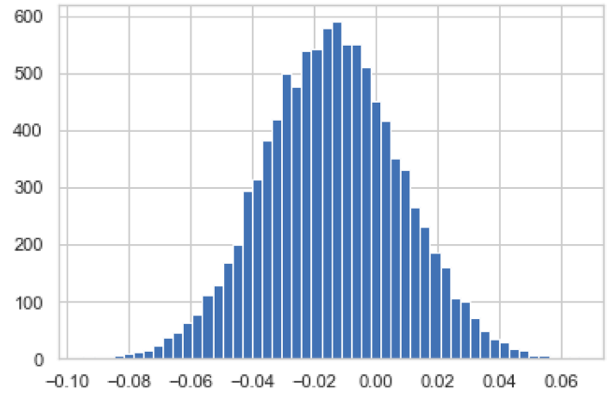


BD_3pr3_dif
P_H0_3pr3_dif 0.2606

**Figure S8|** Histograms of bootstrap difference for U5 bp types at each position of the exon junctions between +5Gsub and +5G datasets (Violinplots of the same **Figure 6A-C**)
BD = Distributions of **B**ootstrap **D**ifferences of the frequencies for U5 base pair types
Base pair types: **W**atson-**C**rick, **WC**; **iso**steric, **iso**; and non-isosteric (**dif**ferent), **dif**
Splice junction positions key: **5pr1** is 5'exon position -1; **3pr1** is 3'exon position +1

*Histograms of bootstrap difference for U6 bp geometry in introns preceded by -1Gsub and -1G exons*

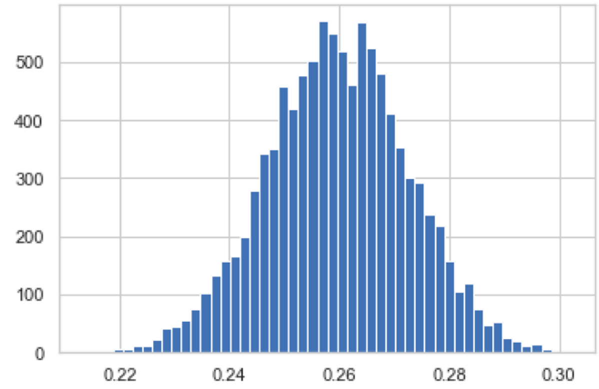


BD_in5_WC
P_H0_in5_WC 0.0

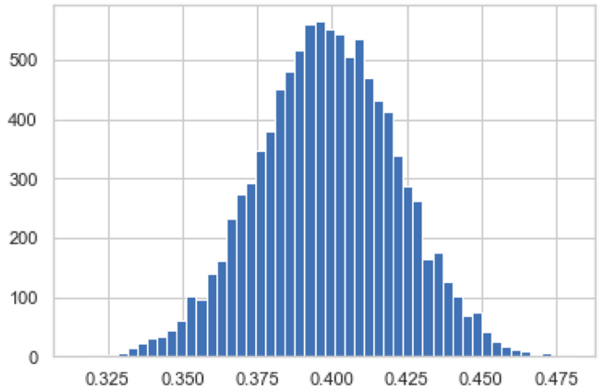


BD_in6_WC
P_H0_in6_WC 0.0

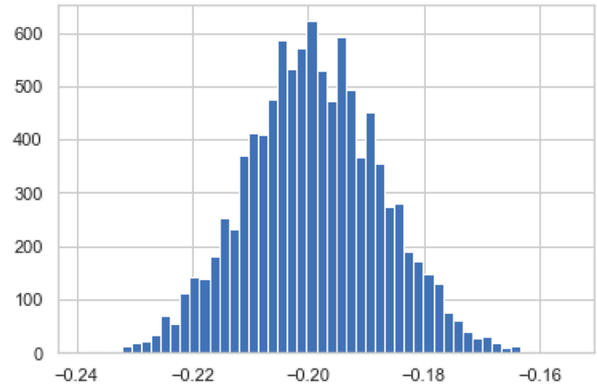


BD_in5_iso
P_H0_in5_iso 0.0

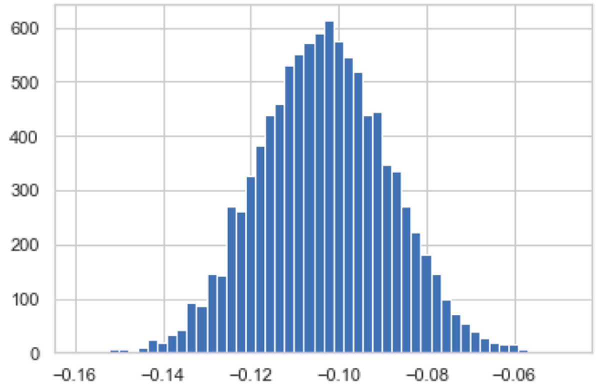


BD_in6_iso
P_H0_in6_iso 0.0

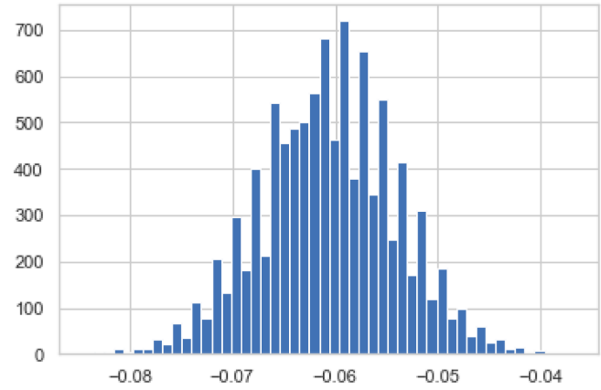


BD_in5_dif
P_H0_in5_dif 0.0

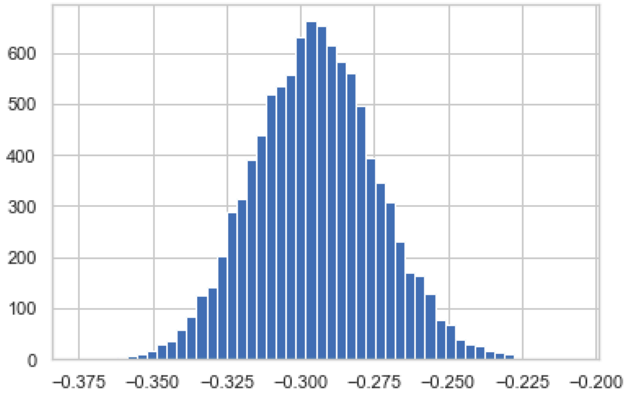


BD_in6_dif
P_H0_in6_dif 0.0

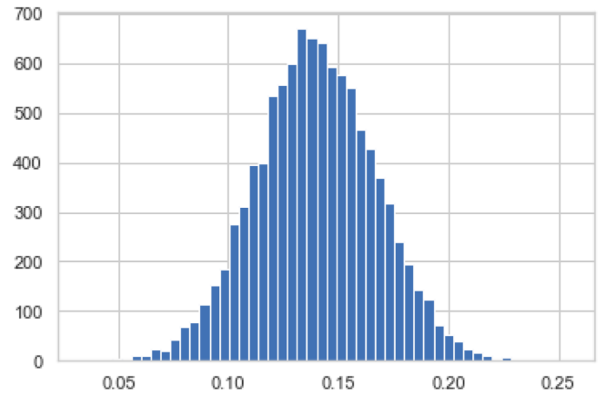


BD_in7_WC
P_H0_in7_WC 0.0

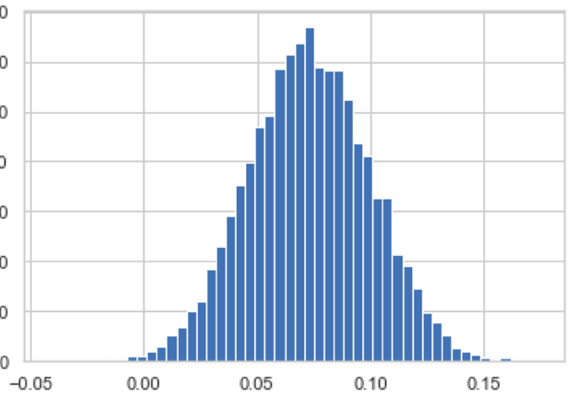


BD_in8_WC
P_H0_in8_WC 0.0029

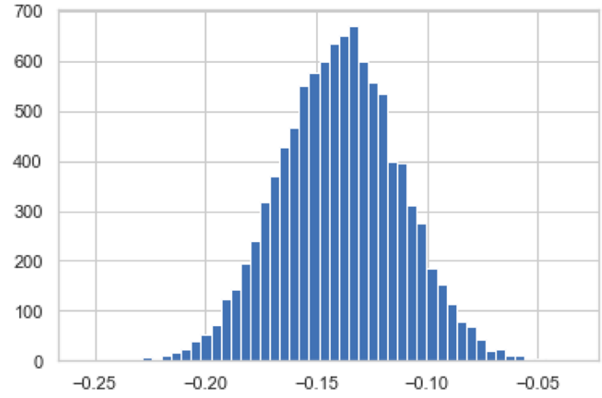


BD_in7_iso
P_H0_in7_iso 0.0

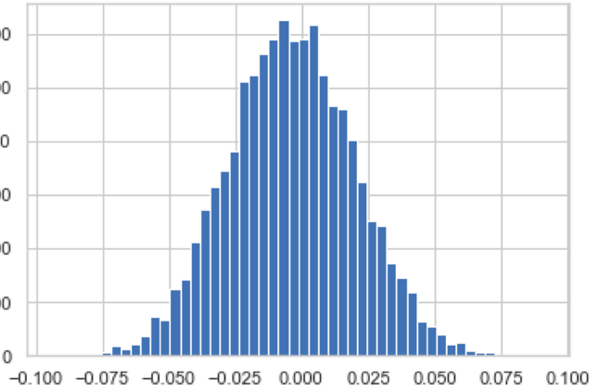


BD_in8_iso
P_H0_in8_iso 0.434

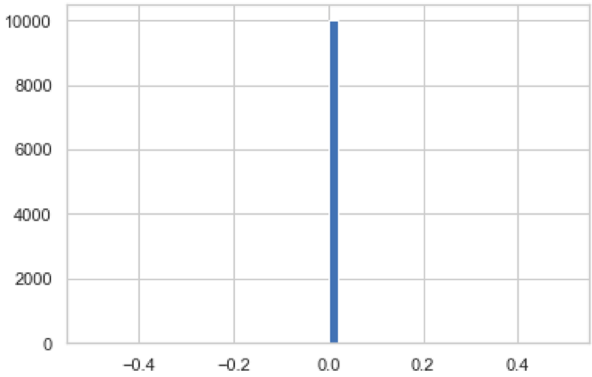


BD_in7_dif

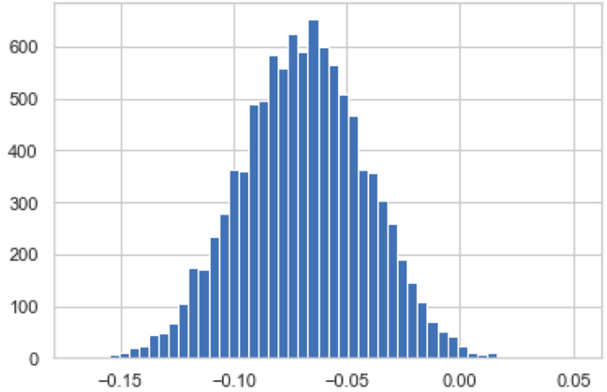


BD_in8_dif
P_H0_in8_dif 0.0051

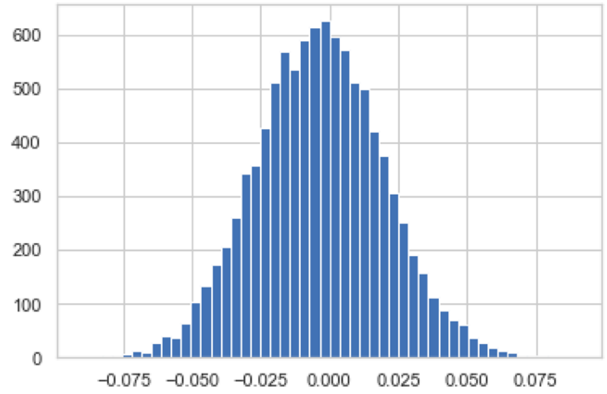


BD_in9_WC
P_H0_in9_WC 0.4314

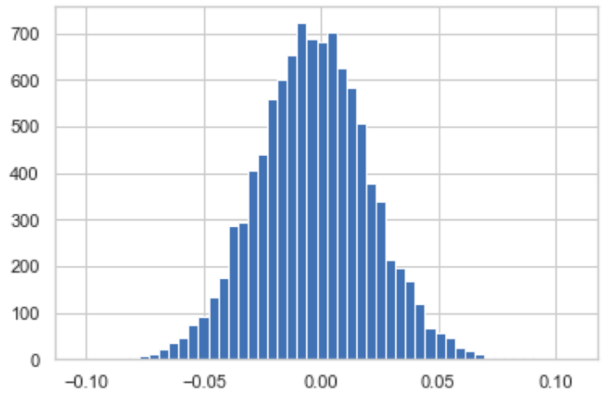


BD_in10_WC
P_H0_in10_WC 0.4462

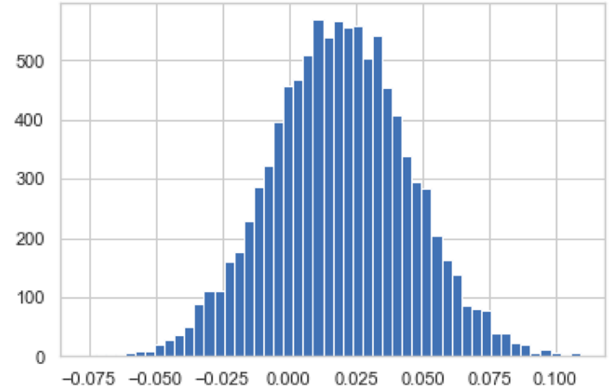


BD_in9_iso
P_H0_in9_iso 0.2326

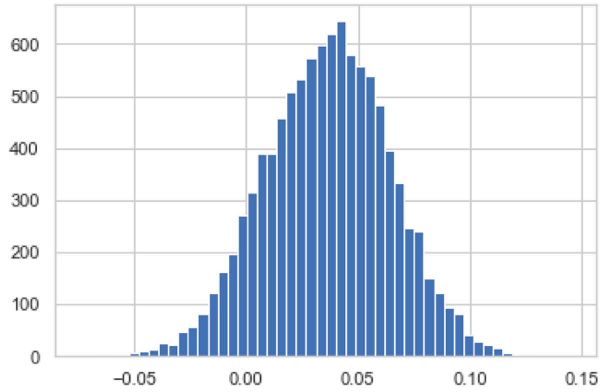


BD_in10_iso
P_H0_in10_iso 0.0993

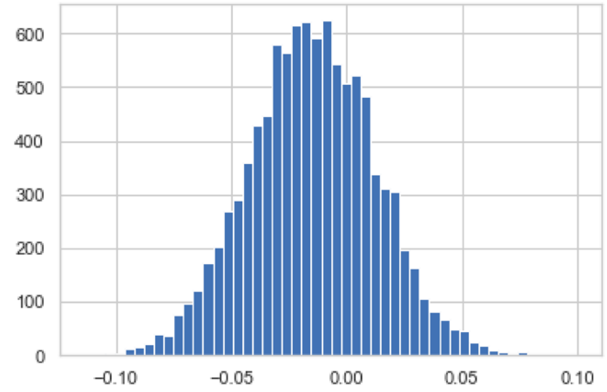


BD_in9_dif
P_H0_in9_dif 0.2933

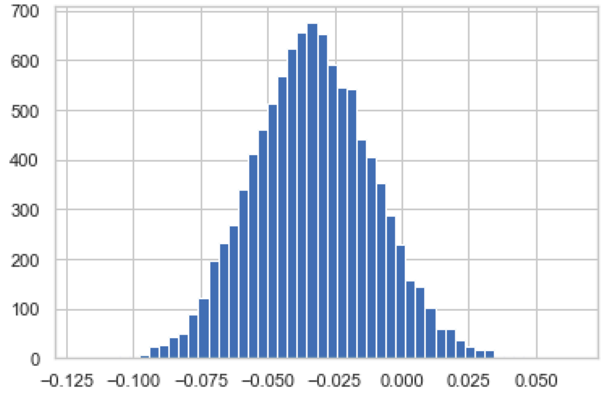


BD_in10_dif
P_H0_in10_dif 0.0717

**Figure S9|** Histograms of bootstrap difference for U6 bp types at the start of intron position +5 to +10 between -1Gsub and -1G datasets (-1G=exon-end G; Violinplots of the same **Figure 6D-F**)
Base pair types: **Watson-Crick**, **WC**; **iso**steric, **iso**; and non-isosteric (**dif**ferent), **dif**

*Histograms of bootstrap difference for U5 bp geometry at exon junctions of -3Csub and -3C introns*

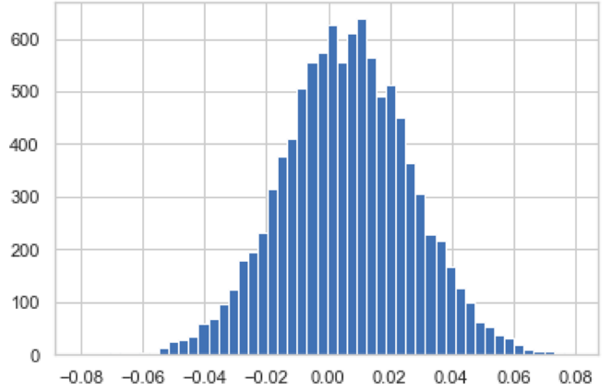


BD_5pr8_WC
P_H0_5pr8_WC 0.3906

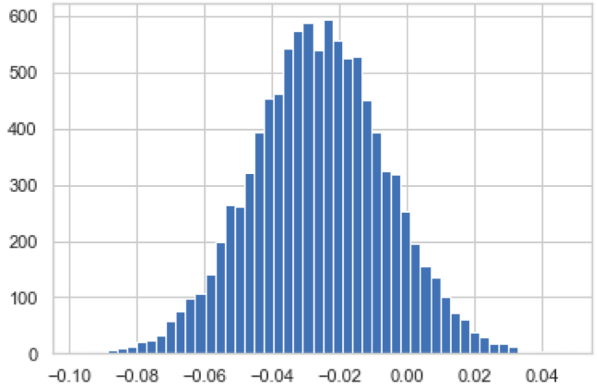


BD_5pr7_WC
P_H0_5pr7_WC 0.0944

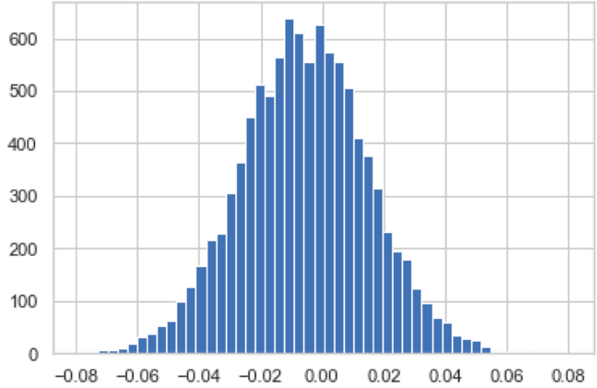


BD_5pr8_iso
P_H0_5pr8_iso 0.3906

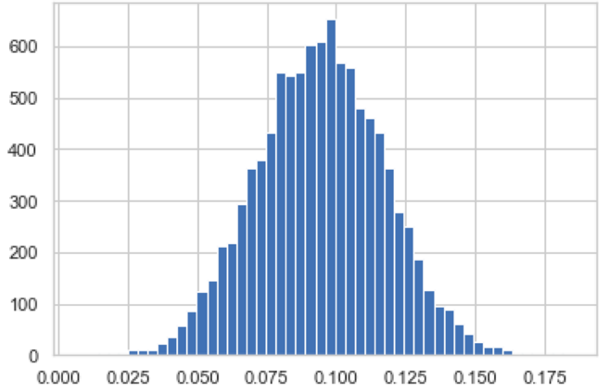


BD_5pr7_iso
P_H0_5pr7_iso 0.0

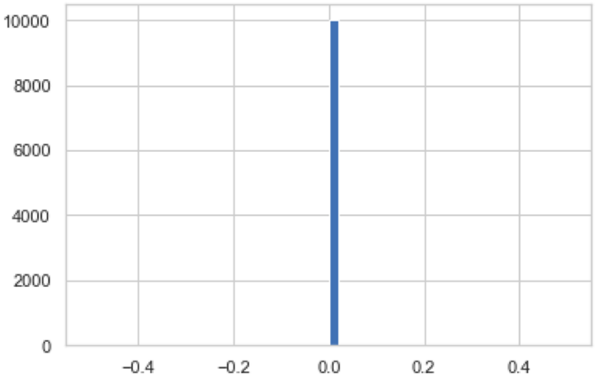


BD_5pr8_dif

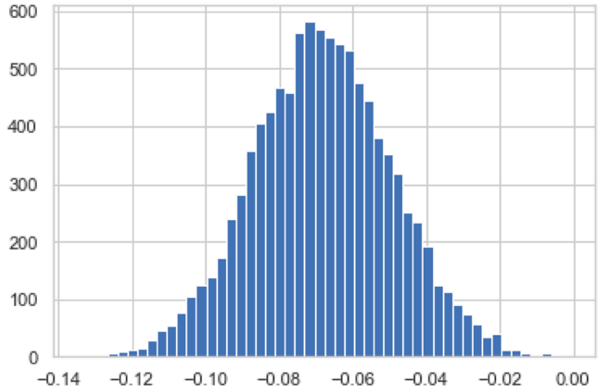


BD_5pr7_dif
P_H0_5pr7_dif 0.0

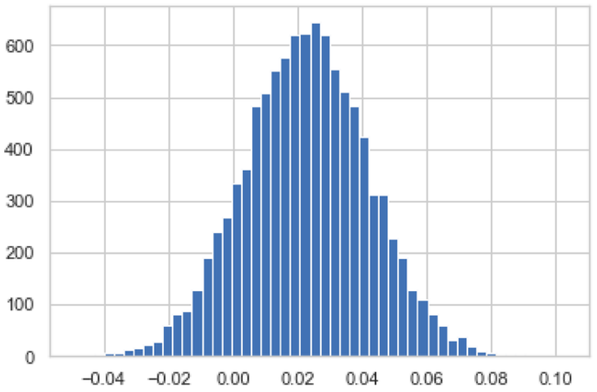


BD_5pr6_WC
P_H0_5pr6_WC 0.1224

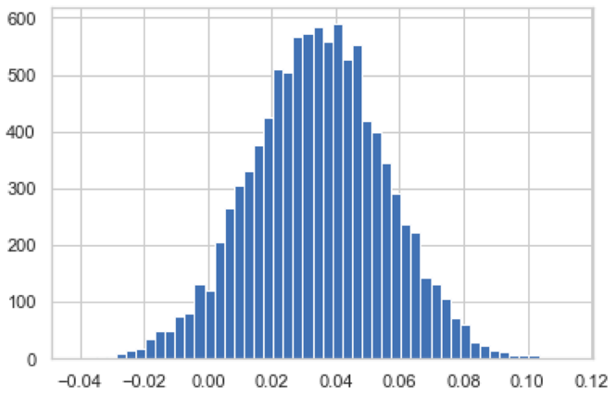


BD_5pr5_WC
P_H0_5pr5_WC 0.0518

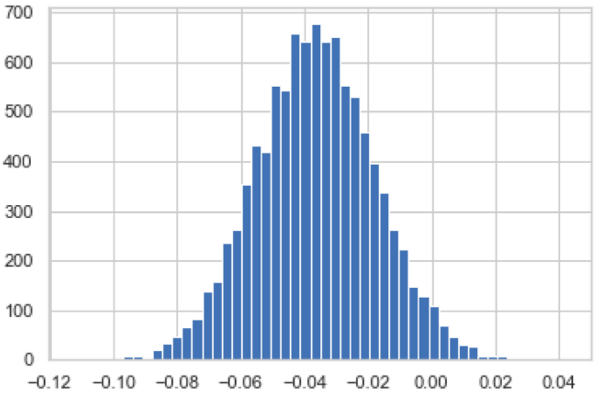


BD_5pr6_iso
P_H0_5pr6_iso 0.0268

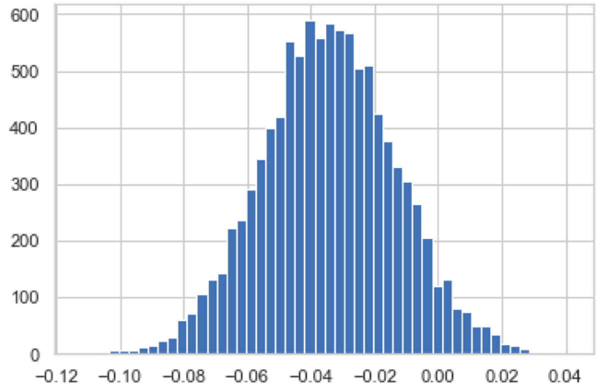


BD_5pr5_iso
P_H0_5pr5_iso 0.0518

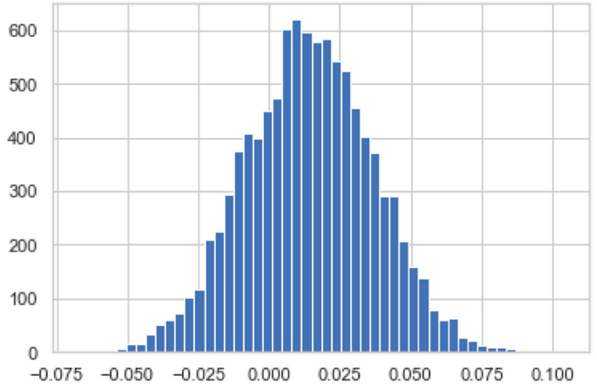


BD_5pr6_dif
P_H0_5pr6_dif 0.2722

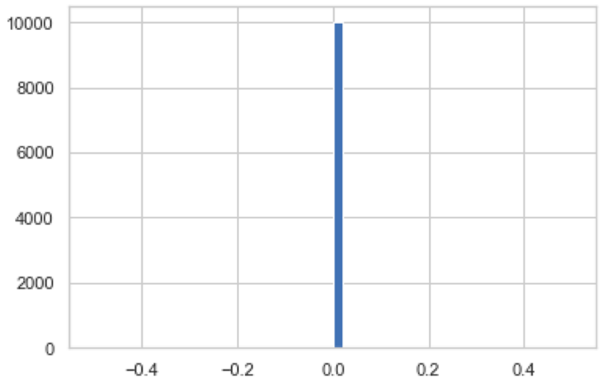


BD_5pr5_dif

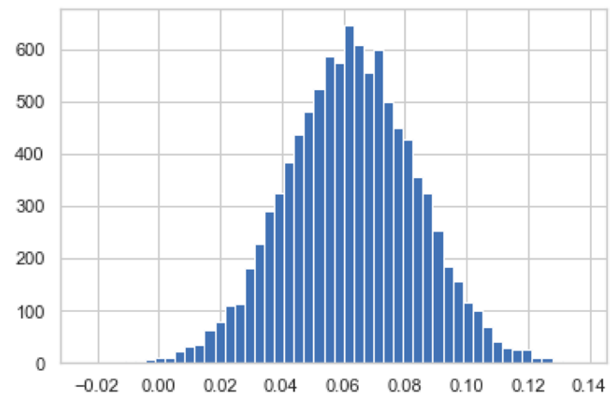


BD_5pr4_WC
P_H0_5pr4_WC 0.002

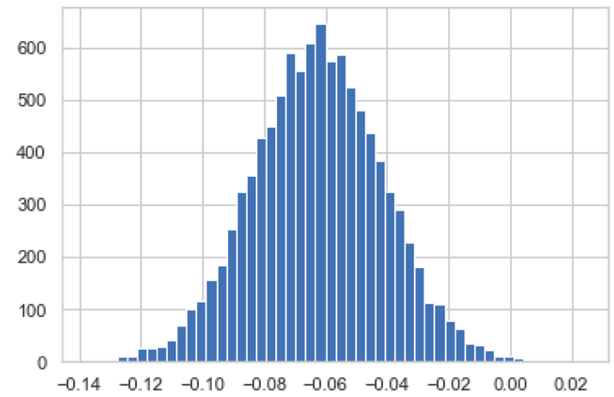


BD_5pr4_iso
P_H0_5pr4_iso 0.002

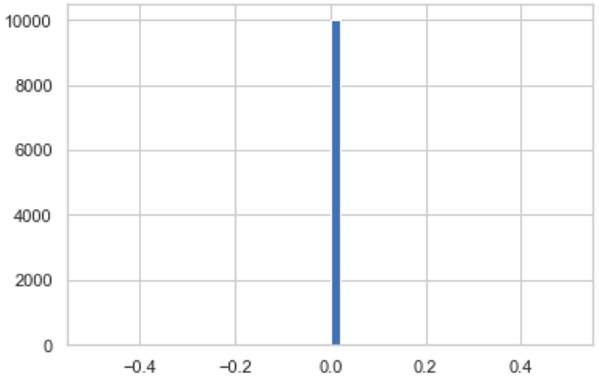


BD_5pr4_dif

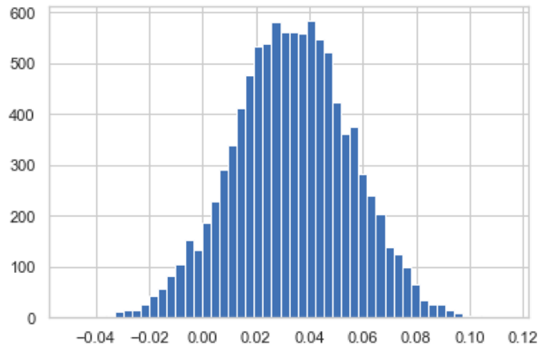


BD_5pr3_WC
P_H0_5pr3_WC 0.0659

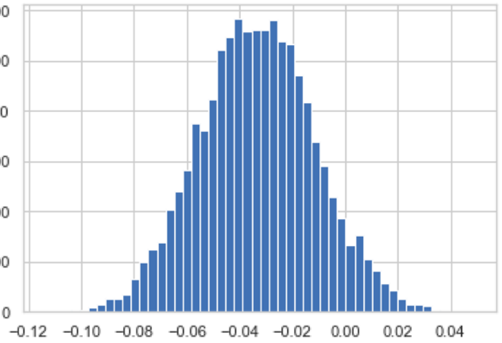


BD_5pr3_iso
P_H0_5pr3_iso 0.0659

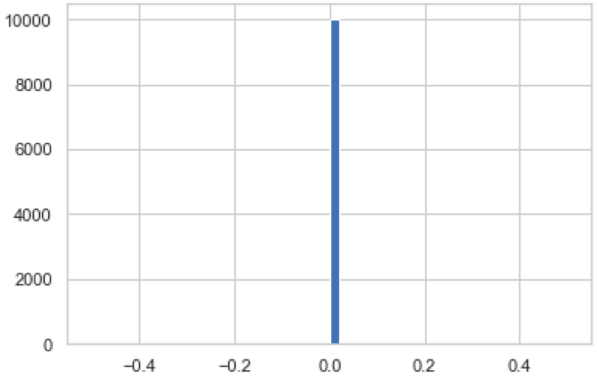


BD_5pr3_dif

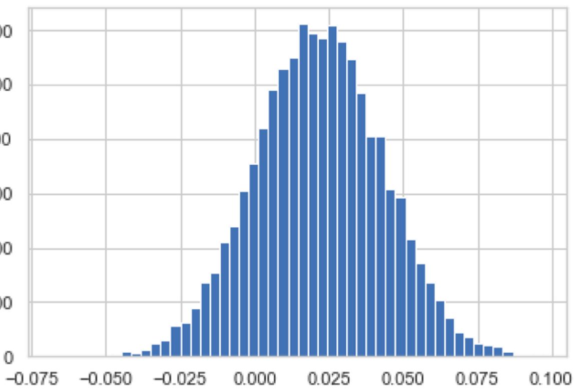


BD_5pr2_WC
P_H0_5pr2_WC 0.1551

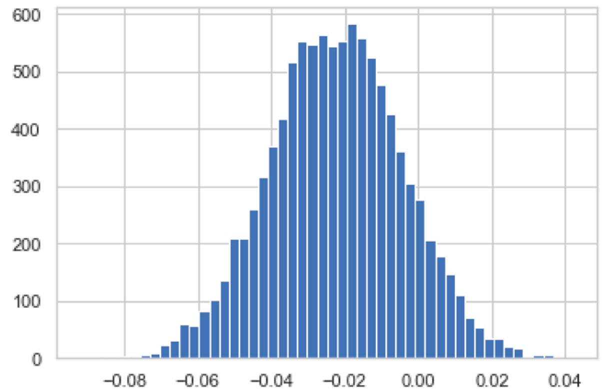


BD_5pr1_WC
P_H0_5pr1_WC 0.1067

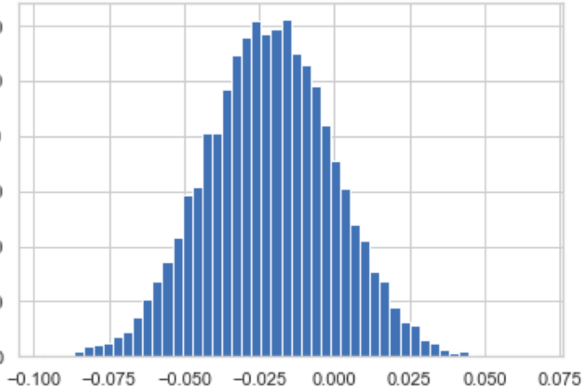


BD_5pr2_iso
P_H0_5pr2_iso 0.1551

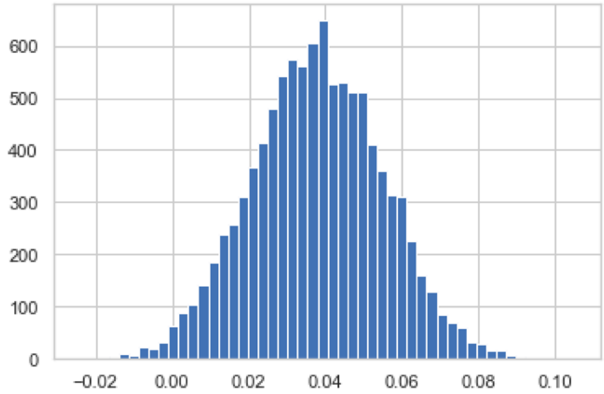


BD_5pr1_iso
P_H0_5pr1_iso 0.0119

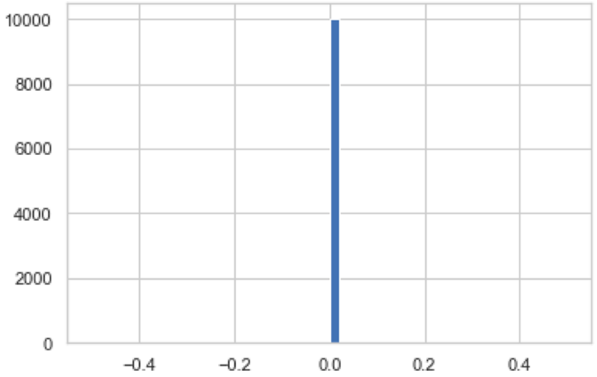


BD_5pr2_dif

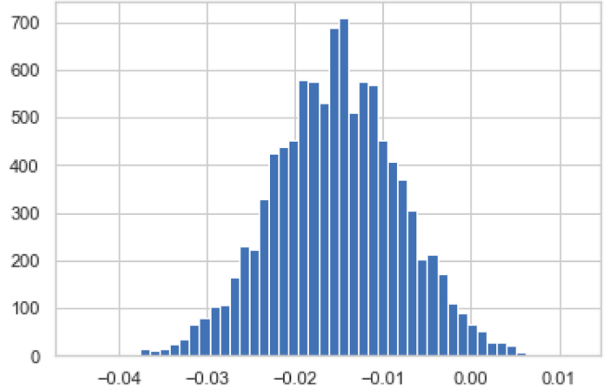


BD_5pr1_dif
P_H0_5pr1_dif 0.0194

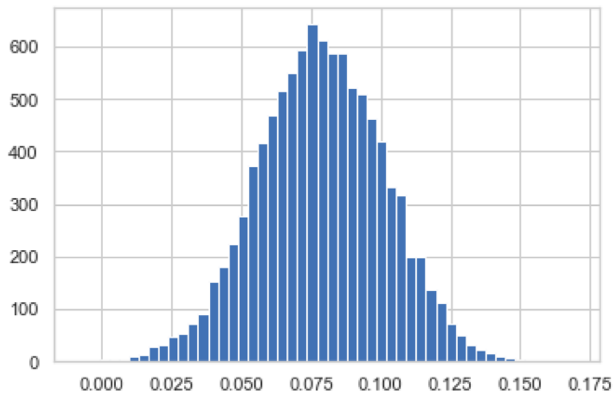


BD_3pr1_WC
P_H0_3pr1_WC 0.0001

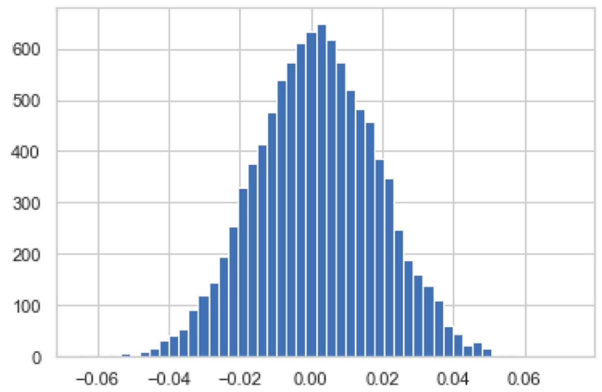


BD_3pr2_WC
P_H0_3pr2_WC 0.464

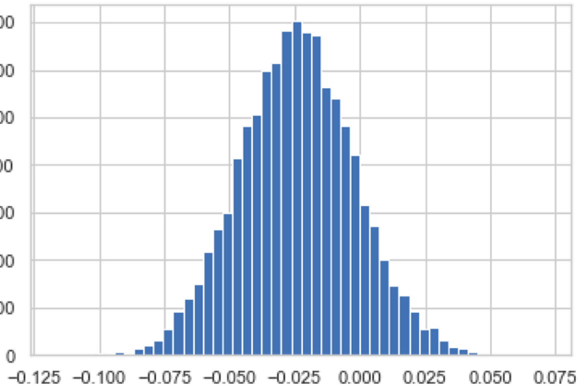


BD_3pr1_iso
P_H0_3pr1_iso 0.1345

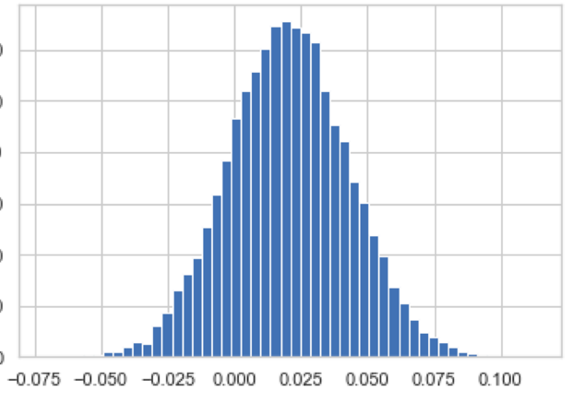


BD_3pr2_iso
P_H0_3pr2_iso 0.1825

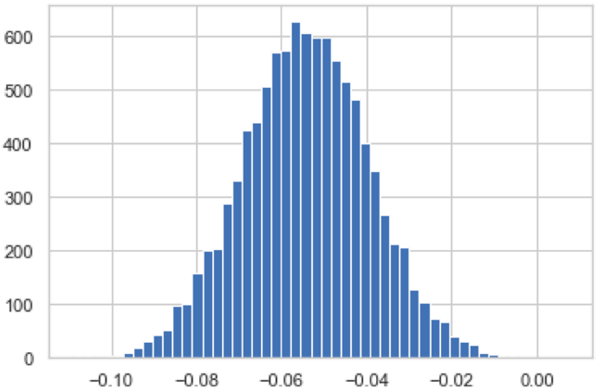


BD_3pr1_dif
P_H0_3pr1_dif 0.0002

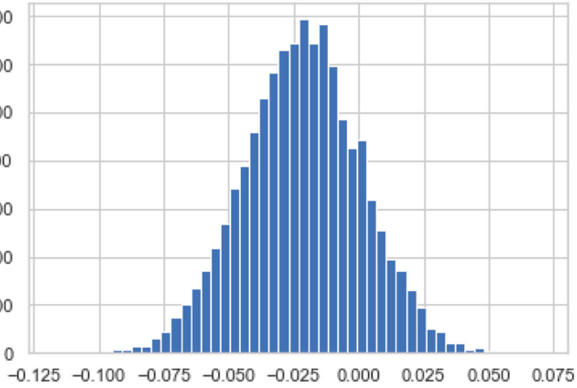


BD_3pr2_dif
P_H0_3pr2_dif 0.1712

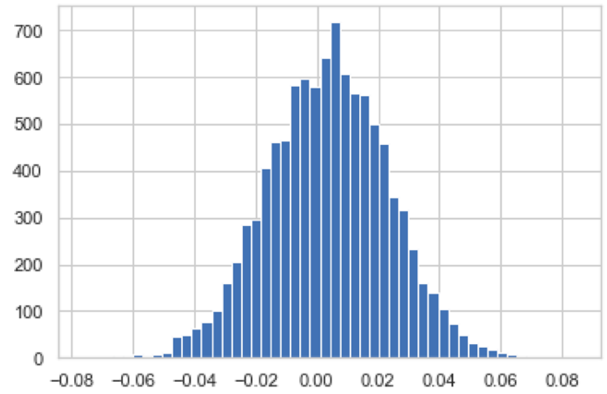


BD_3pr3_WC
P_H0_3pr3_WC 0.4227

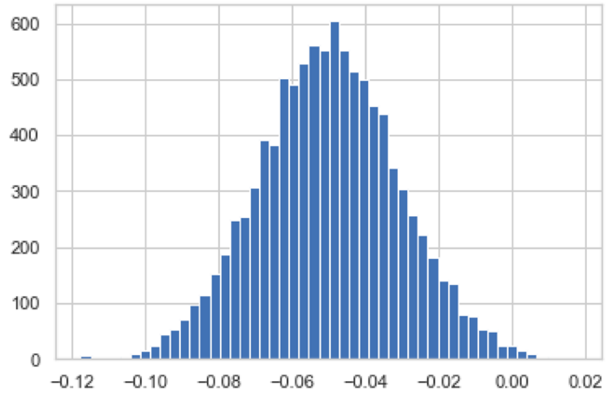


BD_3pr3_dif
P_H0_3pr3_dif 0.0043

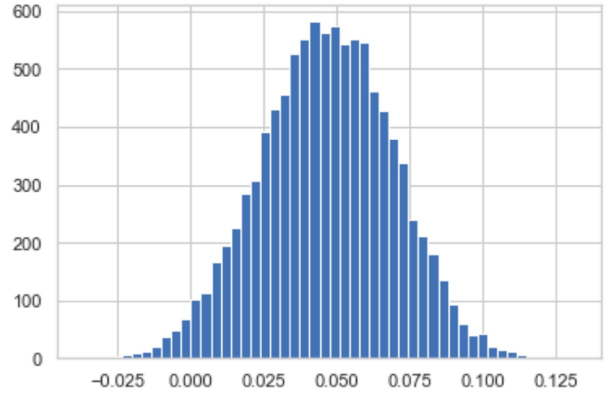


BD_3pr3_iso
P_H0_3pr3_iso 0.0212

**Figure S10|** Histograms of bootstrap difference for U5 bp types at each position of the splice junction between -3Csub and -3C datasets (Violinplots of the same **Figure 9D-F**)

Base pair types: **W**atson-**C**rick, **WC**; **iso**steric, **iso**; and non-isosteric (**dif**ferent), **dif**
Splice junction positions key: **5pr1** is 5'exon position -1; **3pr1** is 3'exon position +1
